# *In vivo* affinity maturation of the HIV-1 Env-binding domain of CD4

**DOI:** 10.1101/2024.02.03.578630

**Authors:** Andi Pan, Charles C. Bailey, Tianling Ou, Jinge Xu, Xin Liu, Baodan Hu, Gogce Crynen, Nickolas Skamangas, Naomi Bronkema, Mai Tran, Huihui Mu, Xia Zhang, Yiming Yin, Michael D. Alpert, Wenhui He, Michael Farzan

**Affiliations:** Skaggs Graduate School, Scripps Research, La Jolla, CA 92037, USA; The Center for Integrated Solutions to Infectious Diseases, The Broad Institute of MIT and Harvard, Cambridge, MA 02142, USA; Division of Infectious Disease, Boston Children’s Hospital, Boston, MA 02115, USA; Department of Pediatrics, Harvard Medical School, Boston, MA 02115, USA; The Herbert Wertheim UF Scripps Institute for Biomedical Innovation & Technology, Jupiter, FL 33458, USA; Emmune Inc., Cambridge, MA 02129, USA

## Abstract

Many human proteins have been repurposed as biologics for clinical use. These proteins have been engineered with *in vitro* techniques that improve affinity for their ligands. However, these approaches do not select against properties that impair efficacy such as protease sensitivity or self-reactivity. Here we engineer the B-cell receptor of primary murine B cells to express a human protein biologic without disrupting their ability to affinity mature. Specifically, CD4 domains 1 and 2 (D1D2) of a half-life enhanced-HIV-1 entry inhibitor CD4-Ig (CD4-Ig-v0) were introduced into the heavy-chain loci of murine B cells, which were then adoptively transferred to wild-type mice. After immunization, transferred B cells proliferated, class switched, affinity matured, and efficiently produced D1D2-presenting antibodies. Somatic hypermutations found in the D1D2-encoding region of engrafted B cells improved binding affinity of CD4-Ig-v0 for the HIV-1 envelope glycoprotein (Env) and the neutralization potency of CD4-Ig-v0 by more than ten-fold across a global panel of HIV-1 isolates, without impairing its pharmacokinetic properties. Thus, affinity maturation of non-antibody protein biologics *in vivo* can guide development of more effective therapeutics.

## INTRODUCTION

Protein biologics play major and increasingly important roles in the treatment of human diseases. These biologics include antibodies, hormones, recombinant cytokines, and immunoadhesins forms of cellular receptors^1^. A number of *in vitro* approaches for improving the affinity of these proteins have been developed, including phage, yeast, and mammalian-cell display techniques, as well as structure-guided design^2–4^. Often these improvements associate with properties that impair the bioavailability of these biologics, for example by increasing interactions with serum or cell-surface proteins or cell membranes^5–11^. In contrast, the natural process of affinity maturation in germinal centers of immunized animals selects for higher affinity while eliminating self-reactive, unstable, poorly expressed, or protease-sensitive antibody variants^12–15^. In addition, *in vitro* selection methods typically proceed in a small number of discrete steps without ongoing diversification of selected intermediates, whereas diversification and selection *in vivo* are continuous, coordinated, and sensitive to small affinity differences^12, 16, 17^. Thus, the affinity maturation in the mammalian germinal centers has critical advantages over *in vitro* techniques for improving the affinity and bioavailability of protein therapeutics.

We have previously shown that human heavy- and light-chain variable regions can directly replace their murine counterparts in primary mature B cells, a technique we describe as ‘native-loci’ editing^18^. When these CRISPR/Cas12a-engineered cells were adoptively transferred to recipient mice, their B-cell receptors (BCRs) affinity matured in response to antigen, facilitating identification of broader, more potent, and more bioavailable forms of the original human antibody. We further observed that somatic hypermutation and affinity maturation of the engineered BCR were markedly more efficient when exogenous variable chain genes were introduced at their native-loci rather than in a commonly used intronic region, indicating that the latter insertion disrupted the activities of the engineered cell. The greater versatility and rapid turnaround time of native-loci editing of mature B cells suggested that this approach could also supplant transgenic mice as a tool for optimizing the efficacy of therapeutic antibodies *in vivo*.

However, many important protein therapeutics are not antibodies. Rather, they were developed as soluble forms of cellular receptors and recombinant forms of cytokines and serum proteins, such as hirudin^19^, IL-27^20^, factor IX^21^, CTLA4^22, 23^, tumor necrosis factor^24, 25^, and the HIV-1 receptor CD4^26, 27^. To determine if *in vivo* affinity maturation could serve as a useful tool for improving the potency of non-antibody biologics, we selected the domains 1 and 2 (D1D2) of human CD4 for proof-of-principle studies. CD4-Ig, the immunoadhesin form of CD4 D1D2, had been evaluated as a potential therapy for HIV-1 infection, but its limited half-life precluded its clinical use^26, 28, 29^. Here we sought to improve a recently developed CD4-Ig variant, CD4-Ig-v0, a product of iterative *in vitro* optimization for half-life, breadth, and potency^30, 31^. We show that when a sequence encoding D1D2 fused to an antibody heavy chain variable region was introduced into the VDJ-recombined locus of primary murine B cells, these cells affinity matured in recipient mice immunized with mRNA lipid nanoparticles (LNP) that expressed the HIV-1 envelope glycoprotein. High frequency somatic hypermutations observed in multiple mice markedly enhanced the neutralization potency of CD4-Ig-v0 to below 1 μg/ml while retaining its near absolute breadth, high thermostability and long *in vivo* half-life. Thus, non-antibody protein therapeutics can be expressed from the BCR locus and affinity matured in mouse germinal centers.

## RESULTS

### Engineering primary mouse B cells to express a protein biologic *ex vivo*

We have shown that introduction of human antibody heavy chain complementarity determining regions (HCDR3s) or heavy- and light-chain variable genes into their respective murine loci enabled *in vivo* affinity maturation of the resulting chimeric BCRs^18, 32^. Extension of this approach to immunoadhesins such as CD4-Ig is complicated by the absence of the first heavy constant region 1 (CH1) that noncovalently associates with a light-chain constant domain. We therefore evaluated three designs of BCR capable of presenting a half-life and potency optimized form of human CD4 domains 1 and 2^30^ (D1D2; residues 1-173 of the mature CD4 protein) in primary murine B cells while ensuring that CH1 associates with a Cκ domain (**Extended Data** Fig. 1a-b). Among these designs, the highest expression of D1D2 was observed when it was fused to the amino-terminus of the heavy-chain variable domain of the murine antibody OKT3 (OKT3-V_H_) through a (G_4_S)_3_ linker (**Fig. 1a; Extended Data** Fig. 1c). In this design, OKT3 VH and CH1 domains associate with an endogenous light chain expressed in these cells. We therefore sought to replace the VDJ-recombined heavy-chain variable (VH) gene of naïve mature murine B cells with a sequence encoding this construct, D1D2-OKT3-V_H_, while utilizing a native murine variable-segment promoter and a native heavy-chain signal sequence (**Fig. 1b**). Accordingly, we introduced a double-stranded break by CRISPR/Mb2Cas12a ribonucleoproteins (RNPs) into the 3’-most JH segment, JH4 in mice^18, 32, 33^. Repair of this break was guided by a homology-directed repair template (HDRT) delivered by recombinant adeno-associated virus DJ (AAV-DJ). This HDRT included homology arms complementary to a 570 bp 5’ untranslated region (UTR) upstream of VH1-34 or VH1-64, and a 600 bp 3’ intronic region immediately downstream of the J4 segment. An insert sequence encoding D1D2-OKT3-V_H_ and terminating with a modified heavy-chain splice donor was located between these homology arms. The result is that the VH, DH, and JH segments between the targeted VH (VH1-34 or VH1-64) and JH4 segment are replaced with this chimeric heavy-chain gene and the expressed protein resembles a mouse BCR except that it initiates with D1D2.

**Fig. 1.**
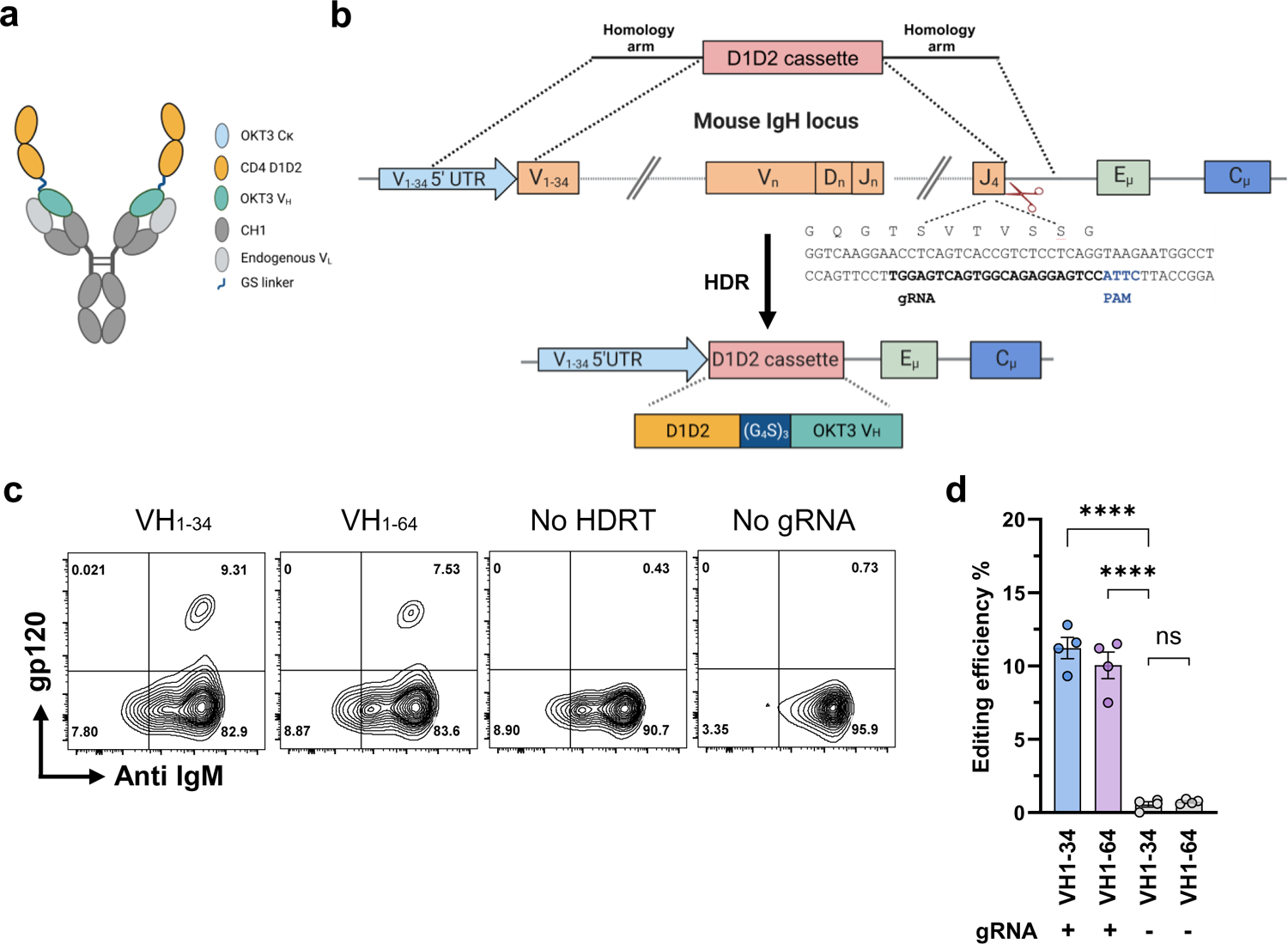
Engineering primary murine B cells to express a B-cell receptor with CD4 domains 1 and 2. **a** A representation of an engineered BCR with a potency and half-life enhanced form of CD4 domains 1 and 2 (D1D2) fused through a (G_4_S)_3_ linker to the amino-terminus of the heavy-chain variable region of the mouse antibody OKT3 (D1D2-OKT3-V_H_). The OKT3 heavy chain pairs with an endogenous mouse light chain. **b** Introducing D1D2-OKT3-V_H_ at the murine heavy-chain locus. The CRISPR effector protein Mb2Cas12a targets the J4 coding region 5’ of a CTTA PAM, as represented. An rAAV-delivered homology directed repair template (HDRT) complements the 5’ UTR of a VH segment and the intron 3’ of JH4 using 576 bp and 600 bp homology arms, respectively. The edited genome replaces the VDJ-recombined heavy chain with a cassette encoding D1D2-OKT3-V_H_. **c** Expression of D1D2-OKT3-V_H_ in primary mouse B cells. Expression of D1D2 in edited cells was measured by flow cytometry with monomeric HIV-1 gp120. Representative flow cytometry plots of B cells edited with HDRT targeting the 5’ UTR of VH1-34 or V1-64 were generated 48 h after electroporation. HDRT were delivered with rAAV transduced at 10^4^ multiplicity of infection (MOI). Controls include cells electroporated with Mb2Cas12a ribonucleoproteins (RNP) without rAAV (No HDRT) or without gRNA but transduced with HDRT-encoding rAAV (No gRNA). Plots were gated on viable singlet B cells. **d** Quantitation of editing efficiency in **c** from independent experiments. Each dot represents an average from two biologically independent replicates. Error bars represent standard error of mean (SEM). Statistical significance was determined by two-way ANOVA followed by H-Šídák’s multiple comparisons (****p < 0.0001). **a**, **b** are created with BioRender.com.

We first compared the efficiency of HDRTs with 5’ homology arms targeting either VH1-34 or VH1-64. VH1-64 is upstream of VH1-34 in the heavy-chain locus and more commonly used in C57BL/6 mouse splenic IgM+ B cells^34, 35^. We observed that HDRT targeting VH1-34 resulted in an average of 11% editing efficiency, modestly greater than observed with AAV targeting VH1-64, as indicated by flow cytometry using a fluorescently labeled anti-CD4 antibody or HIV-1 envelope glycoprotein gp120 (**Fig. 1c and d**). This efficiency is higher than necessary for expansion and affinity maturation of engineered B cells ^16, 18, 32^. Thus, primary murine B cells can be engineered *ex vivo* to efficiently express D1D2-OKT3-V_H_.

### Engineered B cells generated neutralizing sera in immunized mice

To evaluate the *in vivo* activity of engineered cells, we harvested splenic B cells from B6 CD45.1 mice, replaced their heavy-chain variable genes with D1D2-OKT3-V_H_ and adoptively transferred these cells to wild-type (CD45.2) C57BL/6J mice. Because our observed editing efficiency was higher than necessary for expansion and affinity maturation of engineered B cells^16, 32^, HDRT levels were adjusted so that roughly 15,000 B cells (0.3%) expressing D1D2-OKT3-V_H_ were engrafted. Mice were then immunized intramuscularly 24 hours post-engraftment and boosted twice at two- or four-week intervals (2 wk, 4 wk) with mRNA lipid nanoparticles (mRNA-LNP) encoding an engineered HIV-1 envelope glycoprotein (Env) trimer, namely the previously described 16055-ConMv8.1 SOSIP-TM^32^ (**Fig. 2a**). The neutralization potency of sera was monitored after each immunization using pseudoviruses expressing the BG505 or CE1176 Envs (**Fig. 2b-d; Extended Data** Fig. 2a). No neutralization was observed using sera from mice engrafted with edited cells one week after the first vaccination, or from mice engrafted with unedited cells throughout the study. Lack of neutralization after priming is consistent with the low number of successfully edited cells^36^. After two immunizations, sera from LNP-vaccinated mice detectably neutralized the heterologous BG505 pseudovirus with average 50% inhibitory dilutions (ID_50_) ranging from 20 to 45. Note that CD4-Ig-v0 neutralizes comparably but less efficiently than an antibody with a D1D2-OKT3-V_H_ variable chain (**Extended Data** Fig. 2b). Similar results were observed with an additional group of five mice immunized with mRNA-LNP at four-week intervals (**Fig. 2c-d**). Enzyme-linked immunosorbent assay (ELISA) measurements of sera from immunized mice indicate that, after two or three LNP immunizations, average D1D2-OKT3-V_H_-IgG concentrations ranged from 40 to 75 μg/ml (**Fig. 2e**). A parallel study with adjuvanted gp120-based protein antigens was also performed. Specifically, mice were engrafted with 5 million cells, including either 1.5×10^4^ or 5×10^5^ D1D2-OKT3-V_H_-expressing cells, followed by immunization at two-week interval with multimeric (F10) and monomeric gp120 constructs (**Extended Data** Fig. 2c). Notably, in contrast to mice vaccinated with mRNA-LNP, no neutralizing activity was observed in mice receiving 1.5×10^4^ successfully edited cells and vaccinated with these protein antigens (**Extended Data** Fig. 2d). However, robust responses were observed in mice infused with 5×10^5^ successfully edited cells. Thus, as previously reported, adoptive transfer of more antigen-reactive B cells can compensate for a less immunogenic vaccine^16, 36^. We conclude that immunization of mRNA-LNP-expressing SOSIP-TM antigens can generate therapeutic concentrations^27, 37, 38^ of D1D2 in mice engrafted with a limited number of D1D2-OKT3-V_H_-edited B cells.

**Fig. 2.**
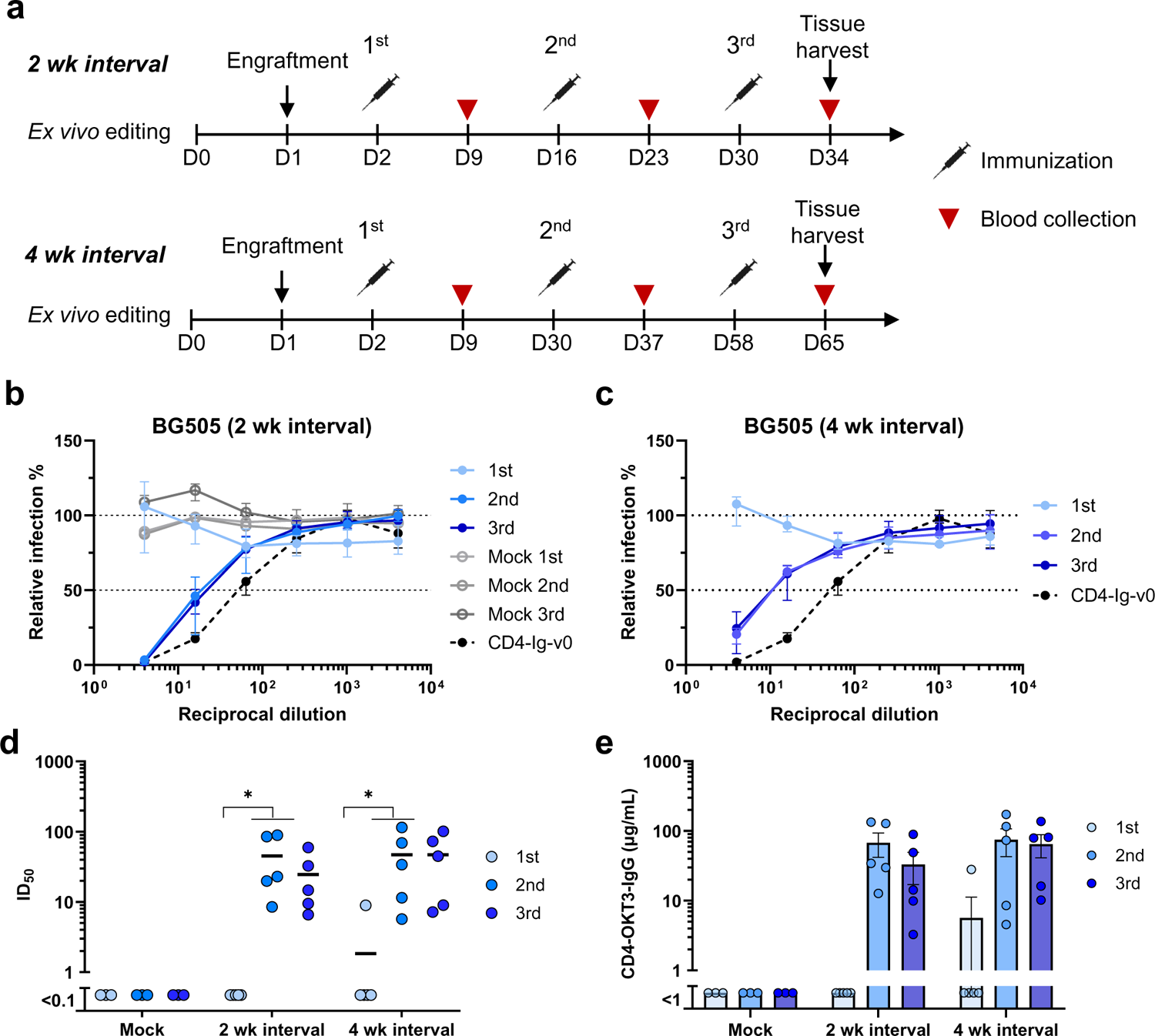
Engineered B cells generated neutralizing responses in immunized mice. **a** Schedule of immunization and blood collections from mice analyzed in subsequent figures. Naïve B cells from CD45.1 donor mice were engineered *ex vivo* and 5 million cells per mouse were adoptively transferred to CD45.2 recipient mice 24 h later. Mice were immunized with SOSIP-TM (16055-ConM-8.1) mRNA-LNP on two-week (2 wk, Day 2, 16, and 30, n = 5) or four-week (4 wk, Day 2, 30, 58, n = 5) intervals, and serum was collected seven days after each immunization. Spleens and lymph nodes were harvested four days after the final immunization and B cells isolated from these tissues were analyzed by flow cytometry and next-generation sequencing (NGS). **b, c** Neutralizing responses of sera from mice immunized with SOSIP-TM mRNA. Sera from mice immunized at two-week (**b**) or four-week (**c**) intervals were measured individually for their ability to neutralize BG505 HIV-1 pseudovirus in TZM-bl cell assays. Sera from mice (n = 3) engrafted with unedited CD45.1 B cells and immunized on two-week interval (grey dots) served as negative controls. 100 μg/ml CD4-Ig-v0 combined with normal mouse serum served as a positive control. Dots and error bars indicate median and interquartile range for each group. **d** A summary of the 50% inhibitory dilutions (ID_50_) of sera from each immunized mouse in **b** and **c**. Statistical significance was determined using repeated-measures two-way ANOVA with Geisser-Greenhouse correction. **E** Serum concentration of CD4-OKT3-IgG after each immunization, measured by ELISA with an anti-CD4 antibody and CD4-OKT3-IgG as the standard.

### Antigen-induced activation of engineered B cells *in vivo*

We further analyzed B cells harvested from the spleens and lymph nodes of mRNA-LNP vaccinated mice engrafted with cells expressing D1D2-OKT3-V_H_ or with unmodified B cells. Similar percentages of CD45.1+ donor cells were found in mice vaccinated at two- and four-week intervals, and percentages were modestly but not significantly higher than mice engrafted with unedited cells (**Fig. 3a; Extended Data** Fig. 3a-b). Higher percentages of antigen-reactive cells were found in the CD45.1+ donor-cell population than in the CD45.2+ host-cell population (**Fig. 3b**). Fewer gp120-reactive CD45.2+ B cells were observed in mice engrafted with D1D2-expressing cells than in mock engrafted mice, suggesting competition between edited donor cells and host B cells. The properties of GC B cells (CD38-,GL7+) from four representative mice (2 wk: M1, M2; 4wk: M7, M9) were further characterized (**Extended Data** Fig. 3c). Roughly 60% of GC CD45.1+ donor B cells bound gp120, whereas less than 10% of CD45.2+ recipient-mouse cells (**Fig. 3c-d; Extended Data** Fig. 3d). Nearly every antigen-reactive BCR from CD45.1+ donor cells had class switched from IgM to an IgG isotype, predominantly IgG1 (**Fig. 3e**). We conclude that gp120-reactive donor B cells class-switched and migrated to the spleen and lymph node germinal centers.

**Fig. 3.**
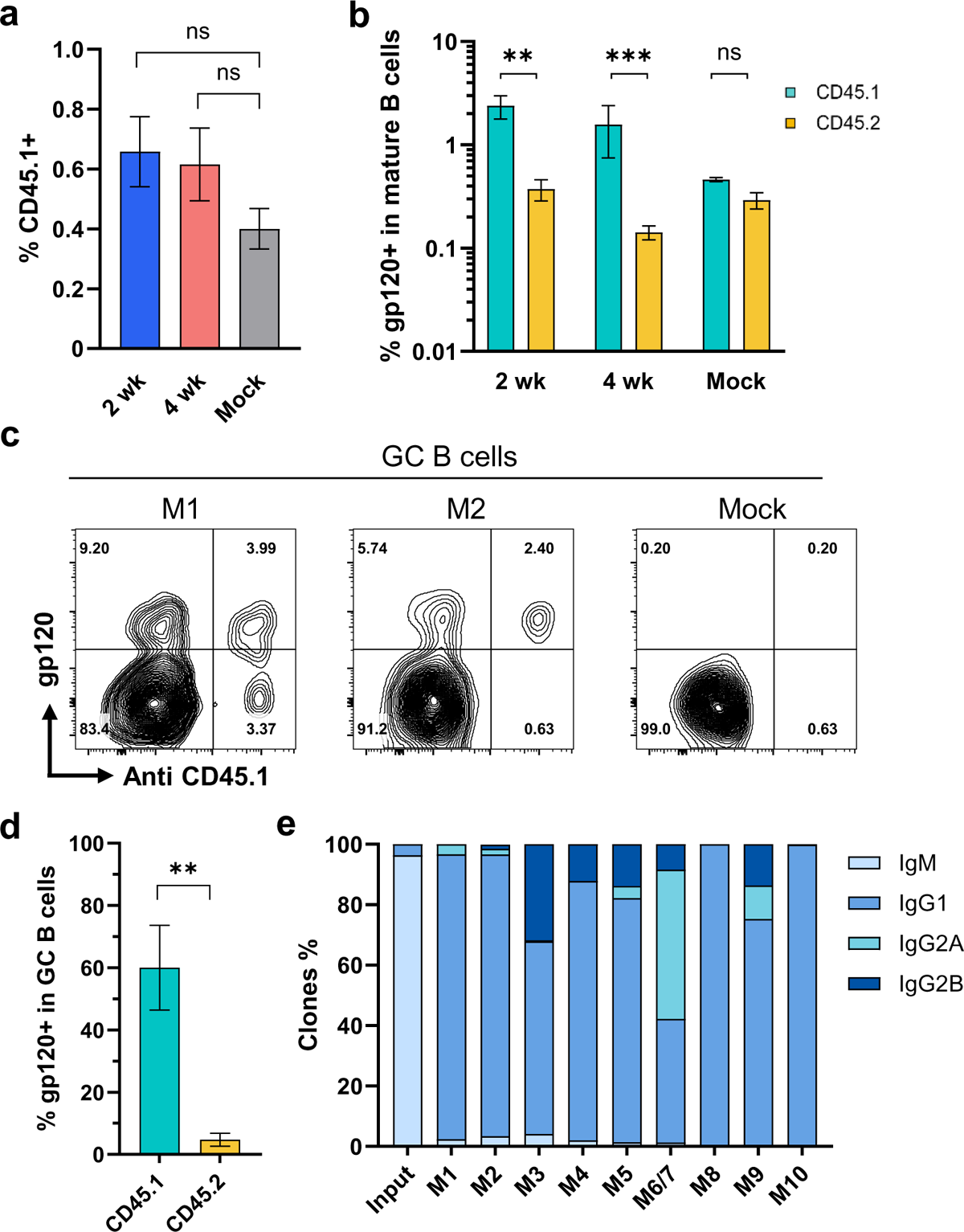
Engineered B cells persisted *in vivo* following immunization. **a** Quantification of CD45.1-positive donor B cells in vaccinated mice. Four days after the final vaccination of mice characterized in Fig. 2, B cells were isolated from their lymph nodes and spleens. B cells were analyzed by flow cytometry. Figure shows the percent of CD45.1-positive donor cells in mice immunized at two-week (2 wk, n = 4) or four-week intervals (4 wk, n = 5). Mice similarly engrafted with unedited cells and immunized in parallel (Mock, n = 3) served as the controls. For gating strategies, see **Extended Data** Fig. 3a. **b** A greater proportion of CD45.1 donor cells binding to HIV-1 gp120. The cells analyzed in panel **a** were measured for binding to HIV-1 gp120. See **Extended Data** Fig. 3b for source flow cytometry analysis. **c** Enrichment of gp120-binding donor cells in the germinal center. Germinal center (GC) B cells (CD38-GL7+) were analyzed by flow cytometry for their ability to bind gp120 and an anti-CD45.1 antibody. Two representative examples (M1 and M2) from mice immunized at two-week interval are shown. Additional examples are provided in Extended Data Fig. 3d. **d** Quantification of results from experiments shown in **c** (n = 4). **e** Distribution of isotypes among D1D2-expressing donor B cells isolated after the final immunization compared with edited donor B cells before engraftment, as determined by NGS. M6 and M7 was combined as one sample. For **a**, **b**, **d,** error bars indicate SEM. Statistical significance was determined by generalized linear mixed model followed by Tukey HSD pairwise comparisons (*p < 0.05; **p < 0.01; ***p < 0.001).

### Somatic hypermutation of the D1D2 region in engineered B cells *in vivo*

We next investigated whether D1D2-expressing gp120-reactive B cells underwent somatic hypermutation (SHM). Accordingly, 3,000 to 7,000 IgG+ CD45.1+ gp120-reactive B cells were isolated and the D1D2 region was sequenced. Substantial SHM was observed in all ten mRNA-LNP-immunized mice (**Fig. 4a**). Despite the heterogeneity of these responses, several mutations were observed at high frequency across multiple mice. Most notably, two mutations in codons for R59 and K90 in CD4 domain 1 were uniformly present among the 10 mice, with average frequencies of 54% and 64% and ranges of 5 to 89% and 30 to 81%, respectively. The predominance of these mutations is more remarkable given the average mutation frequency across all 516 nucleotides was 1.1%. Several non-coding changes were also observed at high average frequency. The seven most frequent non-coding changes coincided with defined AID hotspots (DGYD and WRCH with letters corresponding to the IUPAC nucleotide code), whereas the top three coding mutations emerged away from these hotspot sequences (**Extended Data** Fig. 4), suggesting that these latter changes were products of strong selection pressure. Consistent with ongoing purifying selection, non-coding changes were observed more frequently than coding changes, but several coding changes, again including those at R59 and K90, dominated once they emerged (**Fig. 4b**). The average number of total mutations was the same between mice immunized at two- and four-week intervals (**Fig. 4c**). However, more non-coding changes were found in mice immunized every four weeks, in particular the region encoding CD4 domain 2 (**Fig. 4d**). Non-coding changes distributed evenly across CD4 domains 1 and 2, and thus proximity to the VH1-34 promotor did not affect SHM rates. Most D1D2-encoding sequences included at least two nucleotide changes, and roughly half had three or more (**Fig. 4e**). The frequency of unmodified D1D2 sequences varied from 0% to 20%, with fewer unmutated clones in the mice immunized at four-week intervals than in the two-week group. We conclude that B cells edited to express D1D2-OKT3-V_H_ underwent substantial SHM.

**Fig. 4.**
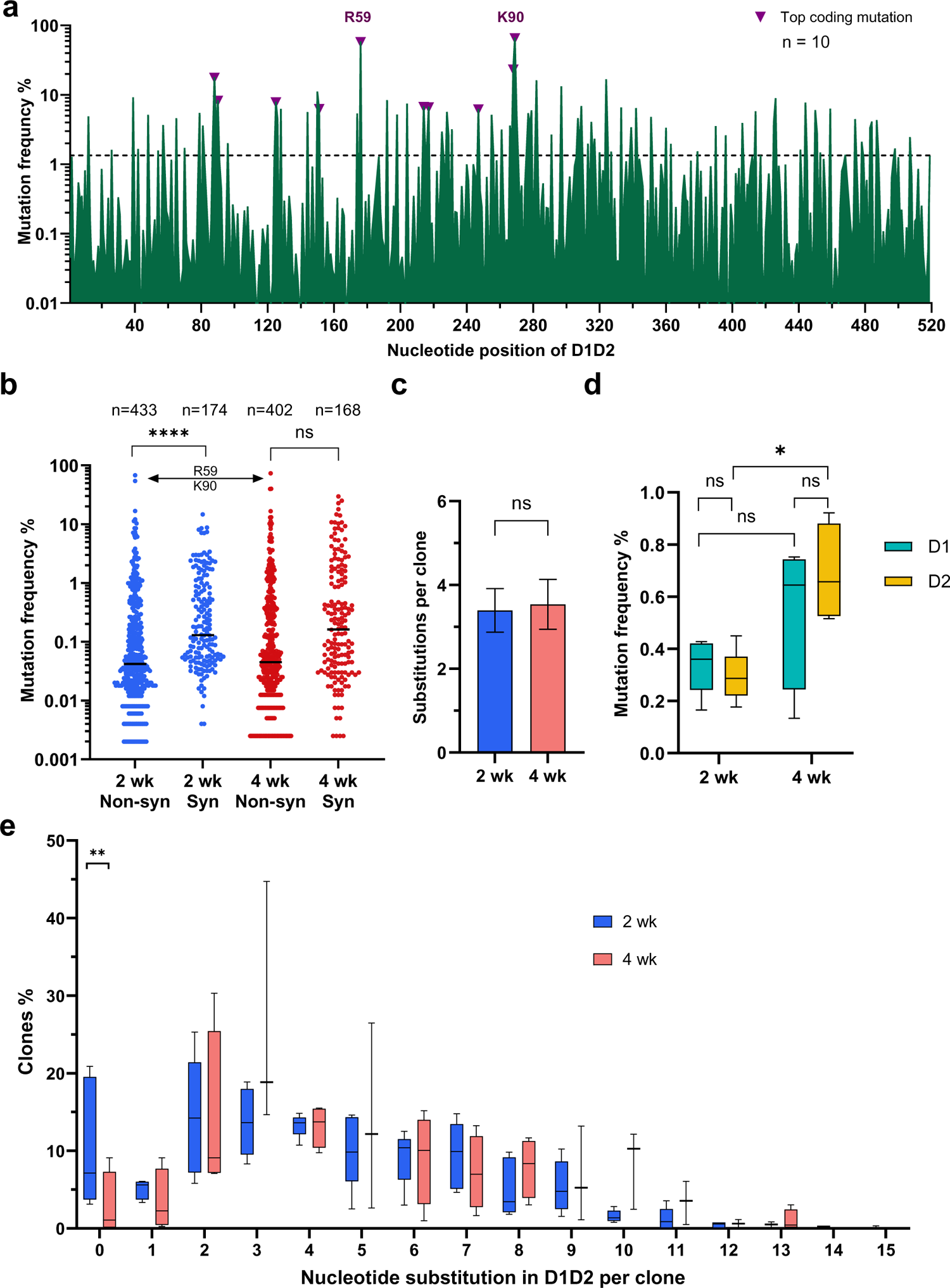
D1D2-expressing B cells hypermutated and class switched *in vivo.* **a** Nucleotide mutation frequency across the D1D2-encoding region. mRNA isolated from CD45+ IgG+ gp120-binding B cells from each of 10 mice was analyzed by NGS and the mean frequency of nucleotide mutations is plotted for each position. Triangles represent the most frequent coding mutations. Codons for R59 and K90 are indicated. **b** Distribution of synonymous (Syn) and non-synonymous (Non-syn) mutation frequency across D1D2. Each dot represents the mutation frequency at one nucleotide position, averaged on five mice in each group. The top two dots indicate the nucleotide mutations leading to R59 and K90 mutations. Line indicates the median. **c** Average number of accumulated mutations per unique sequence. Significance was determined by two-tailed unpair t test. **d** The frequency of synonymous mutations within domains 1 (D1) and 2 (D2) for mice immunized at two-week (2 wk) and four-week (4 wk) intervals. The center line indicates mean, and boxes denote quartile range. Repeated measure mixed effects analysis with H-Šídák’s multiple comparisons (*p < 0.05). **e** Distribution of accumulated nucleotide mutations per unique D1D2 sequence. The center line indicates mean, and boxes denote quartile range. Statistical significance in **b**, **e** was determined by mixed effects analysis with H-Šídák’s multiple comparisons (*p < 0.05; **p < 0.01; ***p < 0.001; ****p < 0.0001).

### High-frequency amino-acid changes recurred in engrafted mice

Further analysis of coding mutations observed in mRNA-LNP-immunized mice showed mouse-to-mouse variation, but also a number of consistencies (**Fig. 5a**). For example, in addition to R59 and K90, most mice encoded changes of N30 (**Fig. 5b**; **Extended Data** Fig. 5a). These N30 mutations, observed in 6 of 10 mice, overlay an AID hotspot motif, whereas R59 and K90 changes, found in 8 and 9 mice, respectively, were not close to any defined hotspots. These changes were also dominant in protein-immunized mice (M16 – M24, **Extended Data 5b**). A minimum spanning tree clustering analysis highlights the underlying diversity of sequences found in each LNP-immunized mouse, and the presence of multiple successful founder sequences that give rise to multiple closely related sequences (**Fig. 5c**; **Extended Data** Fig. 6a-b). High-frequency mutations, including again R59 and K90, were consistently observed among the largest clusters in most mice. Of note, trees generated from mice engrafted with 500,000 D1D2-expressing cells and immunized with adjuvanted protein antigens were sparser and contained more unmutated ancestral sequences than mRNA-immunized mice engrafted with far fewer D1D2-expressing cells, suggesting that somatic hypermutation was less robust with protein antigens (**Extended Data** Fig. 6b). Collectively, these data show that the D1D2 region introduced at heavy-chain locus underwent robust and heterogenous SHM *in vivo*, and that a small subset of these hypermutations were strongly favored in most mice.

**Fig. 5.**
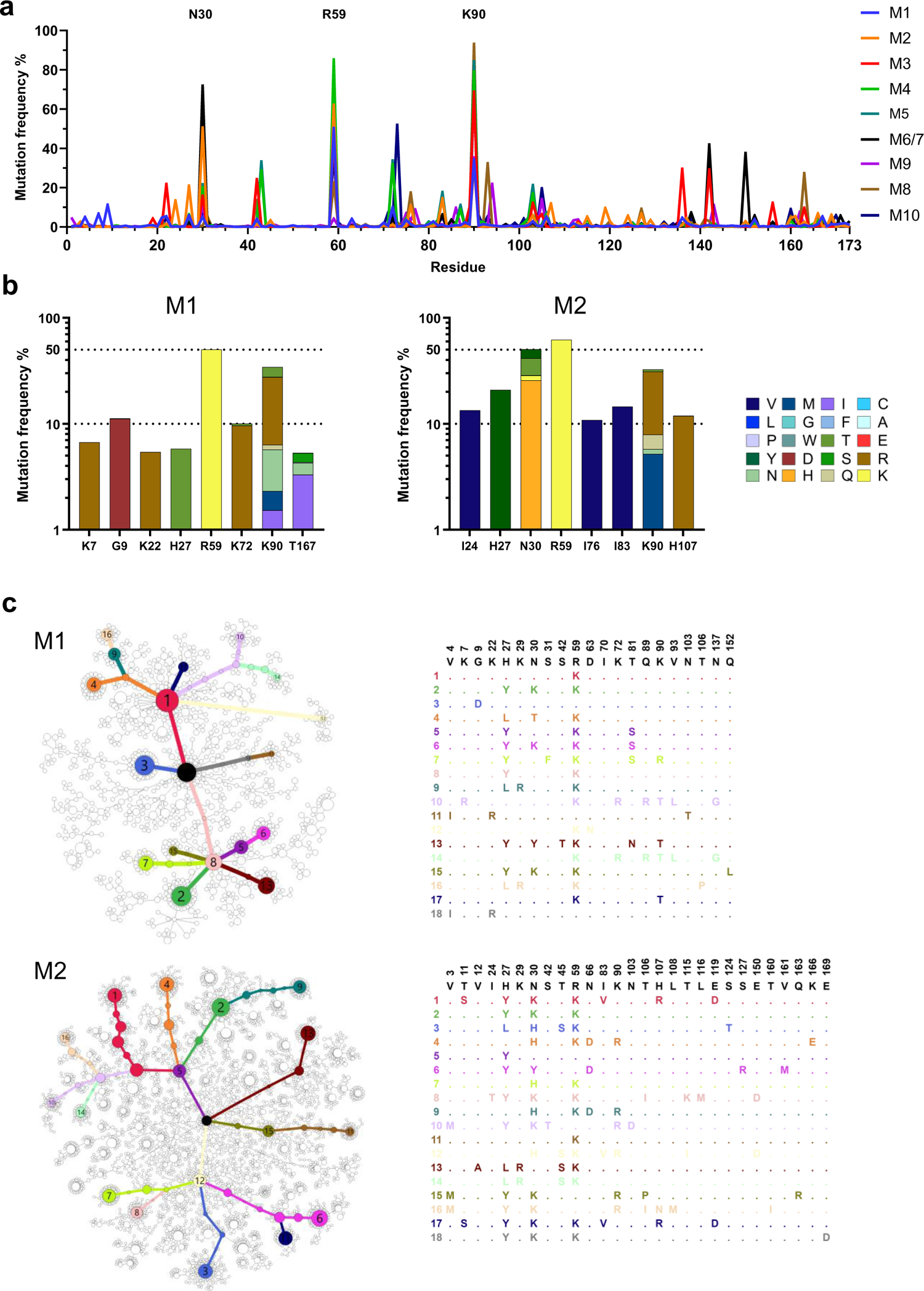
Diverse and convergent amino-acid mutations in engrafted mice. **a** The frequency of amino acid changes across D1D2 sequences from each mouse (M1 through M10). Three amino acids with the highest mutation rate are labeled. M6 and M7 were combined in sequencing analysis. **b** Amino acid changes found in the eight most frequently mutated D1D2 residues from the indicated mice immunized at two-week intervals (M1, M2) are represented. The data for remaining mice are shown in Extended Data Fig. 5. **c** Minimum spanning trees of D1D2 sequences from individual mice. Each tree presents the inferred lineage and all amino-acid mutations found in M1 and M2. The central black dot represents the inferred ancestral sequence which corresponds to the input sequence. Each circle indicates a distinct amino-acid sequence. Circle size is proportional to the number distinct nucleotide sequences with the same translation. Colored circles mark translations encoded by the eighteen largest number of distinct sequences, with the rank order indicated by number. Branch length corresponds to evolutionary distance, defined as the number of amino-acid differences. The figures for the remaining mice are provided in Extended Data Fig. 6.

### *In vivo* hypermutations markedly improve the potency of the CD4-Ig-v0

Because R59K and K90R mutations were prominent in most immunized mice, we compared CD4-Ig variants with these changes to CD4-Ig-v0 for its neutralization potency in TZM-bl cells. (**Fig. 6a**; **Extended Data** Fig. 7a). R59K improved the potency of CD4-Ig-v0 against all nine isolates tested, whereas K90R improved the potency against eight. The combination of these two mutations improved the potency against all isolates, with a geometric mean potency 13-fold greater than CD4-Ig-v0. Parallel analysis of N30H indicated that this mutation did not improve neutralization against most assayed isolates (**Extended Data** Fig. 7b-c), and therefore it was excluded from further analysis. We also characterized a number of less prominent mutations against a CD4-sensitive (BG505) and a CD4-resistant (TRO11) isolate (**Extended Data** Fig. 7c). Among these, only D63N improved neutralization against both isolates, consistent with its presence at the interface between gp120 and CD4. We also included for analysis three mutations away from the gp120-binding interface (**Fig. 6b)**. We speculated that these changes might enhance the *in vivo* stability of CD4-Ig-v0, because they either emerged frequently in multiple mice (H27Y, T160S) or appeared to fill a hydrophobic cavity present in CD4 domain 2 (I138F). Four combinations of these mutations with R59K and K90R were generated (v1-v4) and characterized against a global panel of 12 isolates (**Fig. 6c-d**). In general, all four variants neutralized this panel with comparable efficiencies, with a geometric mean IC_50_ of approximately 0.2 μg/ml (**Extended Data** Fig. 8b; **Table 1**). As expected CD4-Ig-v0, already modified with potency-enhancing mutations, was 7-fold more potent than WT CD4-Ig. The four variants v1-v4 ranged from 9- to 12-fold more potent than CD4-Ig-v0 and 60- to 80-fold more potent than WT CD4-Ig, with v3 modestly more potent than the other three variants. CD4-Ig-v1 was also more potent than R59K/K90R, implying that D63N mutation, the sole difference between these variants, contributes to neutralization potency. To evaluate the affinity maturation of individual B cell clones, we characterized three naturally emerging D1D2 variants (M1-1, M3-1, M6/7-1) with the greatest number of progenies in minimal-spanning trees generated from three mice (**Fig. 5c; Extended Data** Fig. 6). Each naturally emerging variant neutralized the global panel of isolates more efficiently than CD4-Ig-v0 with M3-1 and M6/7-1 approaching the geometric mean potency of v1-v4 (**Fig. 6d; Extended Data** Fig. 8a). The lower potency of M1-1 may be due to the absence of the K90R change. Notably, among both v1-v4 and the naturally emerging variants, greater improvement in potency was observed with HIV-1 variants more resistant to CD4-Ig-v0. The improved potency of v1-v4 and naturally occurring variants demonstrates affinity maturation of D1D2-OKT3-V_H_-modified BCR.

**Fig. 6.**
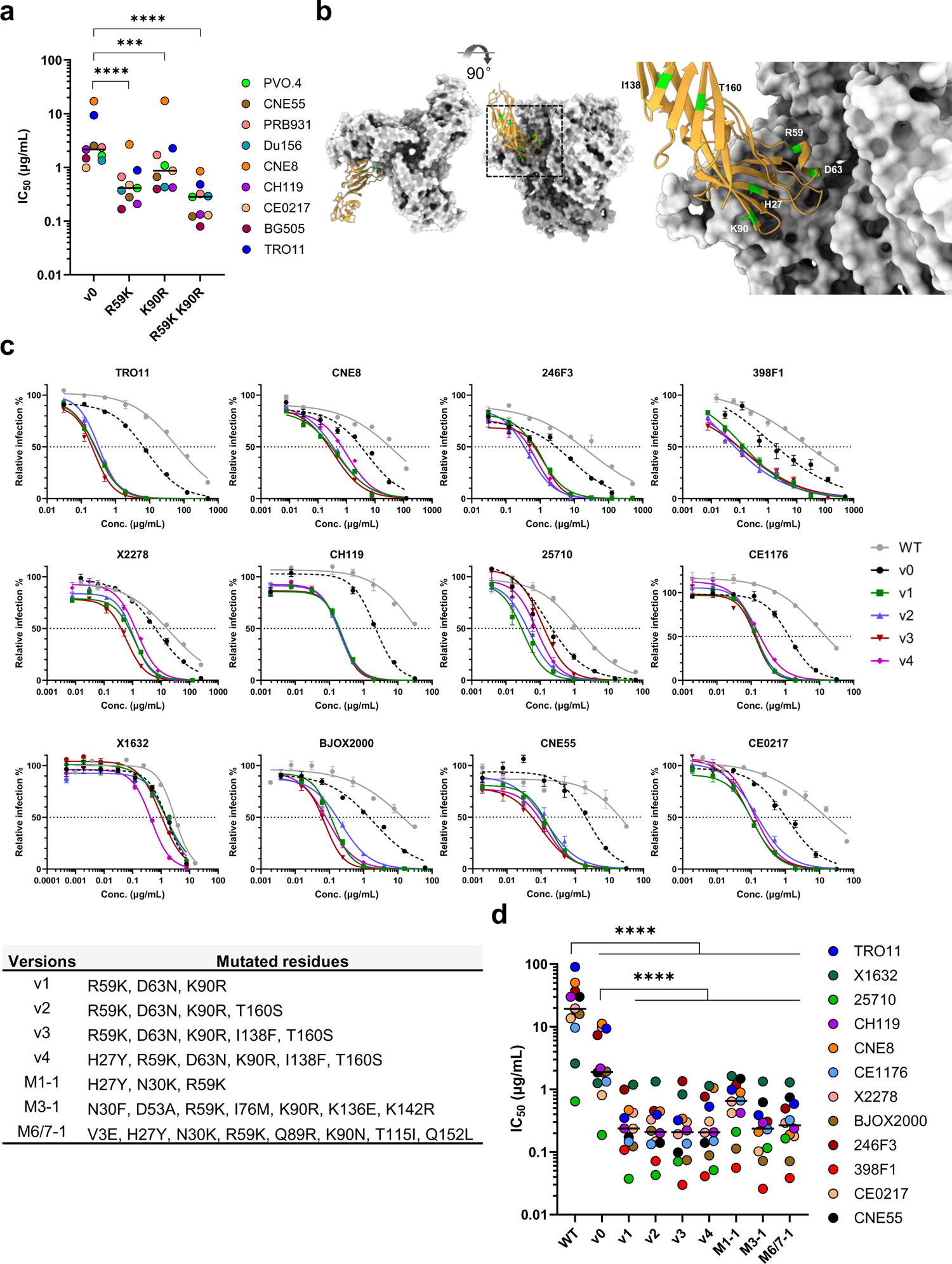
*In vivo* hypermutations in D1D2 improve the neutralization potency of CD4-Ig-v0. **a** Neutralization potency of CD4-Ig-v0 and its variants modified with R59K, K90R, or combined against the indicated isolates in TZM-bl assays. Curves are shown in **Extended Data** Figure 7a. Statistical significance was determined by two-way ANOVA with Dunnett’s multiple comparisons. **b** Location of selected D1D2 mutations (lime) shown on a structure of CD4 (yellow) bound to an HIV-1 Env trimer (light grey). Structure was adapted from pdb:5U1F. **c** Representative neutralization curves of CD4-Ig, CD4-Ig-v0 and v1-v4 variants bearing combinations of mutations listed below the figure, against a 12-isolate global panel of HIV-1 pseudoviruses. All curves were fitted with a variable slope four parameters dose response model. **d** IC_50_ of CD4-Ig, CD4-Ig-v0 and the indicated engineered (v1-v4) and naturally emerging variants (M1-1, M3-1, M6/7-1) against the global panel (****p < 0.0001). Each dot represents an average of two independent experiments. Center lines indicate median. Statistical significance in **d** was determined by repeated-measures two-way ANOVA with Dunnett’s multiple comparisons.

**Table 1.**
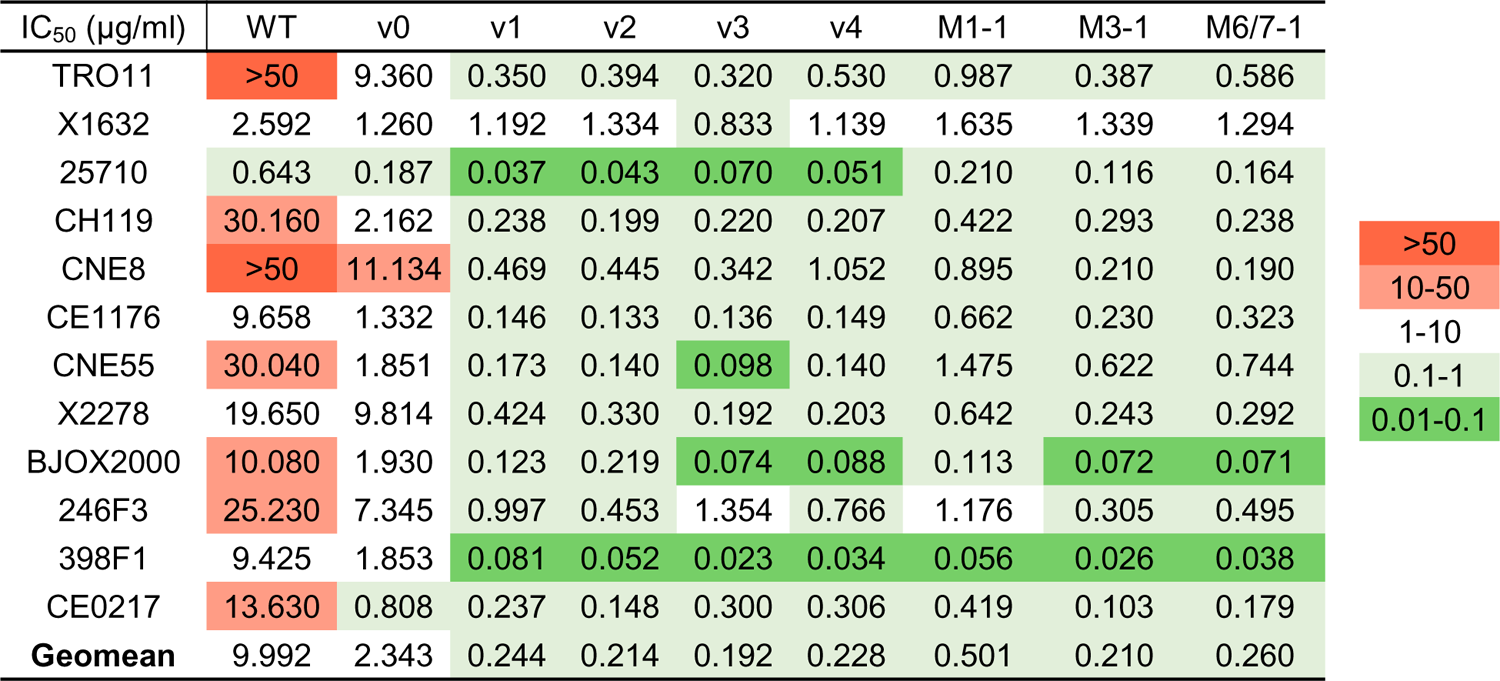
IC_50_ values of CD4-Ig variants plotted in **Fig. 6d**.

We also evaluated by surface plasmon resonance (SPR) whether these potency enhancements could be explained by higher affinity for Env. Accordingly, we compared CD4-Ig-v0 to v1-4 and the three naturally emerging variants (**Fig. 7a-b**). Each variant bound the ConM SOSIP with higher K_d_ than CD4-Ig-v0. Among v-1-4, v3 bound most tightly, with five-fold higher affinity than CD4-Ig-v0 (52 nM versus 267 nM). Although the potency of v3 against a global panel of isolates was greater than those of the three naturally emerging variants, these latter variants bound ConM SOSIP with even higher affinity. For example, M6/7-1 bound more tightly (22 nM) than CD4-Ig-v0, perhaps reflecting adaptation to the autologous ConM SOSIP immunogen. K_d_ differences between all variants and CD4-Ig-v0 were primarily determined by off-rate differences. Thus the enhanced potency of these CD4-Ig variants correlates with their slower off-rates and higher affinities for ConM SOSIP.

**Fig. 7.**
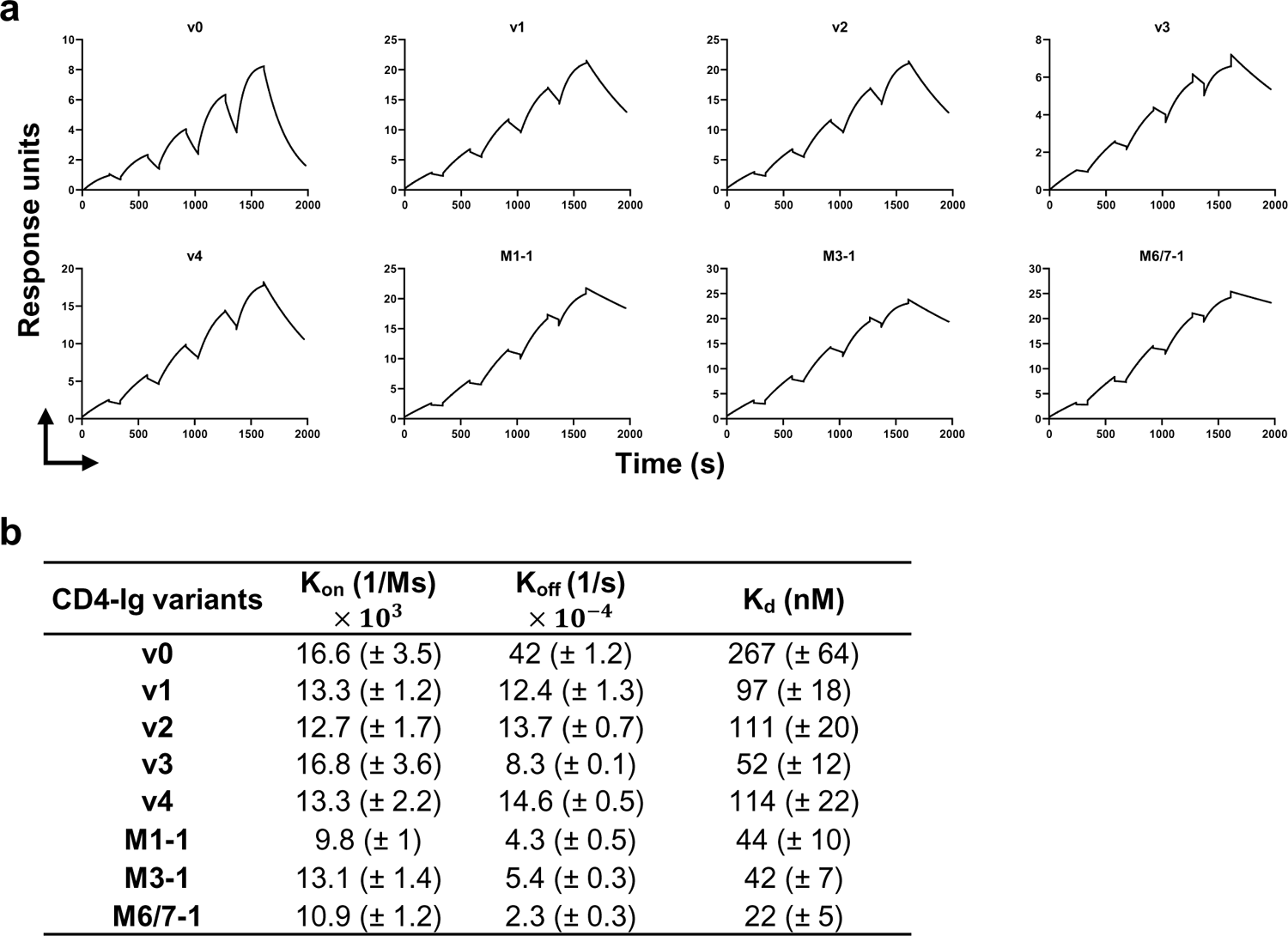
CD4-Ig variants bind Env trimers with higher affinity than CD4-Ig-v0. **a** Fitted sensorgrams show one of two replicates of the indicated CD4-Ig variant binding to 16055-ConM-v8.1 SOSIP trimers. Anti-human IgG antibodies were immobilized on the surface to capture CD4-Ig. After CD4-Ig capture, SOSIP protein was injected at concentrations of 800, 400, 200, 100, 50 nM in single-cycle kinetics at 25°C. The experimental data were fitted with a 1:1 Langmuir model. **b** Summary of K_off_, K_on_ and K_d_ for CD4-Ig variants. K_d_ is calculated from K_off_ and K_on_.

Often when proteins are affinity optimized with *in vitro* techniques such a phage- or mammalian display, the resulting variants lose properties critical to their *in vivo* activity, including their thermal stability and half-life. To determine if the half-life optimized CD4-Ig-v0 similarly reverted to the poor half-life of CD4-Ig with unmodified D1 and D2, we characterized the thermal stability and polyreactivity of WT CD4-Ig, CD4-Ig-v0, and v1 through v4. No differences in polyreactivity were observed between CD4-Ig-v0 and the CD4-Ig variants assayed, except in the case of v2, which was modestly but significantly more polyreactive than CD4-Ig-v0 (**Extended Data** Fig. 9a). Moreover, differential scanning fluorimetry showed that nearly every CD4-Ig variant retained the high thermostability of CD4-Ig-v0, significantly greater than WT CD4-Ig (**Extended Data** Fig. 9b). Similarly, the half-lives of all four variants in transgenic mice expressing the human FcRn receptor remained close to CD4-Ig-v0 (**Extended Data** Fig. 9c). The *in vivo* half-life of all variants remained significantly greater than wild-type CD4-Ig. We conclude that CD4-Ig variants bearing potency enhancing mutations selected *in vivo* retained characteristics important to their *in vivo* efficacy.

## DISCUSSION

We have recently shown that when heavy- and light-chain sequences of HIV-1 and SARS-CoV-2 neutralizing antibodies were introduced at the V(D)J-recombined regions of their respective mouse loci, the resulting BCR affinity matured after immunization^18, 32^. However, many protein-based therapeutics are not antibodies. Rather they derive from human proteins including cytokines and soluble forms of membrane-associated receptors. To determine if non-antibody biologics could be similarly improved through *in vivo* affinity maturation, we selected as a starting point a variant of CD4 domains 1 and 2 (D1D2) that was previously optimized for potency and long half-life when fused to an IgG Fc domain. However, we faced an immediate challenge: direct replacement of the mouse VDJ-recombined region with a sequence encoding D1D2 results in poor BCR expression, perhaps because heavy chains with unpaired CH1 domains are actively retained in endoplasmic reticulum^39^. To overcome this problem, we evaluated three strategies and found that a fusion of D1D2 with a murine heavy-chain variable region resulted in efficient expression. B cells modified to express this heavy-chain fusion protein class switched, entered germinal centers, and underwent robust somatic hypermutation and affinity maturation, implying that we did not significantly disrupt their underlying regulation.

Affinity maturation of the D1D2 domain allowed us to identify variants with markedly greater neutralizing potency against HIV-1 compared to our D1D2 starting point. This approach for improving the efficacy of protein therapeutics is qualitatively distinct from *in vitro* approaches such as phage, yeast, or mammalian display^2–4^. These latter approaches exclusively select for higher affinity for their targets, but they often identify proteins with undesirable properties, limiting their *in vivo* half-lives and efficacies^9, 10^. Rational design often led to reduced stability and yield of antibodies^11^, and CD4-Ig variants rationally designed for greater potency resulted in decreased yield and lower thermostability^31^. This concern is more pronounced with non-antibody biologics which are often short-lived in sera. In contrast, the *in vivo* selection process in germinal centers bypasses many of these pitfalls. Unstable, easily degraded, and poorly expressed variants are outcompeted by BCRs of comparable affinity but greater surface expression^12^. Peripheral tolerance eliminates BCRs that interact with self-proteins, including those with low affinity and high avidity interactions, for example with membranes or extracellular matrices^12^. Moreover, unlike *in vitro* techniques, *in vivo* affinity maturation occurs continuously and coordinately, building successively on selected high affinity variants, as is clear from our minimum-spanning tree analysis of our data. This process presumably has a strong evolutionary bias to sequences and structures that retain their *in vivo* activities. Moreover, this approach greatly simplifies affinity maturation of ligands of multi-pass membrane proteins such as G protein-coupled receptors expressed from mRNA. In contrast, *in vitro* approaches require reconstitution of these proteins in a liposome or nanodisc, complicating antigen production and selection of ligand variants. Thus *in vivo* affinity maturation offers a unique way to improve biologics while preserving properties important to their efficacy. Further, this approach helps establish a foundation for the therapeutic use of engineered B cells. It suggests that B cells expressing a non-antibody biologic could adaptively control an evolving pathogen.

*In vivo* affinity maturation thus allowed us to develop CD4-Ig variants with markedly greater potency against a global panel of HIV-1 isolates. In previous decades, CD4-Ig had been previously evaluated as a promising, difficult-to-escape therapy for HIV-1, with several phase I clinical trials establishing its safety in adults and children^26, 28, 29^. However, this original CD4-Ig faced three main limitations: it had prohibitively short *in vivo* half-life, its potency was lower than most HIV-1 broadly neutralizing antibodies^26, 29^, and it enhanced infection at low concentrations in cell-culture assays, by promoting interaction of the HIV-1 Env to the coreceptor CCR5. Since then, newer variants have been developed, including eCD4-Ig, whose C-terminal sulfopeptide improved potency while blockading binding of Env to CCR5^27, 31, 37, 40^. In recent years, more potent forms of CD4-Ig and eCD4-Ig have been developed with half-lives approximating many HIV-1 broadly neutralizing antibodies. We initiated these studies with one of these newer CD4-Ig variants, referred here as CD4-Ig-v0. We affinity matured the D1D2 of CD4-Ig-v0 *in vivo* and identified several mutations that recurred with high frequency across multiple mice, most notably R59K and K90R. Introduction of these mutations into CD4-Ig-v0 substantially improved its potency, while preserving its breadth and *in vivo* half-life. These CD4-Ig variants neutralized a global panel of HIV-1 isolates with IC_50_ values well below 1 μg/ml, similar to that of potent broadly neutralizing antibodies while maintaining the greater breadth of CD4-Ig. The most potent of these variants, CD4-Ig-v3, bound SOSIP trimers with a K_d_ of approximately 50 nM, five-fold slower than CD4-Ig-v0. Moreover, several natural emerging D1D2 variants bound the immunogen SOSIP with even higher affinity. Future *in vivo* studies will help determine if any of these potency enhancements can translate into more effective control of HIV-1 in humans.

This study pioneers the use of *in vivo* evolution to affinity mature non-antibody biologics. However, it has several limitations. First, nearly every amino-acid mutation derived from a single nucleotide change, implying that some potentially beneficial mutations have not been accessed. This limitation could be addressed with additional immunizations^41^, or by introducing libraries of variants. A related concern is that several neutral or unfavorable mutations emerged from AID hotspots. Introducing genes with more usefully distributed hotspots or using multiple templates with synonymous variation may increase the efficiency of affinity maturation. Another limitation is that, although we did not observe a significant loss of CD4-Ig half-life or thermostability, we did not observe any improvement. Thus, while this *in vivo* system might eliminate some poorly bioavailable variants, there may not be strong selection pressure to improve an already long-lived, well expressed biologic. In addition, B-cell tolerance might impede affinity maturation of biologics whose binding partners are homologous to their mouse orthologs. In many cases, this limitation could be overcome by employing alternative species or using knockout mice lacking this ortholog. Finally, we have focused primarily on mutations that emerge with high frequency in multiple mice, a strategy that clearly identifies useful individual mutations. However, this approach may overlook synergistic combinations of mutations. More thorough characterization of successful nodes identified through minimal spanning tree analysis may help identify these combinations.

Despite these limitations, this work describes a novel approach for refining biologics *in vivo*, bypassing several important limitations of *in vitro* methods. It provided the first example whereby non-antibody biologic affinity matures in the germinal center. We demonstrate the potential of this *in vivo* affinity maturation by markedly improving the potency of CD4-Ig while maintaining its half-life and breadth, suggesting that many protein therapeutics can be similarly improved.

## METHODS

### Mice

Mouse studies were approved and carried out in accordance with protocols provided to the Institutional Animal Care and Use Committee (IACUC) at UF Scripps (Jupiter, FL; approved protocol 17-026 and 21-010), at Boston Children’s Hospital (Boston, MA; approved protocol 00001921), and at The Jackson Laboratories (Sacramento, CA; approved protocol 20110). 9 to 12 weeks old CD45.1-positive mice (B6.SJL-Ptprca Pepcb/BoyJ, 002014) from The Jackson Laboratories were used as a source of splenic B cells. Age- and gender-matched CD45.2-positive C57BL/6J mice (Jackson Laboratories, 000664) were used as host mice for B cell transplantation and immunizations. Nine to ten weeks old SCID hFcRn transgenic mice (B6.Cg-Fcgrt^tm1Dcr^Prkdc^scid^Tg(FCGRT)32Dcr/DcrJ) housed at UF Scripps or The Jackson Laboratories were used for pharmacokinetics evaluation. No more than 5 mice or less than 2 mice were housed together. All procedures were performed on animals anesthetized with isoflurane.

### AAV production

HEK293T cells (CRL-3216) were seeded 18-22 h before transfection and grew to 60%-80% confluency in T225 flasks in Dulbecco’s Modified Eagle Medium (DMEM, Thermo Fisher Scientific, 10566016) containing 10% FBS (Thermo Fisher Scientific, 26140-079) at 37°C in 5% CO_2_. 174 μl (1 μg/ul) polyethylenimine (PEI, Polysciences, 49553-93-7) was vortex mixed with 1930 μl Opti-MEM (Thermo Fisher Scientific, 31985062); 58 μg of plasmids encoding the AAV rep and cap genes, adenoviral helper genes, and desired HDRT flanked by AAV ITRs were mixed with 1800 μl Opti-MEM. Opti-MEM containing PEI was added into Opti-MEM containing plasmids drop by drop followed by vertexing. The mixture was incubated at room temperature for 20-30 min. For each T225 flask, 4 ml mixture was added. Culture media was changed 18-22 h after transfection, and AAV was harvested after an additional 48 h. AAV was purified with the AAVpro Purification Kit (Takara, 6666) according to manufacturer’s instruction, and concentrated in PBS. Viral genome number was quantified by real-time PCR with AAVpro Titration Kit Ver.2 (Takara, 6233) according to manufacturer’s instruction, and capsid assembly was assessed by electrophoresis on reducing SDS-PAGE gels.

### Protein production, purification, and conjugation

Expi293F cells (Thermo Fisher Scientific, A14527) were maintained in Expi293™ Expression Medium (Thermo Fisher Scientific, A1435102) following the manufacturer’s instructions. Cells were diluted to three million/ml in preheated medium, and then transfected with FectoPRO reagent (Polyplus, 116-040). To produce gp120 proteins, plasmids expressing the gp120 and PDI (protein disulfide-isomerase) were co-transfected at 5:1 ratio in total of 80 μg / 100 ml culture. For CD4-Ig production, plasmids were added at 55 μg / 100 ml culture. For CD4-OKT3-IgG production, heavy chain and light chain plasmids were added at 1:1 ratio at 60 μg / 100 ml culture. Five days post-transfection, cell supernatant was harvested, centrifuged, and filtered. Monomeric ConM gp120 was captured with C-Tag affinity column (Thermo Scientific, 2943072005), ConM gp120 F10 and SOSIP were captured with house-made PGT145 or CH01 columns, while CD4-Ig was captured by HiTrap Mabselect SuRe columns (Cytiva, 11003493). SOSIP was eluted with gentle elution buffer (Thermo Scientific, 21027), buffer exchanged in desalting columns (Thermo Scientific, 89894) to EQB (10mM HEPES pH 8.0 in H2O with 500mM NaCl), and concentrated in 100K Amicon Ultra - 15 centrifugal filter devices (Sigma, UFC9050). Other proteins were eluted with IgG elution buffer (Pierce, 21004). pH was adjusted with 1/10 elution volume of 1M Tris-HCl, pH 9.0 (Thermo Fisher Scientific, J62085.K2). Elute was buffer exchanged and concentrated in PBS with 50K or 100K Amicon Ultra - 15 centrifugal filter devices (Sigma, UFC9050). Proteins were purified by size exclusion chromatography in the Superdex 200 Increase 10/300 GL column (Cytiva, 28990944) or HiPrep 26/60 Sephacryl S400 HR column (Cytiva, 28935605), concentrated and stored in PBS at −80°C. Fractions were assessed by SDS-PAGE to be >95% pure.

### mRNA lipid nanoparticle production

Codon-optimized genes encoding SOSIP variants fused to the Env C-terminal transmembrane domain (TM) sequence were inserted into a pUC vector with 5’ UTR, 3’ UTR, and polyA sequences under T7 promotor. For *in vitro* transcription (IVT), the DNA templates were linearized by digestion with HindIII and ScaI (NEB) and purified by phenol-chloroform extraction. IVT was then performed using MEGAscript® T7 Transcription Kit (Thermo Fisher Scientific, AMB-1334-5) according to the manufacturer’s instructions with modifications as using the CleanCap® Reagent AG (TriLink, N-7413) and m1-pseudouridine-5’-triphosphate (TriLink, N-1081). Template DNA was digested with Turbo DNase, and synthesized mRNA was purified by LiCl precipitation and 75% ethanol washing. After RNA qualification via electrophoresis in a denaturing agarose gel, double stranded RNA was then removed by cellulose (Sigma, C6288) depletion. The mRNA solution was then precipitated with 3M sodium acetate pH 5.2 and washed with isopropanol and then 75% ethanol. Finally, the RNase free water suspended mRNA were quantified and stored at −80°C before LNP formulation.

mRNA-LNP were formulated via mixing cartridges in the NanoAssembr BenchTop instrument (Precision) according to the manufacturer’s instruction. First, mRNA was diluted to 0.1-0.35 mg/ml in RNase free water with 25 mM sodium acetate pH 5.0 as the aqueous phase. The lipid phase was prepared with an N:P ratio of 6:1 by adding the lipid solutions SM-102 (MedChemExpress, HY-134541), DSPC (Avanti, 850365), cholesterol (Sigma, C8667), and PEG2000 PE (Avanti, 880150) at the molar ratio of 50:10:38.5:1.5 into ethanol. Aqueous phase and lipid phase were then transferred into individual syringes at 3:1 ratio and loaded to the pre-washed NanoAssemblr Benchtop Acetone Cartridge (Precision, NIT0058). LNP were formulated by mixing of the aqueous phase and lipid phase at a flow ratio of 3:1 and a flow speed of 6 ml/min. After formulation, LNP were buffer exchanged to PBS by dialysis and concentrated via ultrafiltration. mRNA encapsulation efficiencies and concentrations were determined with the Quant-iT RiboGreen RNA Assay Kit (Thermo Fisher Scientific, R11490). Diameters of LNP were measured by dynamic light scattering (DLS) using a Dynapro Naostar (Wyatt Technologies) and, finally, LNP were sterilized by filtration and stored at −80°C in PBS with 10% sucrose.

### Mouse splenic B cell activation and electroporation

Whole spleens from 9-12 weeks old CD45.1 positive donor mice were pulverized and mechanically crushed on the inner top of 70 μm cell strainers in RPMI 1640 medium (Thermo Fisher Scientific, 61870127) with 2% FBS (Thermo Fisher Scientific, 26140-079). After red blood cell lysis in a NH_4_Cl solution (BD Biosciences, 555899) at room temperature for 3 minutes, B cells were neutralized with Ca2+/Mg2+ free DPBS with 0.5% BSA (Miltenyi Biotec, 130-091-376) and 2 mM EDTA and then isolated using mouse B cell purification kit (Miltenyi Biotec, 130-090-862) and LS columns (Miltenyi Biotec, 130-042-401). Before electroporation, B cells were activated for 36-42 hours in RPMI 1640 medium with 10% FBS, 100 μM Non-Essential Amino Acids (NEAA, Thermo Fisher Scientific, 11140050), 1 mM sodium pyruvate (Thermo Fisher Scientific, 11360070), 10 mM HEPES (Thermo Fisher Scientific, 15630080), 55 μM 2-Mercaptoethanol (Thermo Fisher Scientific, 21985023), 100 units/ml penicillin and 100 μg/ml streptomycin (Thermo Fisher Scientific, 15140163), and 5 μg/ml anti-mouse CD180 antibody (Biolegend, 117708).

After activation, B cells were washed twice with Ca2+/Mg2+ free DPBS at room temperature. For each 100 μl of electroporation reaction using the nucleocuvette vessels (Lonza, V4XP-4024), 5 million cells were suspended in 74 μl of P4 Primary Cell solution (Lonza, VSOP-4096). In parallel, 3.12 μl of PBS, 1.26 μl of 1M NaCl, 1.12 μl of 250 μM Mb2Cas12a (produced in house), and 4.5 μl of 100 μM gRNA were mixed and incubated at room temperature for 15 min for RNP complex formation. The RNP were then incubated with 16 μl of 100 μM single strand DNA enhancer for 3 minutes at room temperature. The above 26 μl mixture was then mixed with the 74 μl of suspended B cells and transferred to the nucleocuvette vessels for electroporation in the Lonza 4D nucleofector under the DI-100 program. For larger scale electroporation in the 1 ml scale of Nucleocuvette Cartridge (Lonza, V4LN-7002), reactions were scaled up 10-fold for cell suspension, RNP, and donors. After electroporation, cells were rested for 10-15 min in nucleocuvette vessels or cartridges before transferred to preheated activation medium with 20% FBS without penicillin-streptomycin, which was added two hour later.

### Mouse B cell transplantation, immunization, and blood collection

Approximately 18 hours after electroporation, B cells were washed with prechilled Ca2+/Mg2+ free DPBS for three times and then suspended in prechilled DPBS with Ca2+/Mg2+ (Thermo Fisher Scientific, 14040133) and 5% horse serum (Cytiva, SH3007403HI). After filtration (Falcon, 352235) the number of cells was adjusted, and each mouse received 5 million cells in 100 μl buffer via retro-orbital injection under anesthesia with isoflurane. An aliquot of approximately 2 million cells were further cultured in RPMI 1640 medium with 10% FBS, 100 μM NEAA, 1 mM sodium pyruvate, 55 μM 2-mercaptoethanol, 10 mM HEPES, 100 units/ml penicillin and 100 μg/ml streptomycin, 5 μg/ml LPS, 10 ng/ml mouse IL-4 (PeproTech Inc, 214-14), and 2 μg/ml anti-mouse CD180 antibody for additional 24 hours to validate editing efficiency by flow cytometry. For protein immunization, 5 μg gp120 was adjuvanted with 10 μg monophosphoryl lipid A (InvivoGen, vac-mpla) and 10 μg saponin (InvivoGen, vac-quil) in PBS, and formulated into 250 μl per mouse. Mice were injected subcutaneously and intramuscularly. For mRNA-LNP, 0.5 μg in 20 μl was injected intramuscularly at each hind leg. Sera were collected one week after each immunization via submandibular bleeding.

### Analytical cytometry, cell sorting, and immunoglobulin repertoire sequencing

For flow cytometry and cell-sorting assays, spleens and lymph nodes were homogenized by mechanical disassociation and filtered through a 70-micron cell strainer. Red-blood cells were lysed with NH_4_Cl Buffer (BD Biosciences, 555899). B cells were then isolated with Pan B Cell Isolation Kit II (Miltenyi Biotec, 130-104-443) and LS columns (Miltenyi Biotec, 130-042-401), and resuspended in DPBS with 0.5% BSA and 2 mM EDTA. For cell sorting, antigen-specific B cells were stained by DAPI (BioLegend), CD45.1-FITC (BioLegend, 110706), IgG-PE (Biolegend, 405307), and ConM gp120-Fc conjugated with APC using the Lightning-Link (R) Fluorescein Antibody Labeling Kit (Novus, 705-0010). The antibody cocktail for GC B-cell staining consisted of CD45.1-FITC (BioLegend, 110706), CD19-BV605 (BioLegend, 115539) CD138-PE-Cy7 (BioLegend, 142513), GL7-PE (BD Biosciences, 561530), and CD38-AF700 (BioLegend, 102741).

Cells were incubated for 20 min on ice in dark, then washed and filtered before analysis or sorting on BD FACS LSR II, BD FACSAria III, BD FACSAria Fusion, or CytoFLEX S. Data were analyzed using FlowJo. CD45.1 and CD45.2 markers were used to distinguish donor and host B cells. Antigen-specific B cells were gated as singlet live CD45.1+ IgG+ gp120+; germinal center B cells were gated as singlet live CD19+ CD138− CD38− GL7+. At least 100,000 events per sample were analyzed.

Sorted B cells were lysed for RNA extraction by the RNeasy Micro Kit (Qiagen, 74004). First-strand cDNA synthesis was performed on 8 μl of total RNA using 5 pmol of IgM and IgG specific primers in a 20 μl total reaction with SuperScript III (Thermo Fisher, 18080044) according to the manufacturer’s protocol. Second-strand synthesis reactions were performed in 50 μl using HotStarTaq Plus (Qiagen, 203603) and 10 pmol of each primer tagged with unique molecular identifiers (UMIs). P5 and P7 flow cell-adaptor sequences and offset were introduced into dsDNA products with 10 pmol of i5i7 primers in a 50 μl total reaction volume for 20 PCR cycles. Libraries were verified in 1.5% agarose gel (Invitrogen, 16550100) and concentration was determined by NanoDrop. Libraries were optionally amplified for additional 6-12 PCR cycles if concentration was below 5 ng/μl. Finally, single indices for demultiplexing were added using the NEBNext Multiplex Oligos for Illumina (NEB, E7335S, E7500S, or E7710S) in 6-cycle PCR. All PCR products were purified by ExoSAP-IT (Thermo Fisher, 78201.1.ML) and SPRI beads (Beckman Coulter Genomics, SPRIselect). Bead-purified libraries were quality controlled on Agilent 2100 Bioanalyzer or 4200 TapeStation, and quantified by qPCR (Roche, KR0405). Pooled samples were sequenced using MiSeq 2×300 bp paired end reads (Illumina, MS-102-3003).

### Neutralization assays

Neutralizing activity was measured as the reduction in luciferase (Luc) reporter-gene expression after a single round of infection in TZM-bl cells. HIV-1 pseudoviruses were produced by co-transfecting Env plasmids with an Env-deficient backbone plasmid (pNL4-3 Env) in HEK293T cells grown in DMEM containing 10% FBS in a 1:3 ratio, using PEI MAX (Polysciences, 49553-93-7). Plasmids were acquired through the NIH HIV Reagent Program. Cell supernatants were harvested, centrifuged at 4000 g and filtered through 0.45μm filter 48 h after transfection and stored at −80°C. Neutralization assays were performed using pre-titrated HIV-1 Env pseudoviruses and TZM-bl cells growing in DMEM containing 10% FBS as previously described^45^. Briefly, 50 μl of titrated pseudoviruses were incubated with 50 μl of serially diluted antibodies/sera for 60 min at 37°C in 96-well flat bottom TC-treated plates (Corning, 353075). Subsequently, 100 μl of TZM-bl cells resuspended at 0.1 million/ml were added to each well and incubated at 37°C. At 48 h post infection, 100 μl supernatant was removed and cells were lysed in wells and subjected to firefly luciferase assays. Luciferase expression was determined using Britelight Plus substrate (PerkinElmer, 6066761). The luminescence signal was acquired with a Victor Nivo plate reader (PerkinElmer) or a GloMax Plate Reader (Promega). Percent infection was calculated using background-subtracted signals from wells containing virus only as a 100% infection reference, and neutralization curves were fitted by nonlinear regression using a four-parameter hill slope equation. Experiments were done in triplicate and repeated twice. IC_50_ and ID_50_ values were determined as the concentration or dilution required to inhibit infection by 50%.

### Surface plasmon resonance

Single-cycle kinetics analysis of CD4-Ig binding to the 16055-ConM-v8.1 SOSIP with C-terminal Spy-tag-2 (ST2) was performed at 25°C on a Biacore T100 (GE Healthcare). 1X HBS-EP containing 10 mM HEPES, pH 7.4, 150 mM NaCl, 3 mM EDTA, and 0.05% surfactant P-20 (Cytiva, BR100669) with 0.25% BSA and additional 150 mM NaCl was prepared and filtered. Mouse anti-human IgG Fc antibody (Novus, #NBP1-51523) was immobilized onto all flow cells of a CM5 sensor to ∼10,000 response units (RU) using the amine-coupling (Cytiva, BR-1000-50) method according to manufacturer’s instruction. SEC-purified CD4-Ig variants were captured at 5 nM and a flow rate of 5 μl/min for 60 s in sample flow cells. The capture level was kept to 5-25 RU to minimize mass transport effects and steric hindrance. The reference flow cells were left blank. A 2-fold increasing series of SEC-purified SOSIP (50 nM, 100 nM, 200 nM, 400 nM, 800 nM) was then injected into both the reference and sample flow cells at 50 μl/min for 240 s in a single cycle, resulting in a maximum of 30 RU. 360s of dissociation phase was followed by regeneration with 3M MgCl_2_. Sensorgrams were corrected with double reference by subtracting the response over the reference surface and the response of a blank injection from the sample binding responses. The association and dissociation phase data sets were globally fitted with Biacore Insight Evaluation software (v5.0) using a 1:1 Langmuir model, because the capture density and conformation of CD4-Ig prevents intra-protomer crosslinking of the SOSIP protein^46^. These kinetic binding studies were repeated twice on different sensor surfaces for each CD4-Ig variant.

### Pharmacokinetics in mice

SCID hFcRn transgenic mice were injected intravenously with 8Lmg/kg of CD4-Ig diluted in PBS. Serum samples were drawn from each mouse via the tail vein on days 1, 3, 6, 14, 21 and 30 post-injection. Sera were collected in tubes containing clotting activators and frozen at −80C right after processing.

### ELISA

To monitor the serum D1D2 concentration in B-cell engrafted mice by ELISA, high-binding 96-well plates (Corning, 3690) were coated overnight at 4°C with a rabbit anti-human CD4 antibody (Cell Signaling Technology, 93518) at a concentration of 2 μg/ml in PBS. To measure the CD4-Ig concentrations in serum samples in pharmacokinetic studies, plates were coated with 2 μg/ml of a mouse anti-human CD4 antibody (BioLegend, 300502) diluted in PBS overnight at 4 °C. Wells were washed two times with 0.05% Tween 20 in PBS, and blocked for 2.5 hours at room temperature with 150 μl of 4% BSA (Thermo Scientific, 37525) in PBS. After blocking, wells were loaded with 50 μl of serially diluted mouse sera, or standards of either SEC-purified CD4-OKT3-IgG for 1.5 hour or SEC-purified CD4-Ig for an hour at 37°C in duplicate. After five washes, wells were incubated with 50 μl of 1:2000 diluted horseradish peroxidase-conjugated goat anti-mouse IgG Fc antibody (Jackson ImmunoResearch,115-035-008) in 4% blocking buffer for 1.5 hours, or horseradish peroxidase-conjugated goat anti-human IgG Fc antibody (Sigma, A0170) diluted 1:20,000 in the blocking buffer for an hour at 37°C. Following eight washes, 50 μl of 1-Step Ultra TMB-ELISA substrate (Thermo Scientific, 34028) was added to each well and incubated at room temperature. The reaction was terminated with 50 μl of TMB Stop solution (SeraCare, 5150-0020). Absorbance at 450 nm was measured with a Victor Nivo plate reader (PerkinElmer) or a GloMax Plate Reader (Promega). The concentrations of the samples were determined by extrapolation from a standard curve made with a four-parameter nonlinear regression model in Prism.

### Differential scanning fluorimetry

Samples were prepared by mixing SEC-purified CD4-Ig with 25X GloMelt™ Dye (Biotium, #99843-20uL) to reach a final concentration of 10 nM CD4-Ig and 2X dye. Fluorescence was detected in SYBR channel of Bio-Rad CFX96. Temperature was incremented by 0.5°C per 30 s, from 25°C to 95°C. Melting curves were monitored for homogeneity. Melting temperatures were determined as the thermal transition points by the instrument software.

### Immunofluorescence assay on HEp-2 cells

Immunofluorescence assays were performed with ANA HEp-2 Test Kits (Zeus Scientific, FA2400EB) according to manufacturer’s instructions. Briefly, 100 μg/ml antibodies or CD4-Ig variants or manufacturer-provided control serum were incubated on HEp-2 cell slides at RT for 40 min, and then washed 3 times with PBS. FITC-conjugated anti-human IgG antibodies were coated to each well at RT for 25 min and slides were again washed. Slides were viewed using a Leica DMIL LED microscope at a 292ms exposure, and mean pixel intensity was measured by Image J.

### Bioinformatic analysis

All fastq files were initially processed with the in-house tool “dsa” (short for <u>d</u>eep <u>s</u>equencing <u>a</u>nalysis). Briefly, reads were first processed by trimming bases from the 3’ ends with Phred quality scores falling below 32. Subsequently, low quality reads that did not contain the adaptor sequences were discarded. Reads with shared UMI were merged into single reads. Merged reads had a minimum UMI group size of 2. dsa identifies open reading frames in deep-sequencing datasets and aligns them to CD4-Ig-v0 D1D2 nucleotide or amino acid templates using the Needleman–Wunsch algorithm. Following alignment and UMI group consensus formation, reads with undetermined nucleotides marked as N were removed. dsa outputs alignments of the reads to the template, tables of amino acid substitution frequencies, and summaries of coding and non-coding substitution frequencies. It further compiles lists of unique amino acid and nucleotide sequences used for further analysis including clustering and phylogenetic inference. Mutational frequency is calculated from these data as the number of sequences bearing a nucleotide mutation at a given position for each mouse divided by the total number of sequences from that mouse.

Clustering and phylogenetic inference was performed with a custom-made tool called “Dandelions”. Dandelions uses a maximum parsimony approach to generate the N-ary phylogenetic trees. The output represents a consensus of at least 500 minimum spanning trees constructed over the input sequences and a set of inferred ancestral sequences not present in the original data. Each node in the final tree corresponds to a unique amino acid sequence. Colored nodes are “centroids”, so designated based on the total of their non-coding variants and the number of their descendants in the tree. The root path of each centroid shares its color. The 18 largest centroids are rank ordered and labeled. The size of a node is proportional to the number of non-coding variants that share the amino-acid sequence of the node. The tree is rooted on the known ancestor, represented as black circle. The layout of the tree is determined by an interactive physics simulation where nodes are modeled as masses in a viscous medium connected by springs.

### Quantification and Statistical analysis

Flow cytometry analysis was carried out using FlowJo v.10 software. Throughout the study, statistical tests were performed using a nominal type I error rate of p = 0.05 and two-tailed tests where applicable. Mixed effects analysis, one- and two-way ANOVA were performed in GraphPad Prism 10. Flow cytometry data was analyzed using generalized linear mixed models in JMP Pro 17, where distribution was binomial, repeated measures aspect was included and generalized chisq/df value was monitored to make sure close to 1 to avoid overdispersion. Statistical information including n, mean, geometric mean, median, standard error of the mean, and statistical significance values are indicated in the figure legends. Normality and equal variances were examined in residual plot, homoscedasticity plot and QQ plot. Continuous outcome variables exhibiting a skewed distribution were log-transformed, and diagnostic plots were evaluated to assure assumptions are met in the final analysis. Mixed effects model with Geisser-Greenhouse correction was used for repeated measures. H-Šídák test (pairwise comparison) or Dunnett’s test (comparison to control) was performed following ANOVA and mixed effects analysis. Adjusted p-values for multiplicity were used in multiple comparison tests. Data were considered statistically significant at *p < 0.05, **p < 0.01, ***p < 0.001, and ****p<0.0001.

## CODE AVAILABILITY

Dandelions and dsa source code and binary installers can be downloaded from Github at https://github.com/baileych-bi/dsa-win64 and https://github.com/baileych-bi/dandelions/tree/Legacy.

## ACKNOWLEDGEMENT

The authors would like to gratefully acknowledge Dr. Haiyong Peng, Dr.Tonia Aristotelous, Dr. Nadine Elowe, and Dr. Shahab Bayani for assisting with running SPR experiments and data analysis, Dr. Hyeryun Choe for her advice and comments on the manuscript. Funding is provided by the following National Institutes of Health awards to M.F.: U19 AI149646; R21 AI152836; R01 DA056771; UM1 AI126623; R01 AI 154989; R37 AI091476.

## AUTHOR CONTRIBUTIONS

A.P., W.H. and M.F. conceived of the study and designed experiments; A.P., J.X., X.Z., N.S. performed and analyzed experiments; A.P., C.B. and B.H. designed and performed bioinformatic analysis; C.C.B. developed software tools; T.O., J.X., X.L., M.T. N.B., H.M., Y.Y., M.D.A. developed key reagents and provided useful insight; A.P. and G.C. conducted statistical analyses; M.F. provided funding support; A.P., W.H., M.F. wrote the manuscript.

## COMPETING INTEREST

A.P., W.H., T.O., Y.Y. and M.F. are inventors of a pending patent describing the *in vivo* affinity maturation of antibodies and biologics. C.C.B., M.D.A., and M.F. have equity stakes in Emmune, Inc., which developed CD4-Ig-v0. The authors have no other competing interests.

