## Supplemental figures 1 to 9 for "*In vivo* affinity maturation of the HIV-1 Env-binding domain of CD4"

**This file includes:**

**Extended Data Figs. 1 to 9**

Extended Data Figure 1

CD4-Cĸ-GS

CD4-OKT-VH

No HDRT

No gRNA

CD4-Cĸ-P2A

**a**

**c**

**b**

**Intron editing strategy**

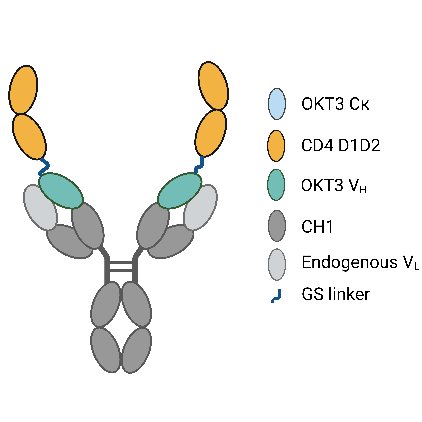

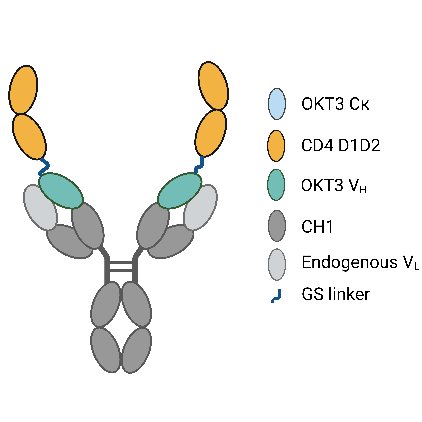

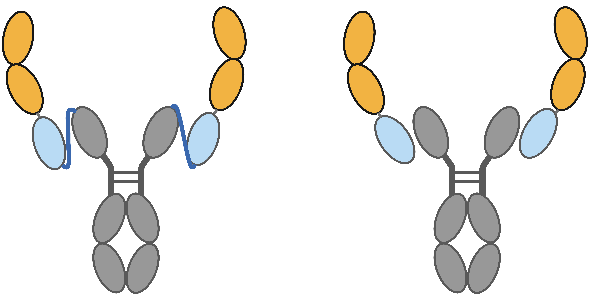

CD4-Cĸ-GS

CD4-OKT-VH

CD4-Cĸ-P2A

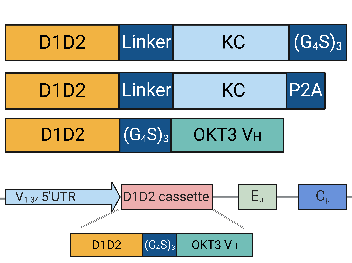

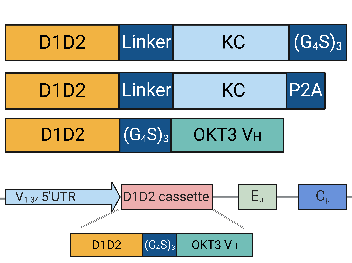

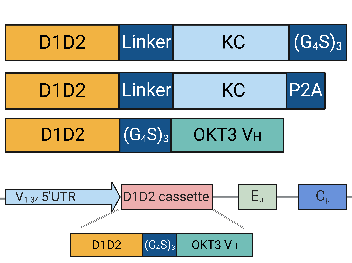

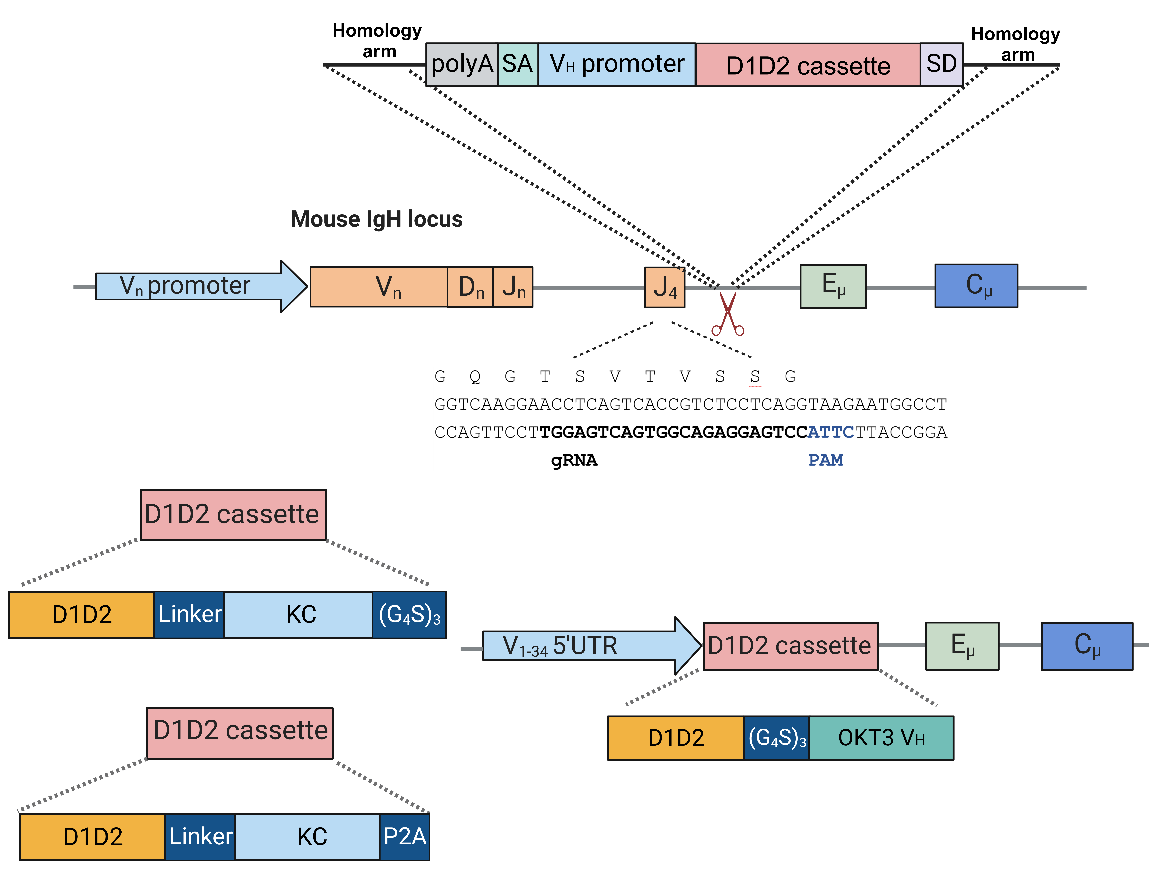

**Anti IgM**

**Anti CD4**

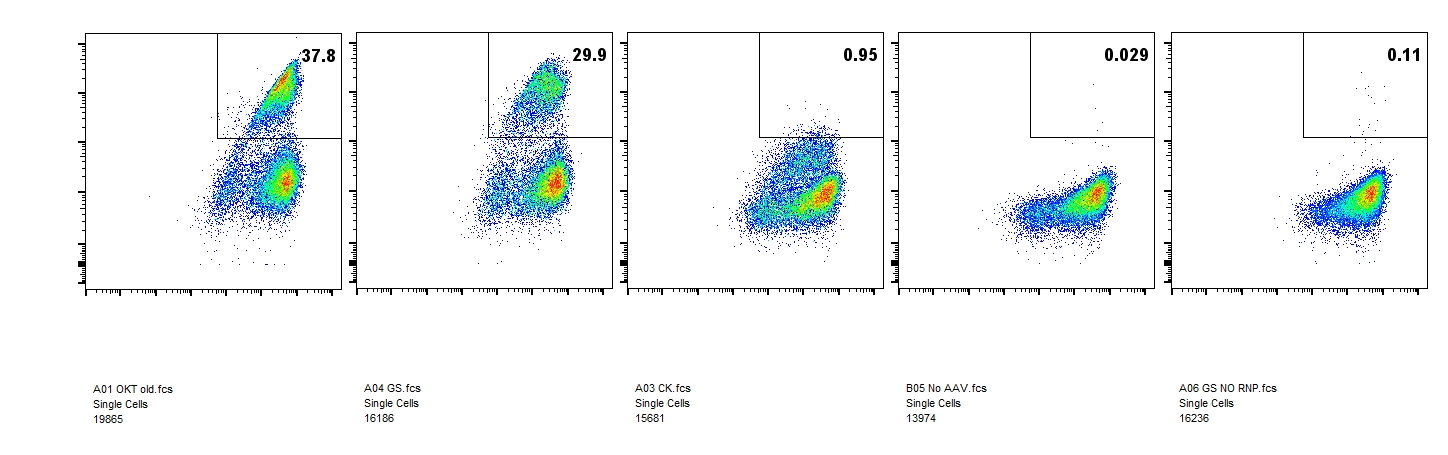

**Anti IgM**

**gp120**

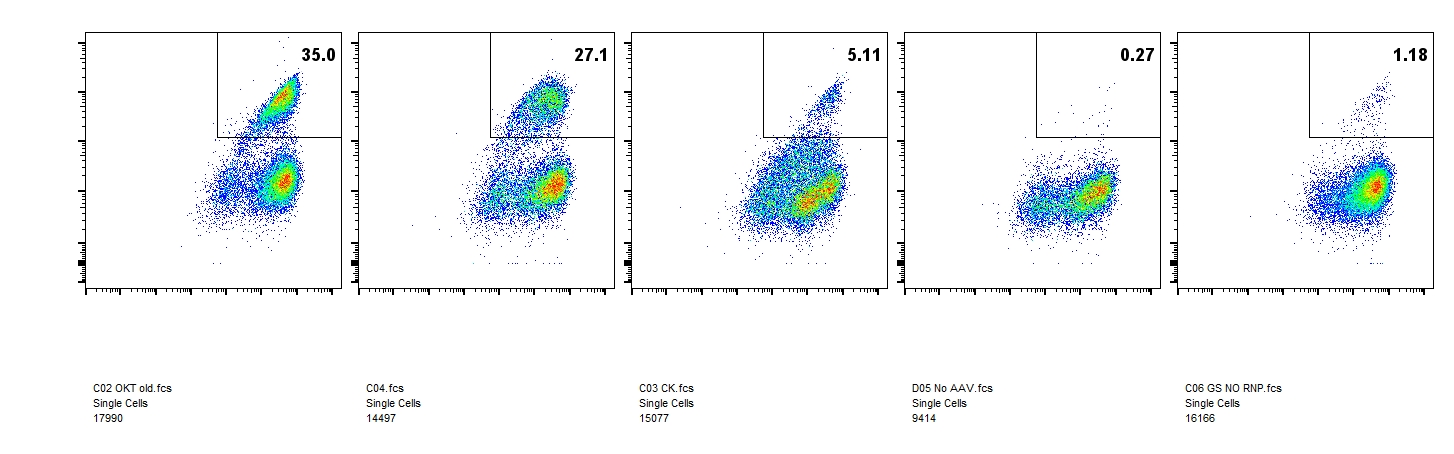

**Extended Data Fig. 1 Comparison of three D1D2 display strategies in primary murine B cells**. **a** Structure of edited BCRs showing three designs of CD4 D1D2 presenting B-cell receptors. Below each structure is a representation of its corresponding cassette. D1D2 (yellow)=) was attached to the N terminus of OKT3 heavy chain variable region (CD4-OKT-V_H_, green, left) or the OKT3 kappa light chain constant region (light blue, center and right). OKT3 kappa light chain constant region was linked to heavy chain constant region via a GS linker (CD4-Cĸ-GS, center), or non-covalently associated with heavy chain constant region by P2A cleavage (CD4-Cĸ-P2A, right). **b** The editing strategy used to express constructs in **a** in primary murine B cells. In this case, homology arms of the repair template complement an intronic region immediately downstream of JH4^18, 42-44^, and a cassette including a poly-A tail that terminates transcription of the VDJ region, a splice acceptor (SA), a heavy-chain promoter, a D1D2 construct shown in **a**, and a splice donor (SD). This ‘intron-editing’ strategy enables efficient introduction of these cassettes, but interferes with efficient somatic hypermutation, preventing affinity maturation of the expressed B-cell receptor^18^. An alternate strategy, used hereafter, is represented in Fig. 1b. **c** Flow cytometric analysis of singlet viable B cells edited using the strategy in **b** were stained with anti-IgM antibodies and either anti-human CD4 antibodies or gp120 at 48 h post-editing. HDRTs were delivered by 10^5^ MOI rAAV-DJ with homology arms targeting J4 and the intron immediate downstream J4. **a**, **b** are created with BioRender.com.

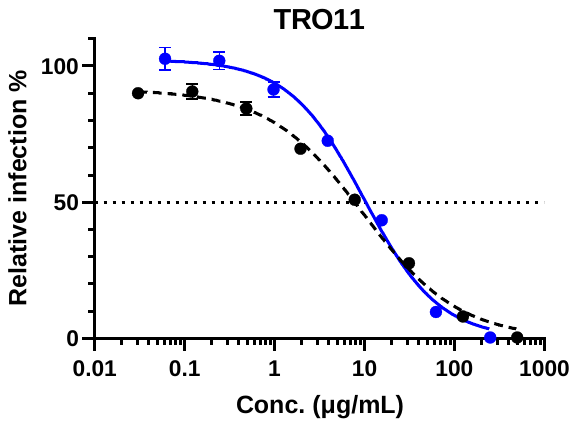

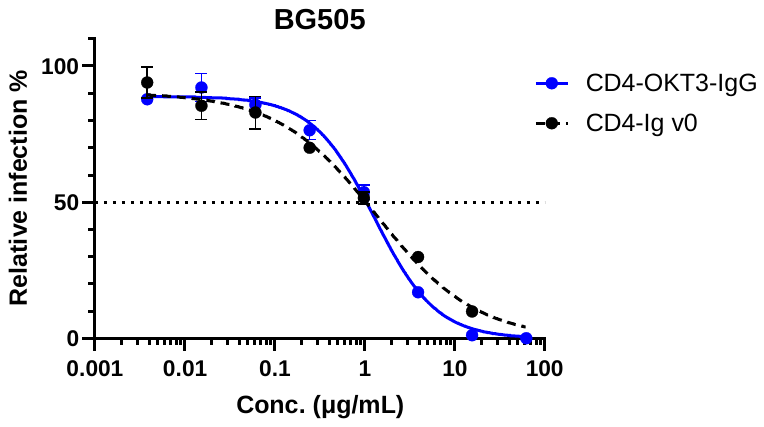

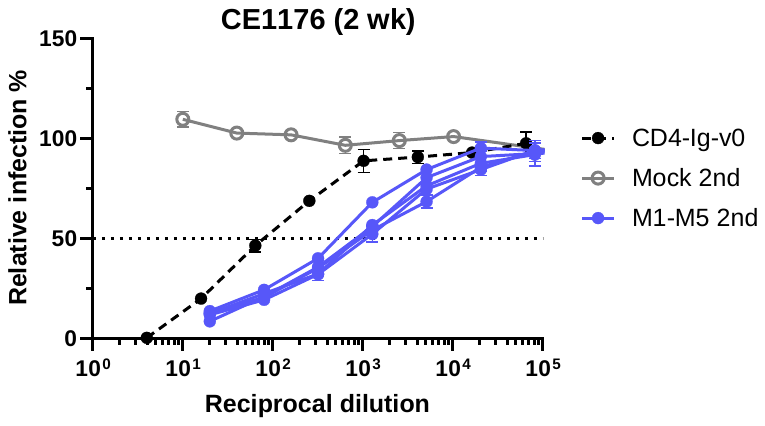

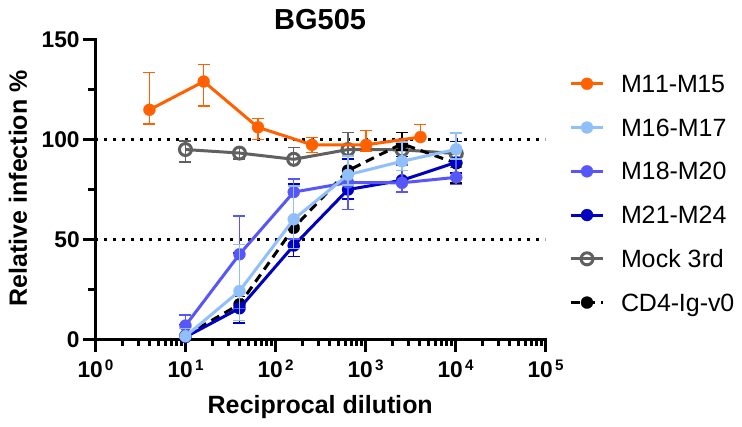

Extended Data Figure 2

**d**

**b**

**a**

**c**

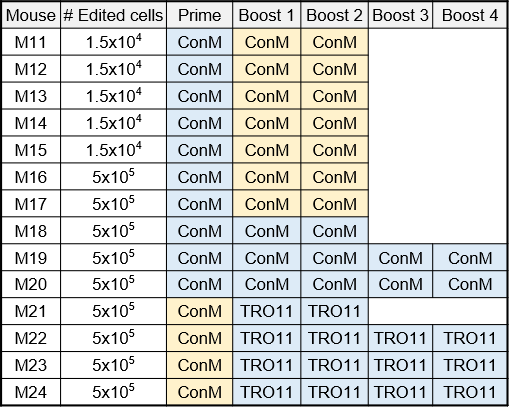

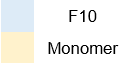

**Extended Data Fig. 2 Neutralization of mouse serum and CD4-OKT-IgG to various pseudoviruses and immunization regimen of mice receiving protein immunogens**. **a** Neutralizing response against HIV-1 CE1176 pseudovirus. Sera collected from mice (n = 5) engrafted with edited cells and immunized twice by mRNA-LNP at two-week interval were tested in TZM-bl assays and represented as individual curves. Sera from mice (n = 3) engrafted with unedited CD45.1 B cells and immunized on two-week interval (grey dots) served as negative controls. Dots and error bars of negative controls indicate median and interquartile range. Positive control CD4-Ig-v0 was mixed with normal mouse serum at 125 μg/ml. Error bars represent SEM. **b**. Neutralizing potency of CD4-OKT3-Ig. CD4-OKT3-Ig purified by size exclusion chromatography (SEC) was compared with CD4-Ig-v0 for IC_50_ for BG505 and TRO11 pseudoviruses (PVs). **c** Engraftment and immunization regimen for additional mice edited with the same method as **Fig. 1** but vaccinated with protein antigens. Mice were engrafted with 15,000 (M11 through M15) or 500,000 (M16 through M24) successfully edited B cells, as determined by flow cytometry analysis. Mice were vaccinated with protein antigens (monomeric or decametric ConM gp120, and monomeric TRO11 gp120) at two-week interval. Schedule of blood and tissue collection is the same as in **Fig. 2a**. Decameric antigens are indicated in blue, and monomeric antigens in yellow. **d** Sera neutralizing response of mice in **c** against BG505 PV. Dots and error bars indicate median and interquartile range. Sera from mice (n = 3) engrafted with unedited CD45.1 B cells and immunized with decametric ConM gp120 (grey dots) serve as negative controls. Note that protein immunization induced neutralizing response only in groups engrafting with a higher number of cells.

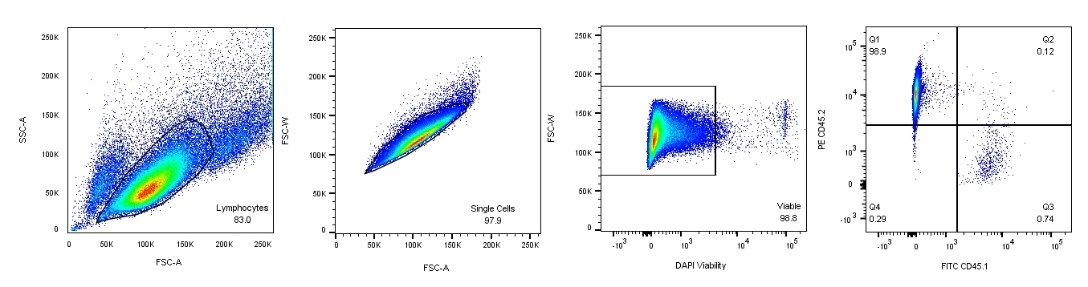

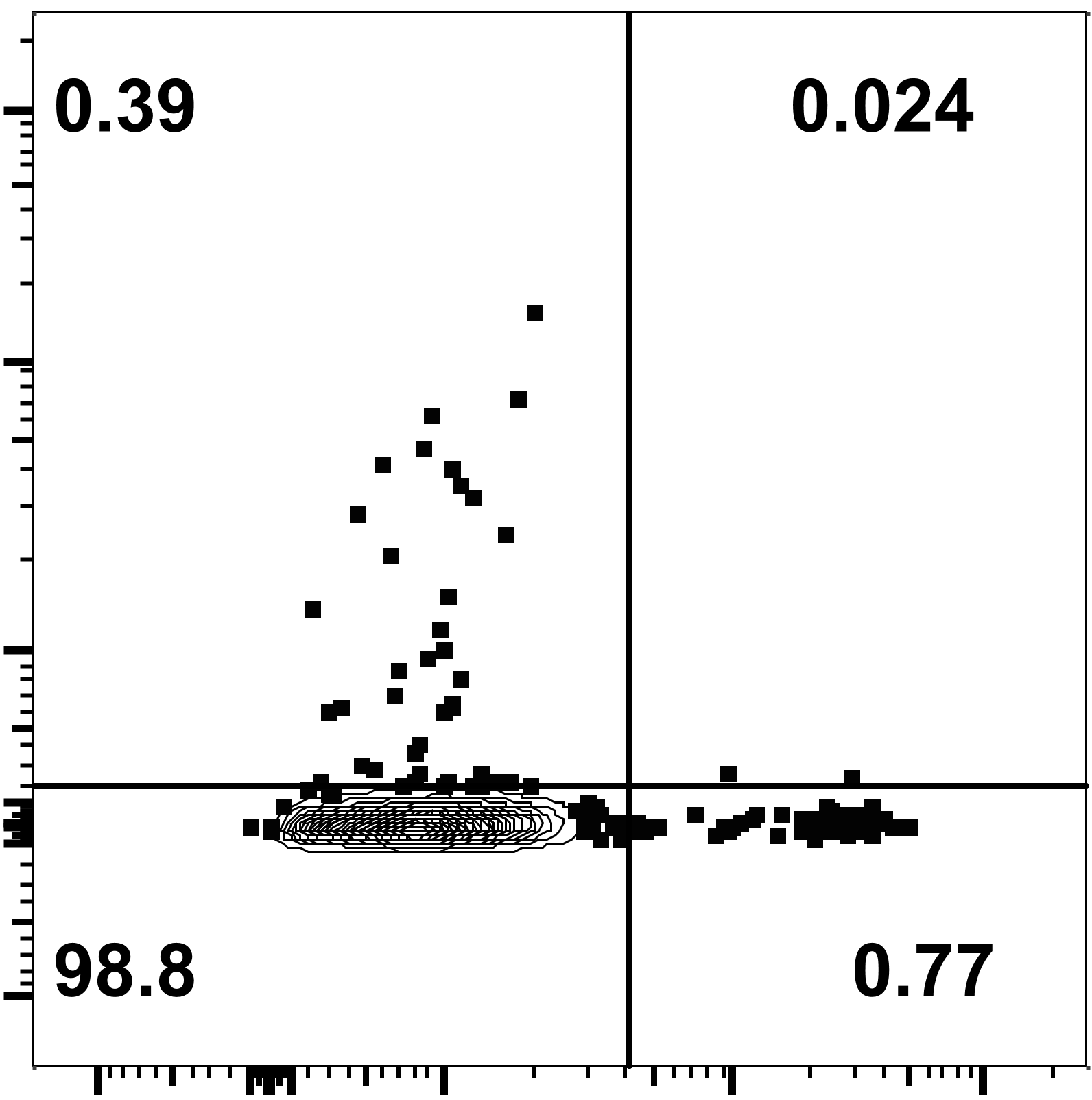

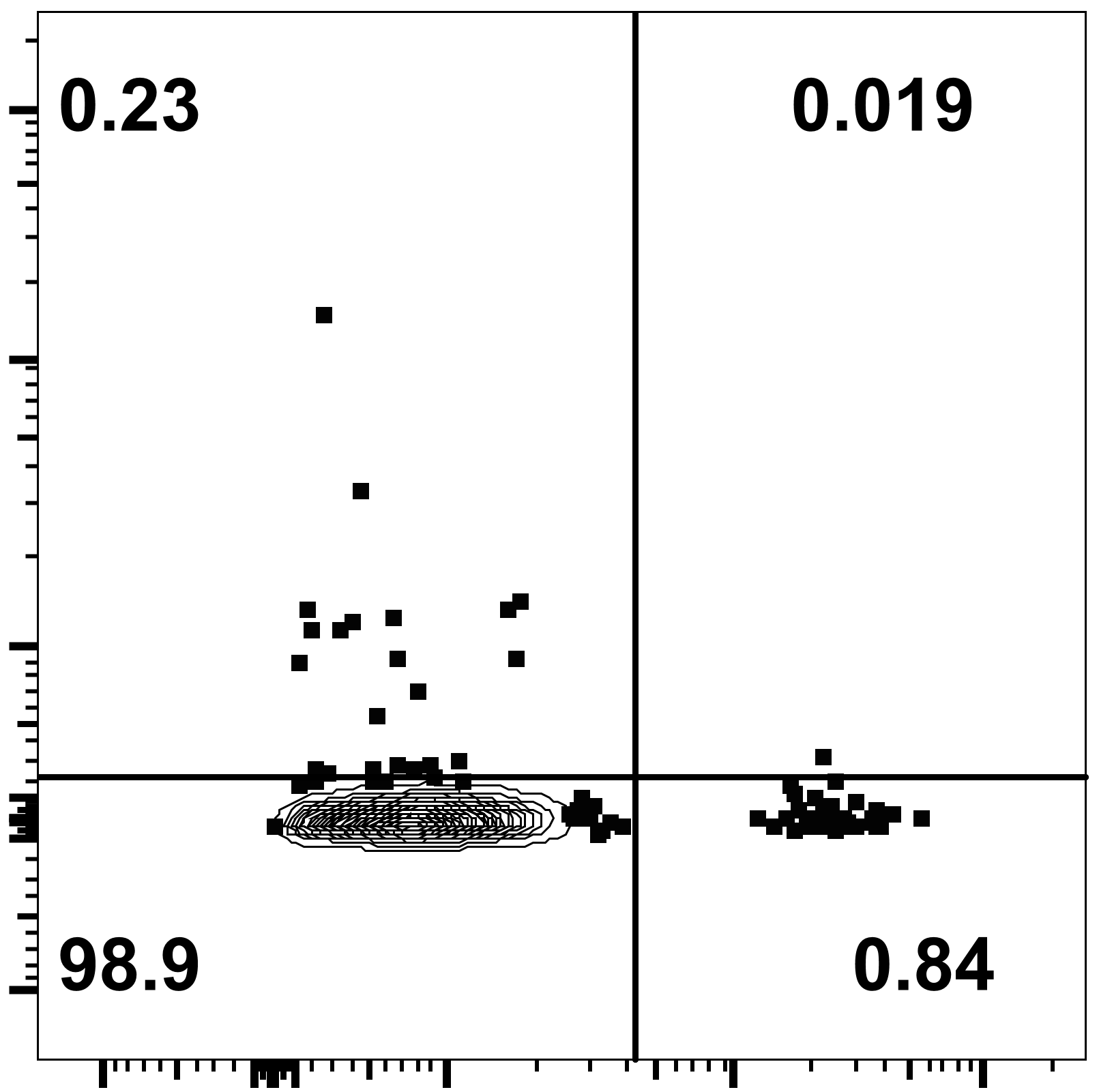

Extended Data Figure 3

Purified B cells	Singlets	Viable

**a**

**b**

CD45.1+ gp120+

IgG+

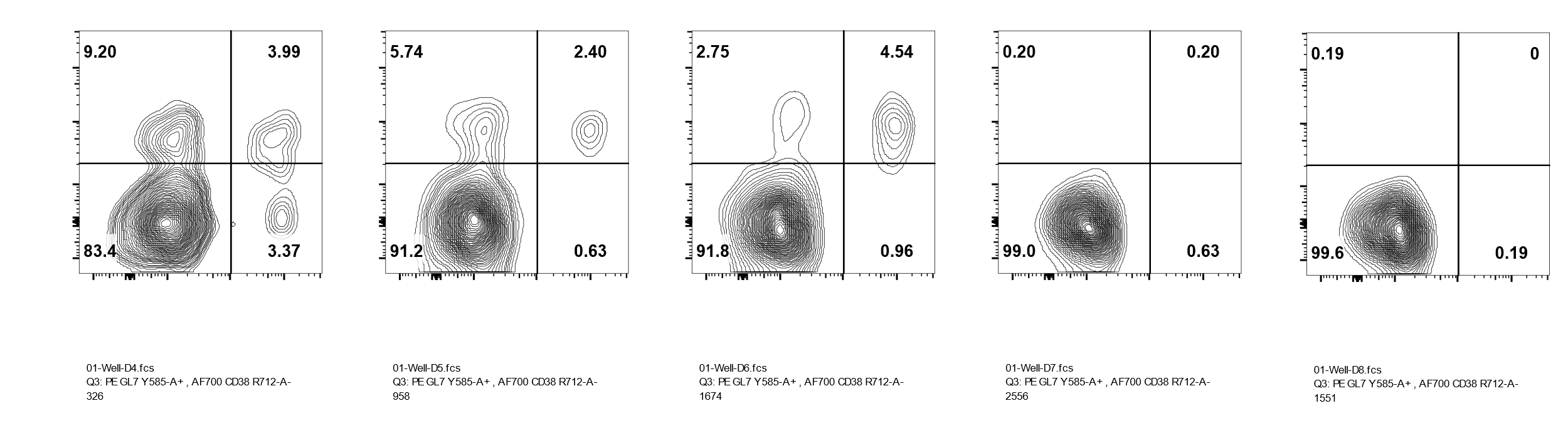

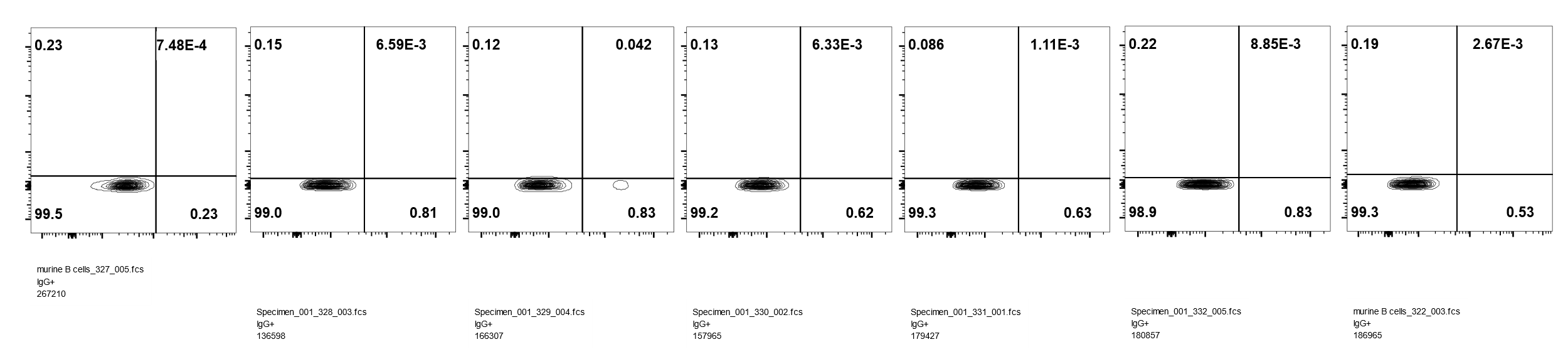

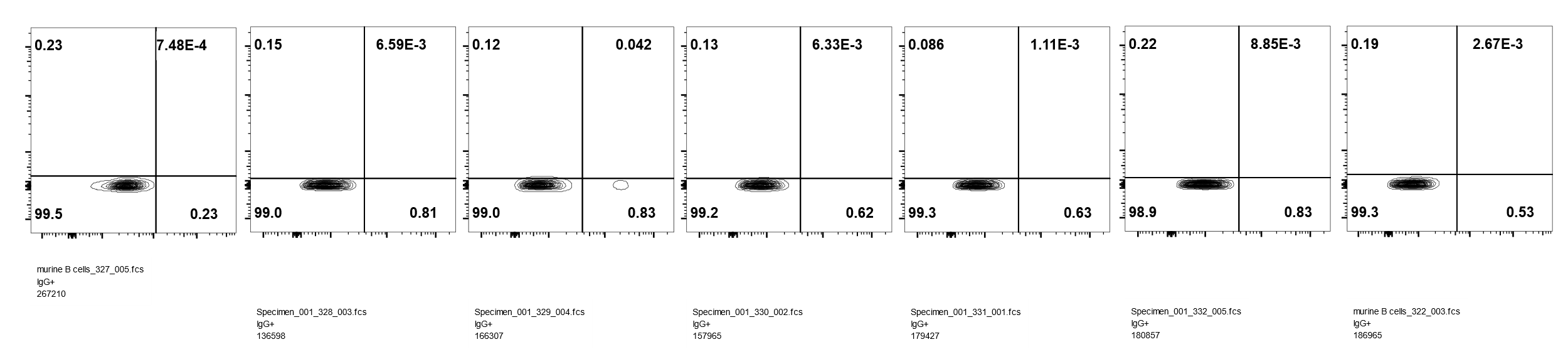

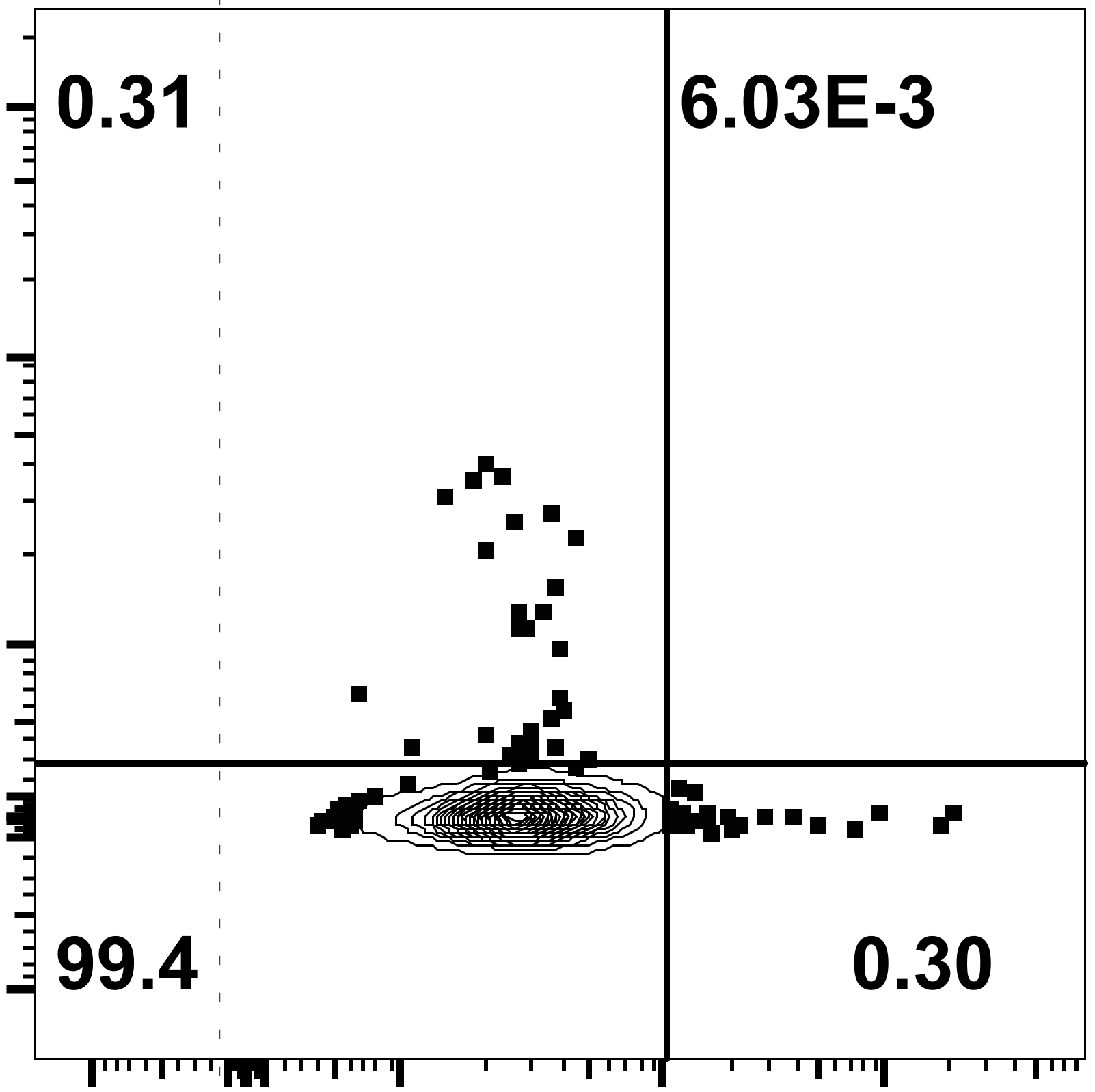

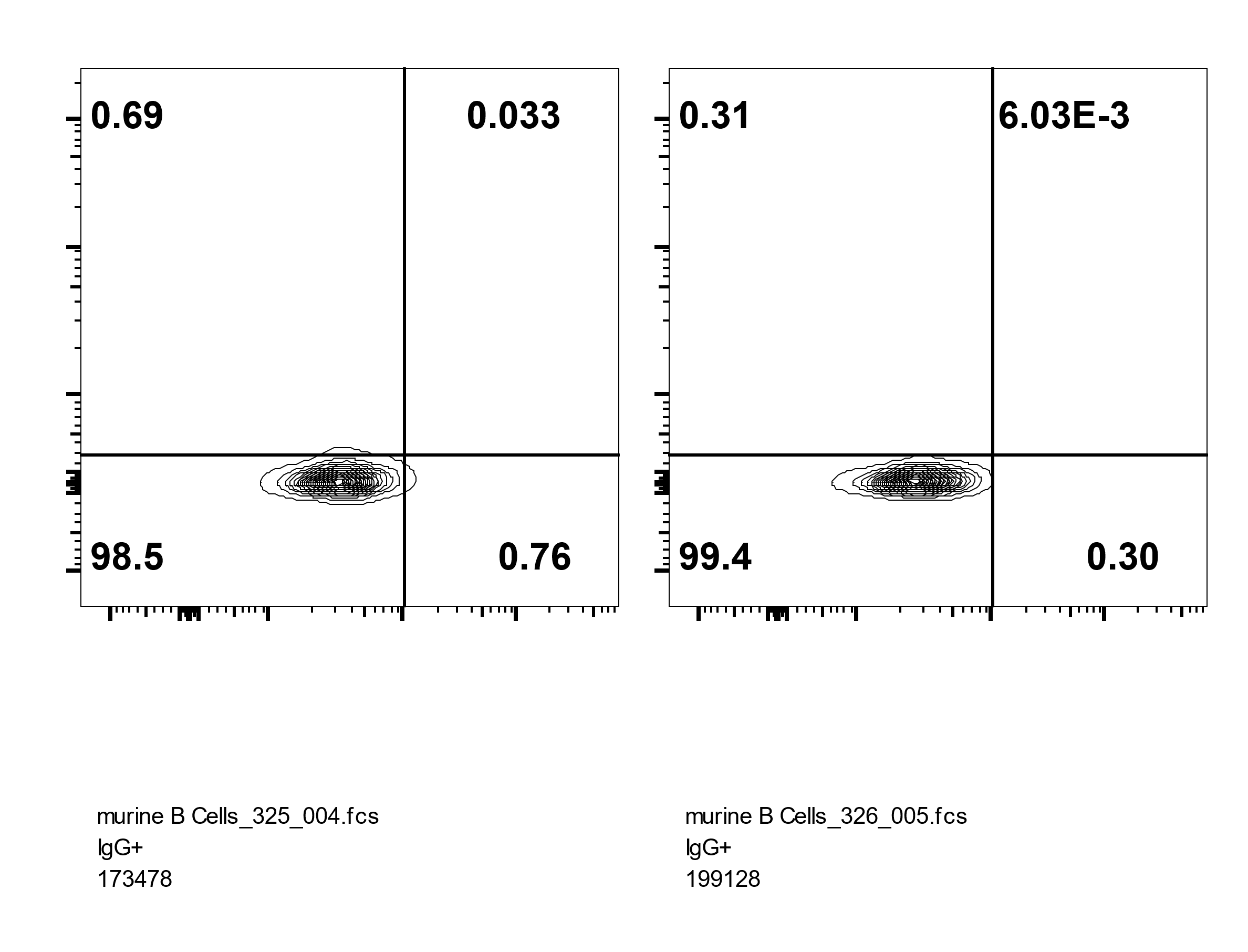

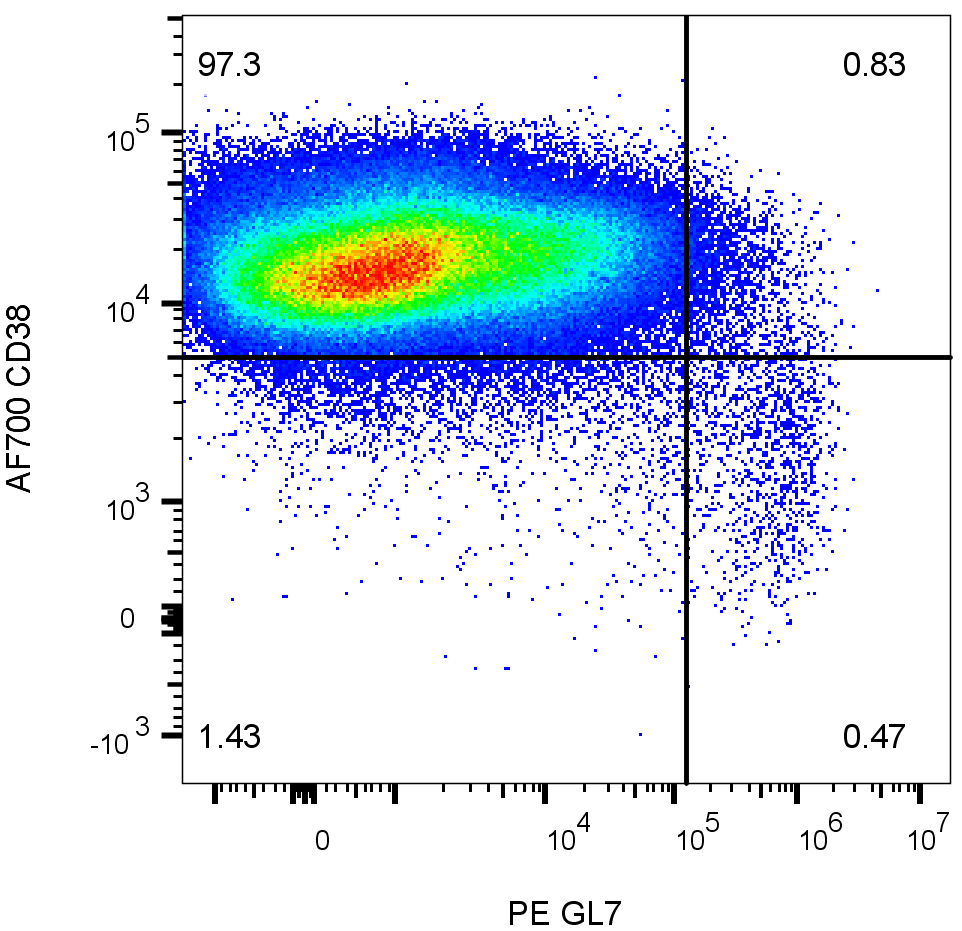

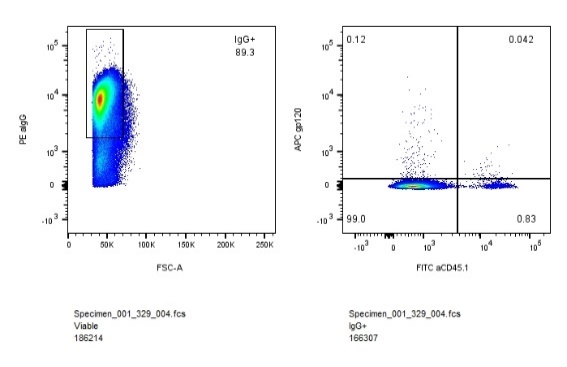

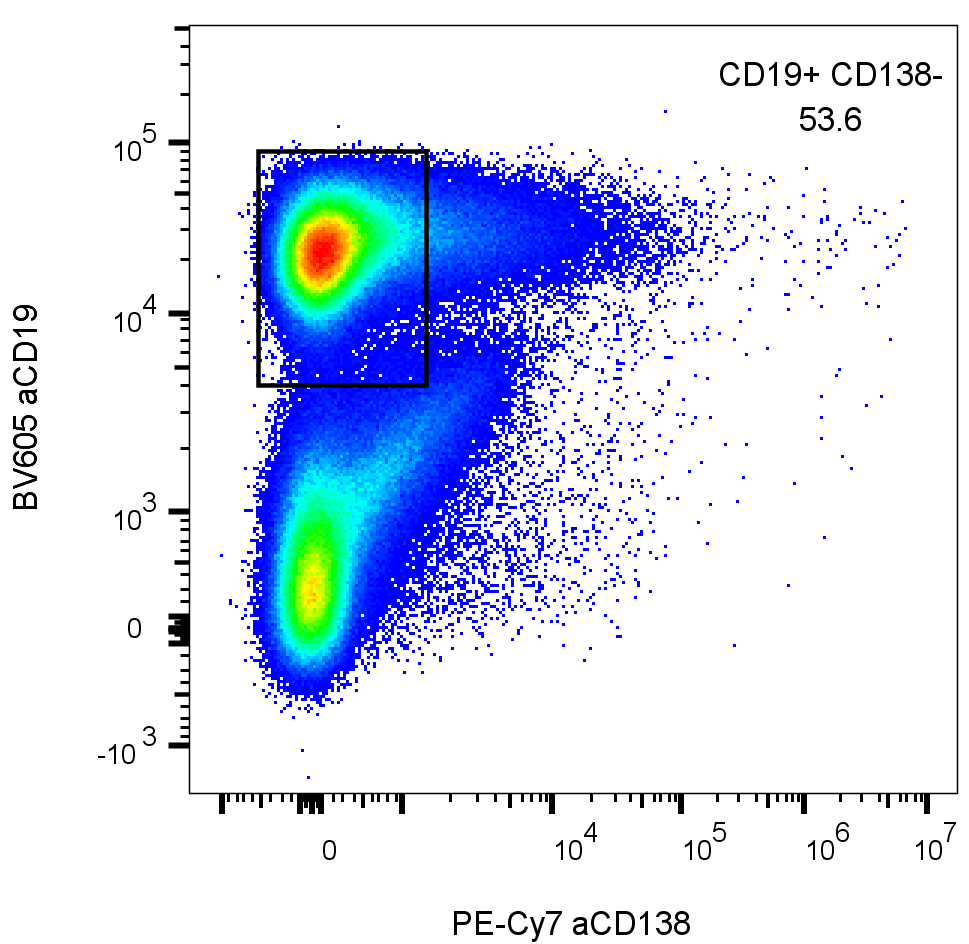

M7

M9

**gp120**

**Anti CD45.1**

**Anti CD45.1**

**gp120**

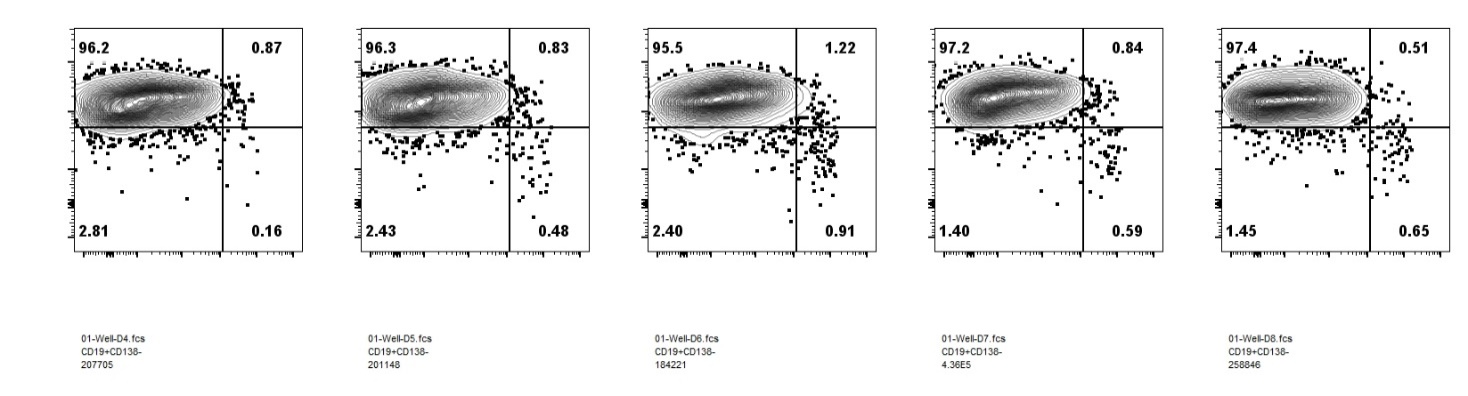

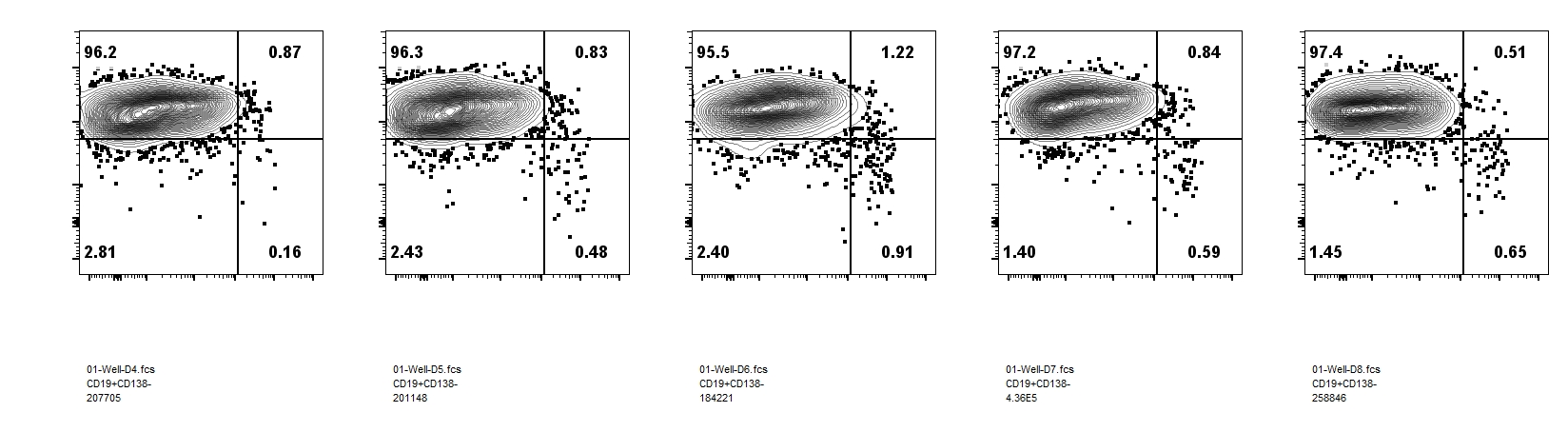

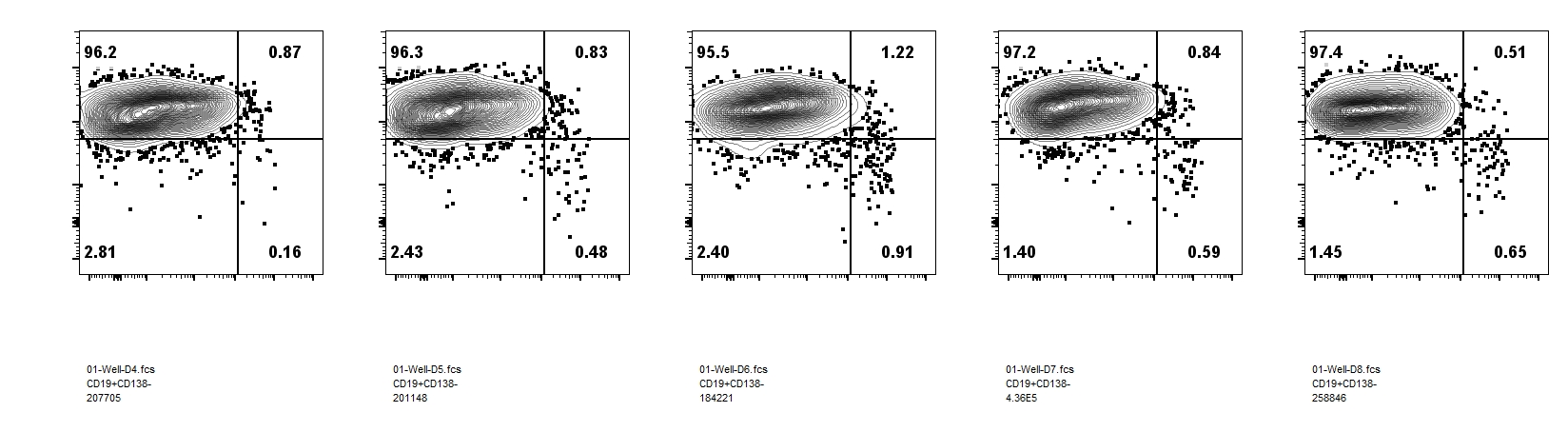

**Anti GL7**

**Anti CD38**

**d**

M7

M6

M9

M10

M8

M3

M2

M1

M4

M5

FSC-A

Anti CD138

Anti GL7

Anti CD38

Anti CD45.1

gp120

**GC staining**

Anti CD19

Anti IgG

FSC-W

DAPI

FSC-W

FSC-A

FSC-A

SSC-A

CD19+ CD138_low_

GL7+ CD38-

Mock

M1

M2

M9

M7

**c**

**Extended Data Fig. 3. Analysis of antigen binding B cells after three immunizations.** **a** Gating strategy for flow cytometry analysis of donor and recipient antigen binding B cells and germinal center B cells in recipient mice. For **Fig. 3a-b**, mature B cells were gated as singlet viable IgG+ cells. For **Fig. 3c-d**, GC B cells were gated as singlet viable CD19+ CD138low cells. **b** Flow cytometry analysis of antigen binding B cells in singlet viable IgG+ donor (CD45.1) and recipient (CD45.2) population for **Fig. 3a-b**. **c** Flow cytometry analysis of germinal center (GL7+ CD38-) B cells from spleen and lymph nodes harvested four days after three immunizations. M1 and M2 were from 2 wk group; M7 and M9 were from 4 wk group. **d** Flow cytometry analysis of antigen binding GC B cells in M7 and M9, as quantified in **Fig. 3d**. M1 and M2 were shown in **Fig. 3c**.

Extended Data Figure 4

| **Amino acid** | **Nucleotide** | **Frequency %** | **A** | **T** | **C** | **G** | **AID motif** | **Found in # samples** | **# sample > 10% frequency** |
| --- | --- | --- | --- | --- | --- | --- | --- | --- | --- |
| **90-K** | **269-A** | 63.93 | 0.0% | 13.4% | 6.8% | 79.9% | N | 9/9 | 9/9 |
| **59-R** | **176-G** | 53.95 | 100.0% | 0.0% | 0.0% | 0.0% | N | 8/9 | 8/9 |
| **90-K** | **268-A** | 22.81 | 0.0% | 81.7% | 15.6% | 2.7% | N | 8/9 | 6/8 |
| **30-N** | **88-A** | 16.69 | 0.0% | 37.4% | 62.5% | 0.1% | Y | 8/9 | 6/8 |
| **108-L** | **324-G** | 16.11 | 51.5% | 1.7% | 46.9% | 0.0% | Y | 5/9 | 4/5 |
| **99-G** | **297-G** | 12.84 | 78.2% | 5.9% | 15.9% | 0.0% | Y | 1/9 | 1/8 |
| **94-Q** | **282-G** | 12.75 | 95.3% | 3.6% | 1.0% | 0.0% | Y | 5/9 | 2/5 |
| **50-K** | **150-G** | 10.48 | 99.5% | 0.0% | 0.4% | 0.0% | Y | 4/9 | 2/4 |
| **64-Q** | **192-A** | 8.06 | 0.0% | 0.1% | 0.0% | 99.9% | Y | 4/9 | 1/4 |
| **13-E** | **39-G** | 7.88 | 99.2% | 0.0% | 0.8% | 0.0% | Y | 4/9 | 2/4 |
| **30-N** | **90-C** | 7.53 | 25.3% | 12.1% | 0.0% | 62.6% | Y | 8/9 | 1/8 |
| **68-P** | **204-T** | 7.20 | 0.2% | 0.0% | 1.9% | 97.9% | N | 2/9 | 1/2 |
| **148-Q** | **444-G** | 7.10 | 100.0% | 0.0% | 0.0% | 0.0% | Y | 4/9 | 2/4 |
| **73-N** | **217-A** | 6.54 | 0.0% | 1.1% | 0.0% | 98.8% | Y | 2/9 | 1/2 |
| **72-K** | **214-A** | 6.51 | 0.0% | 0.0% | 96.2% | 3.8% | N | 2/9 | 2/4 |
| **103-N** | **309-C** | 6.13 | 0.1% | 88.7% | 0.0% | 11.2% | Y | 6/9 | 3/6 |
| **42-S** | **125-G** | 6.12 | 1.6% | 0.1% | 98.3% | 0.0% | Y | 7/9 | 1/7 |
| **153-D** | **459-T** | 6.06 | 0.0% | 0.0% | 100.0% | 0.0% | N | 2/9 | 1/2 |
| **43-F** | **128-T** | 5.94 | 99.9% | 0.0% | 0.1% | 0.0% | N | 3/9 | 2/3 |
| **142-K** | **426-A** | 5.68 | 0.0% | 44.8% | 0.1% | 55.2% | Y | 5/9 | 1/5 |
| **138-I** | **414-C** | 5.54 | 0.1% | 99.9% | 0.0% | 0.0% | N | 2/9 | 2/2 |
| **48-P** | **144-T** | 5.37 | 68.5% | 0.0% | 31.5% | 0.0% | N | 3/9 | 2/3 |
| **150-E** | **450-A** | 5.30 | 68.5% | 0.0% | 31.5% | 0.0% | Y | 3/9 | 1/3 |
| **93-V** | **277-G** | 5.27 | 98.7% | 0.1% | 1.2% | 0.0% | Y | 2/9 | 1/2 |
| **58-R** | **174-G** | 5.13 | 99.7% | 0.3% | 0.0% | 0.0% | N | 2/9 | 1/2 |
| **27-H** | **79-C** | 5.03 | 0.3% | 99.7% | 0.0% | 0.0% | N | 7/9 | 1/7 |

Coding

Silent

**Extended Data Fig. 4 Analysis of AID motif and convergence of the top nucleotide mutations among 10 mRNA-immunized mice**. The first two columns show the original amino acid and nucleotide. AID hotspot motifs are defined as DGYW / WRCH (R=A/G, Y=C/T, and W=A/T). Mutation frequency is calculated as an average from 10 mice immunized by mRNA-LNP every two or four weeks. Blue bars indicate percentage of mutation. Y/N indicates whether the mutation is located at AID motifs. Coding mutations are labeled in light blue; silent mutations are labeled in light orange.

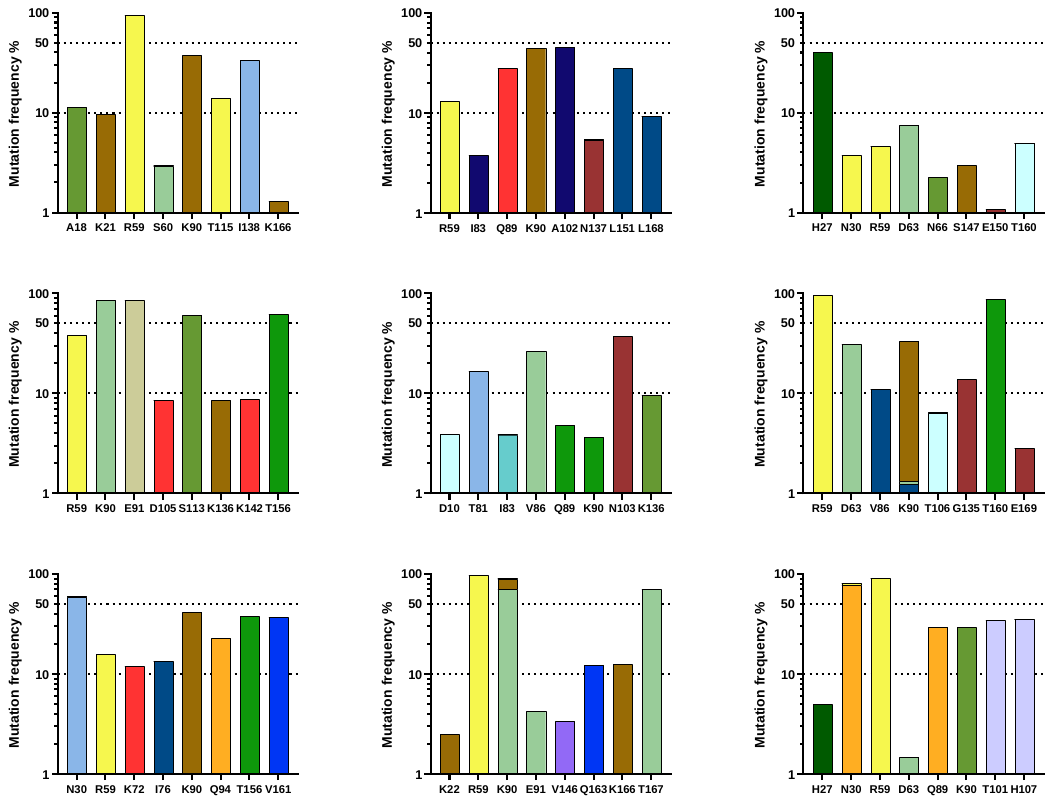

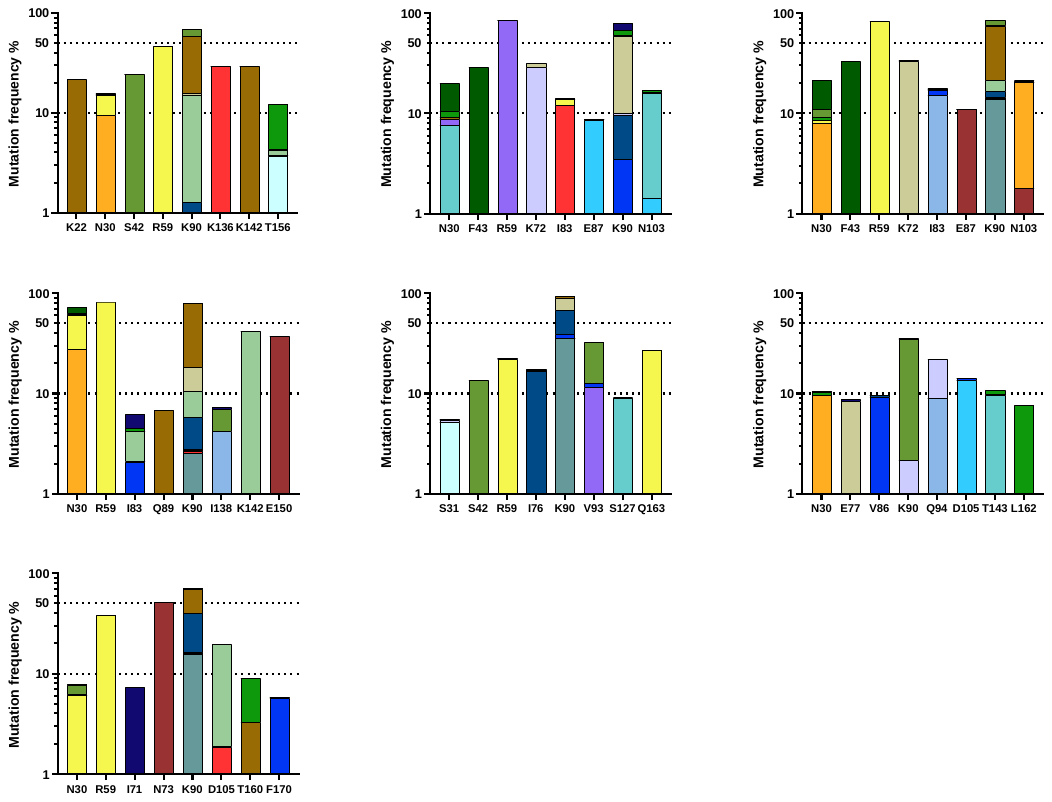

M6/7

M8

M9

M10

M3

M4

M5

Extended Data Figure 5

**b**

**a**

M16

M17

M18

M19

M20

M21

M22

M23

M24

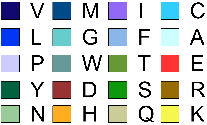

**Extended Data Fig. 5. Diverse and convergent amino-acid mutations in engrafted mice. a** Eight residues with the highest mutation rate across D1D2 from mice (M3 through M10) that were not presented in Fig. 5b, immunized with mRNA-LNP at two-week and four-week intervals. **b** Eight residues with the highest mutation rate from the rest of mice (M16 through M24) immunized with adjuvanted proteins at two-week and four-week intervals, as shown in **Extended Data Figure 2c**. Note that mice M11-M15 were engrafted with low numbers of successfully edited B cells (15,000) like M3-M10; M16-M24, engrafted with 500,000 edited B cells, were immunized with protein antigens. No gp120-binding donor cells were isolated from mice M11-M15, and they were therefore excluded from NGS analysis.

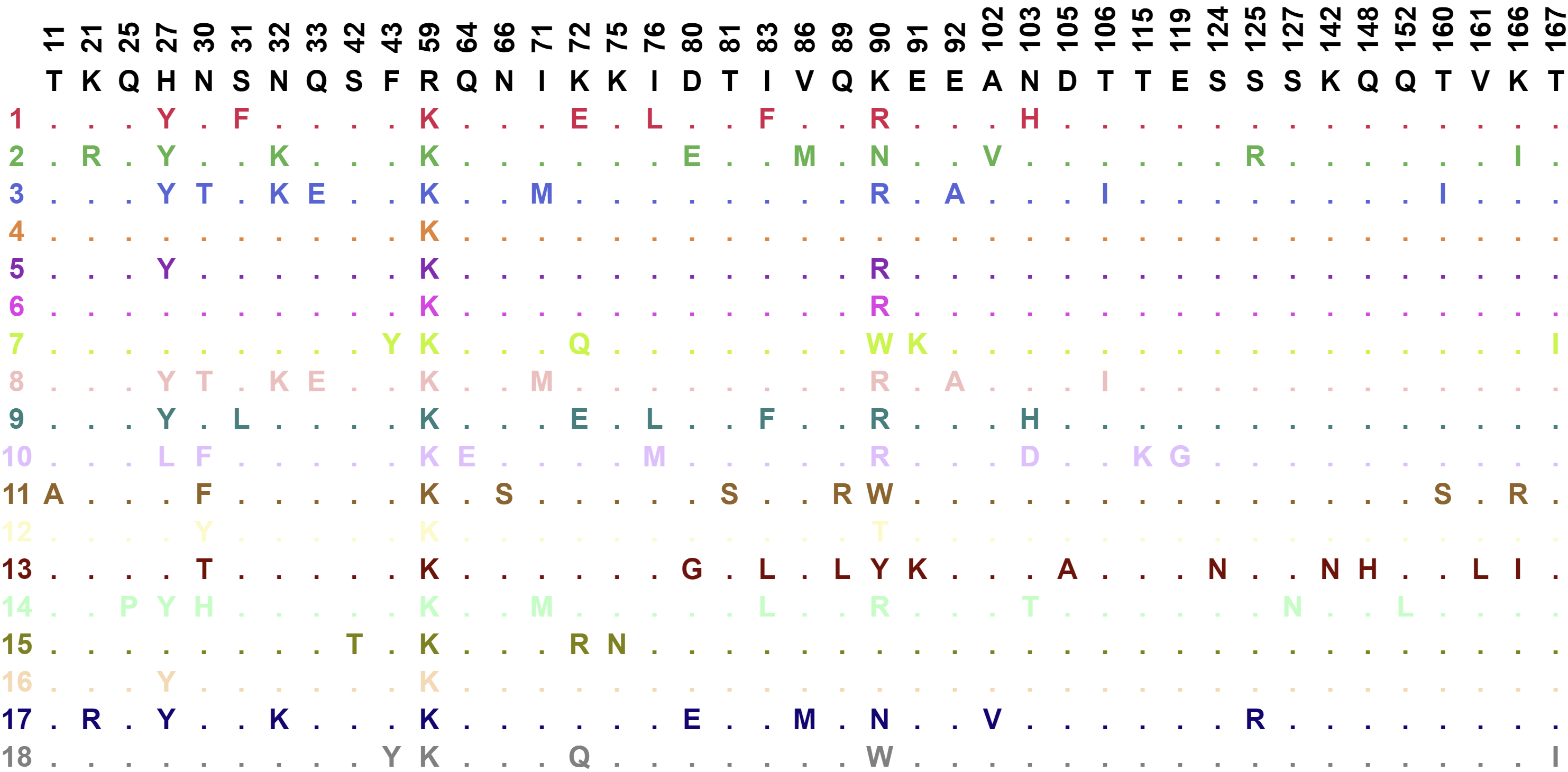

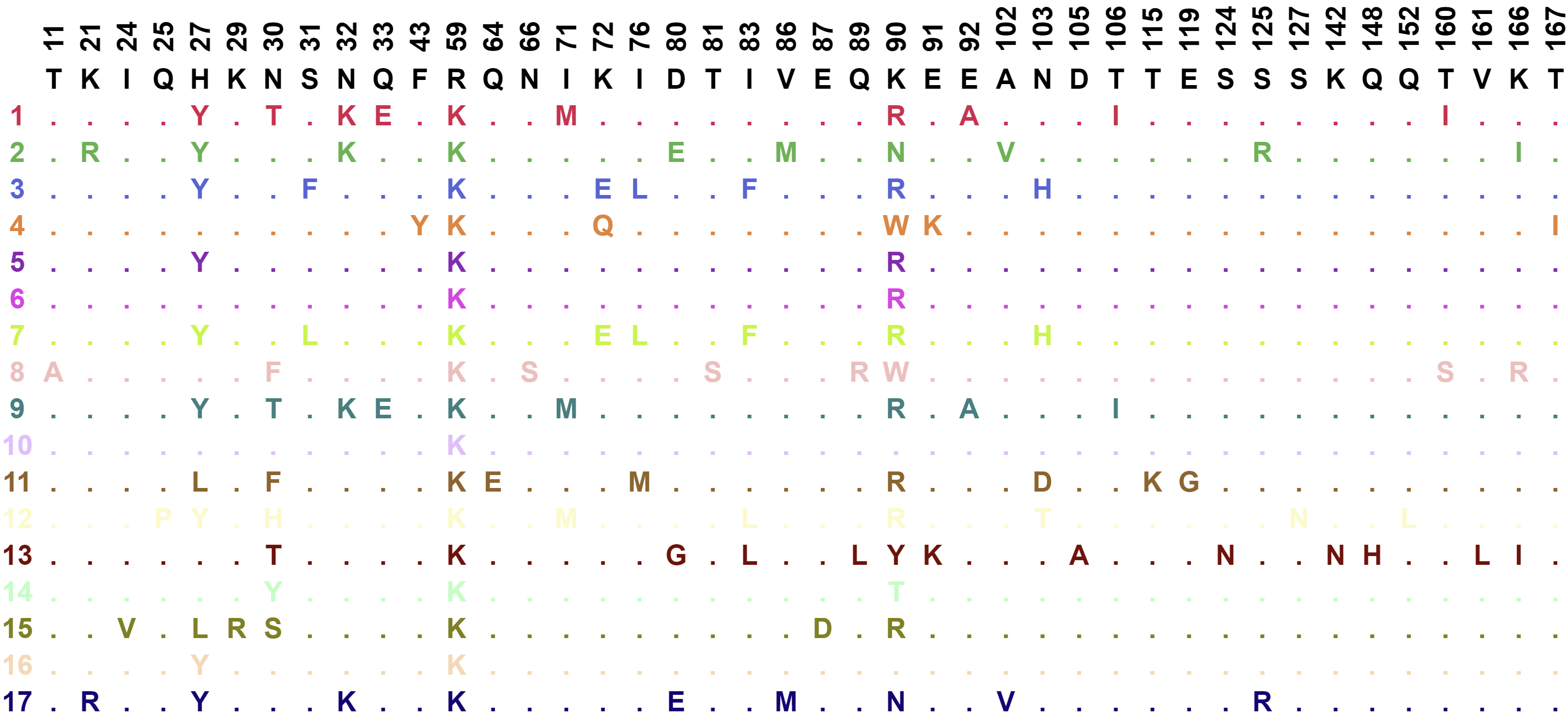

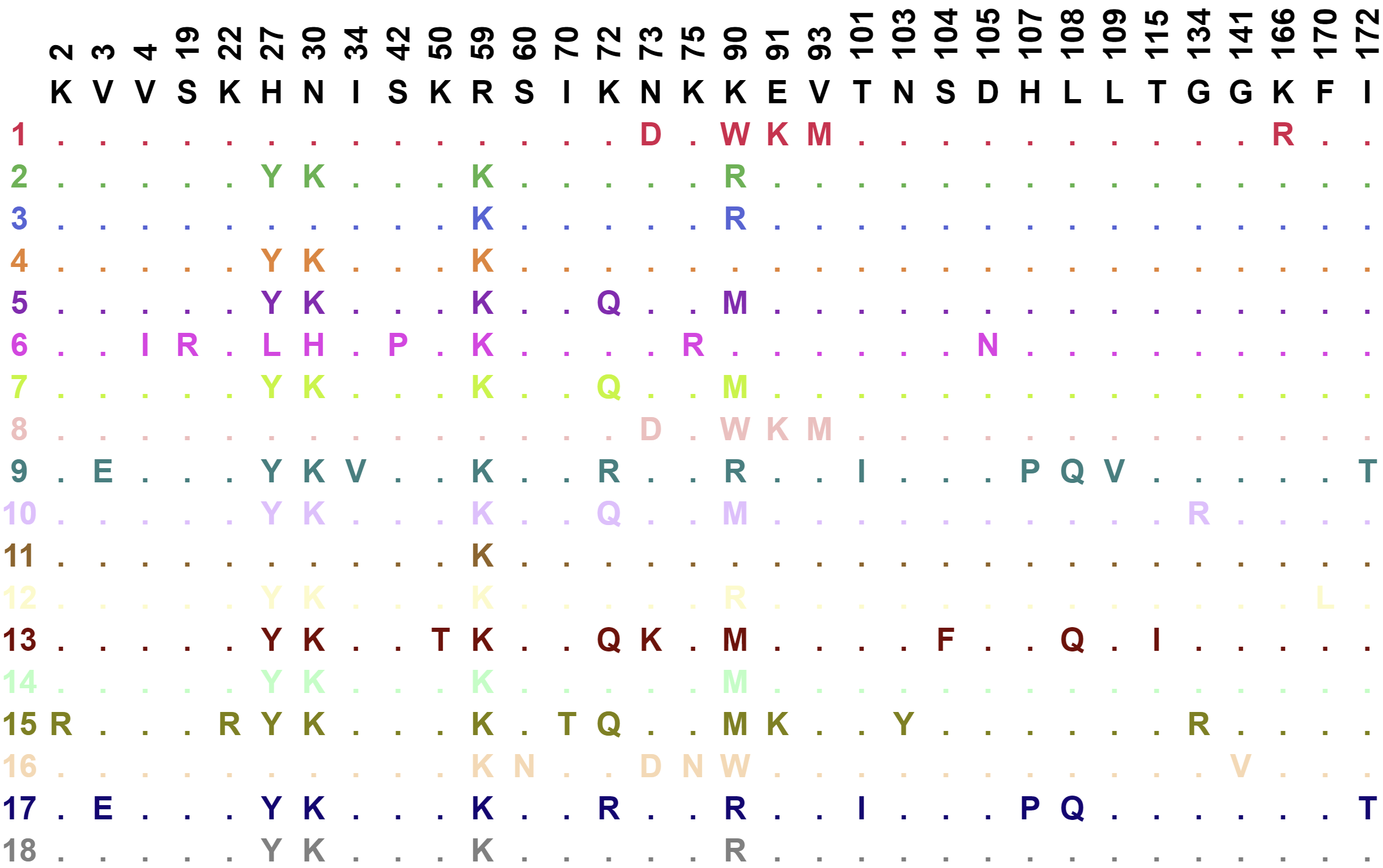

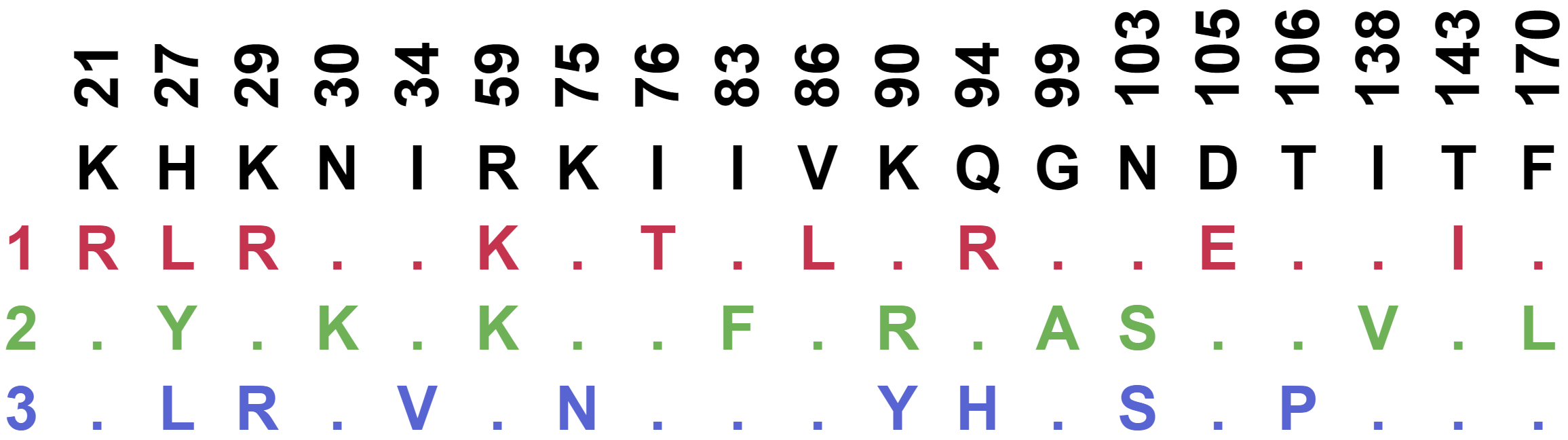

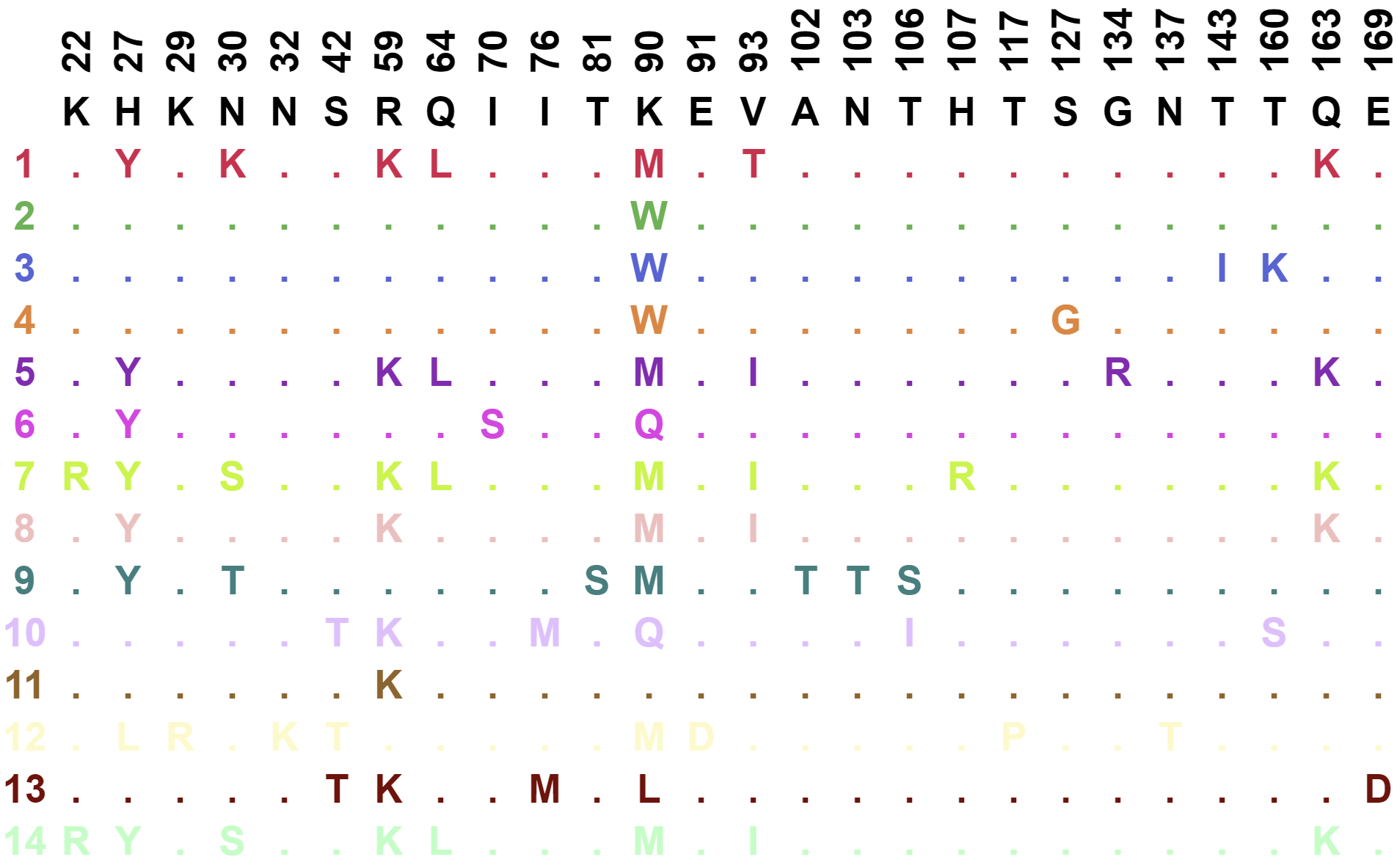

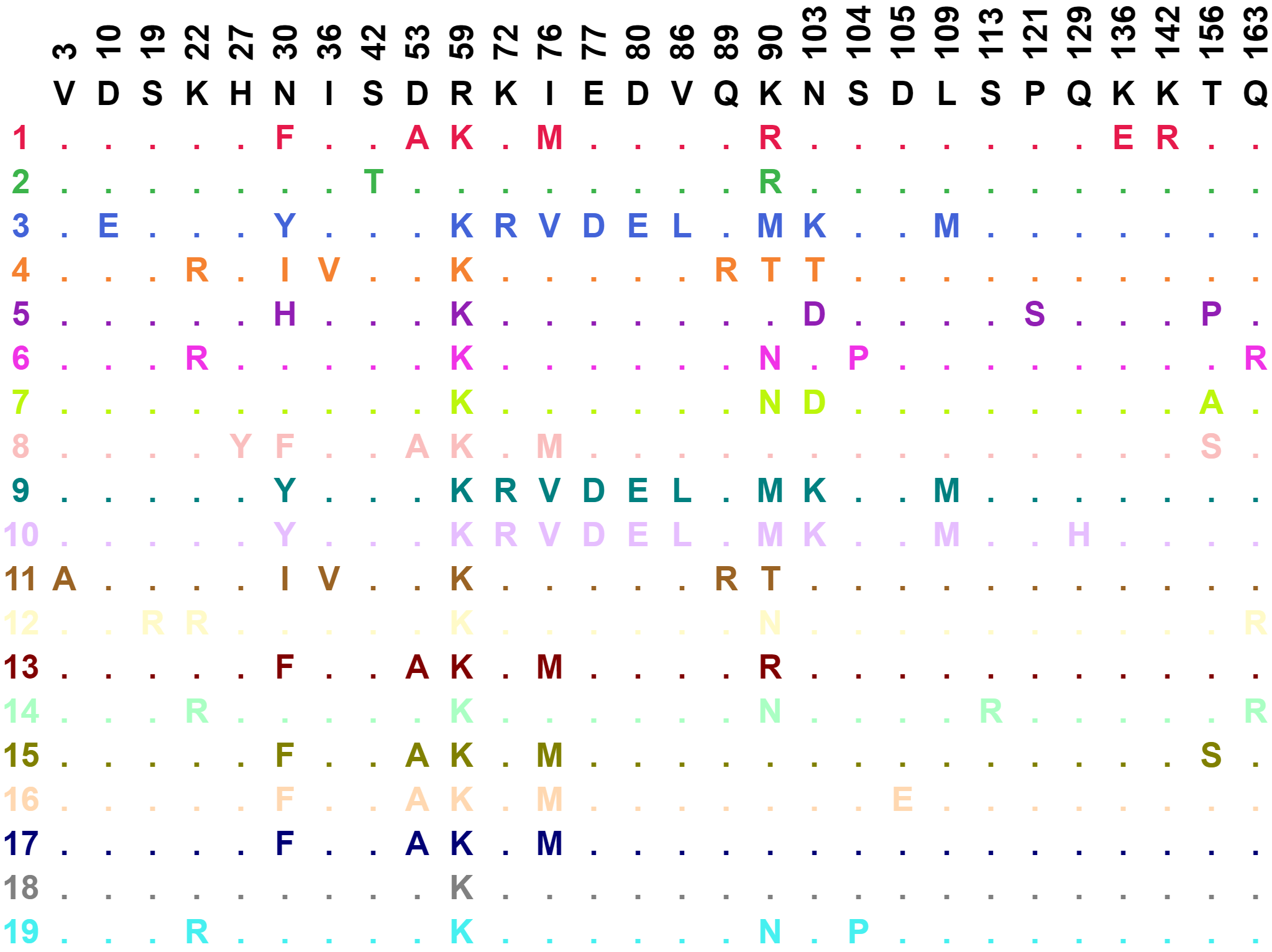

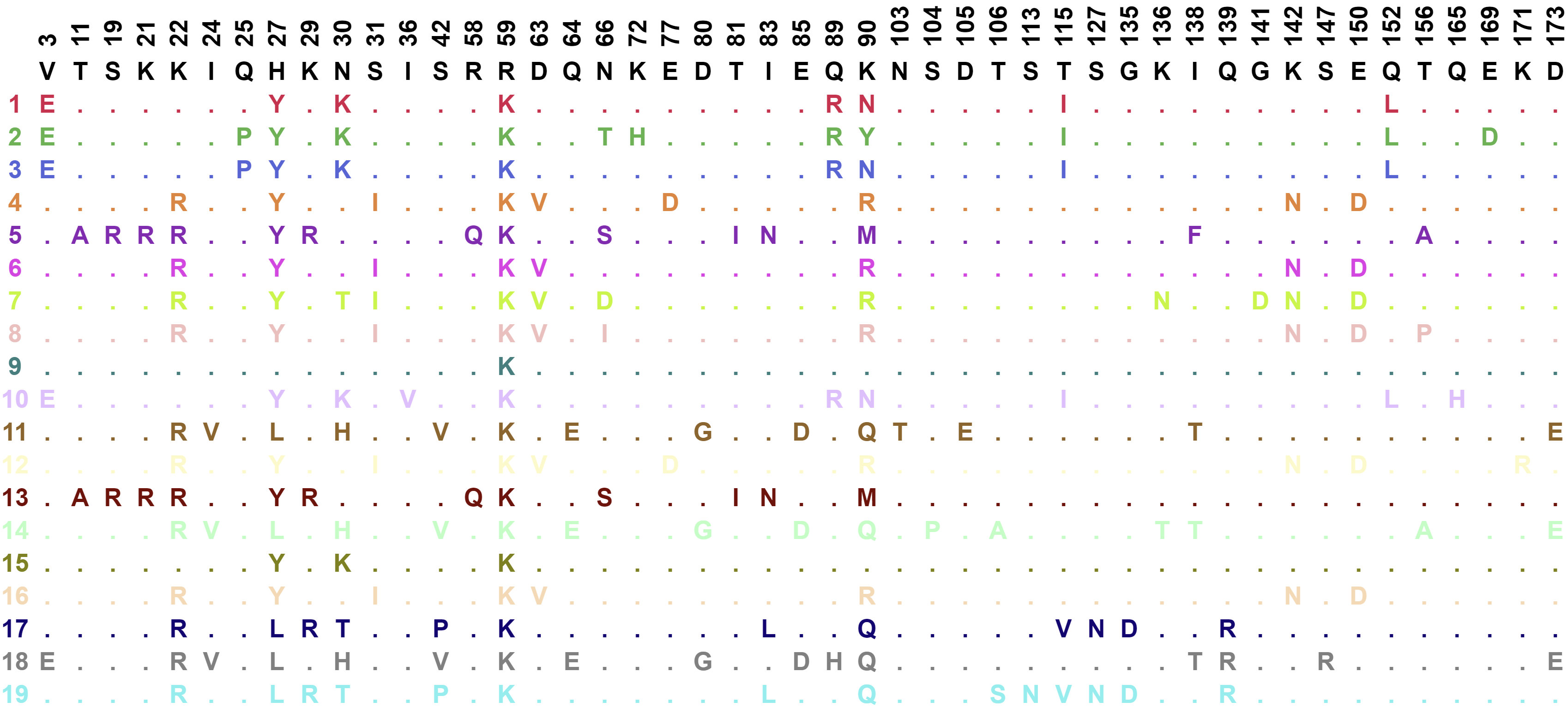

Extended Data Figure 6

M4

M5

M9

M10

M6/7

Unmutated ancestor

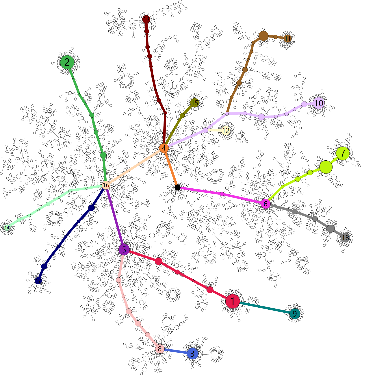

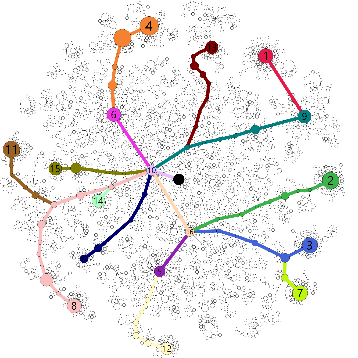

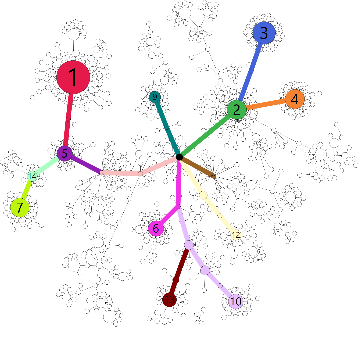

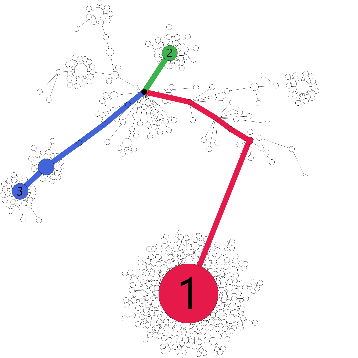

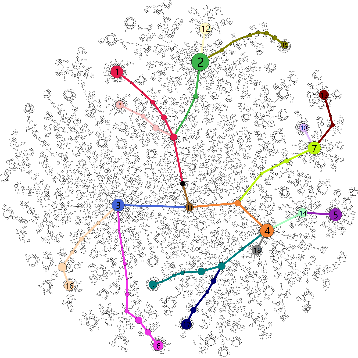

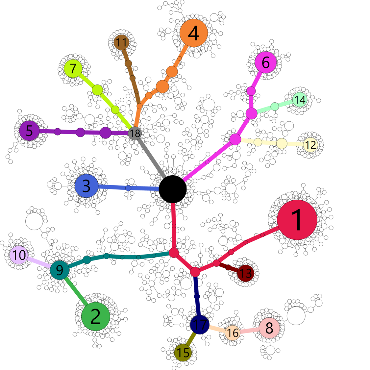

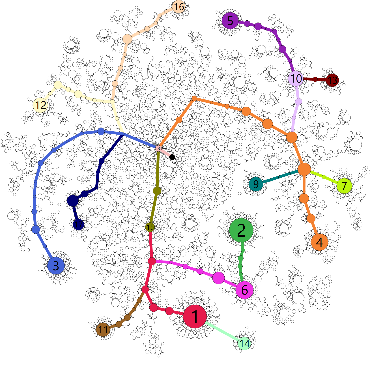

M3

M8

Extended Data Figure 6 (continued)

M17

M18

M20

M22

M19

M16

M21

Unmutated ancestor

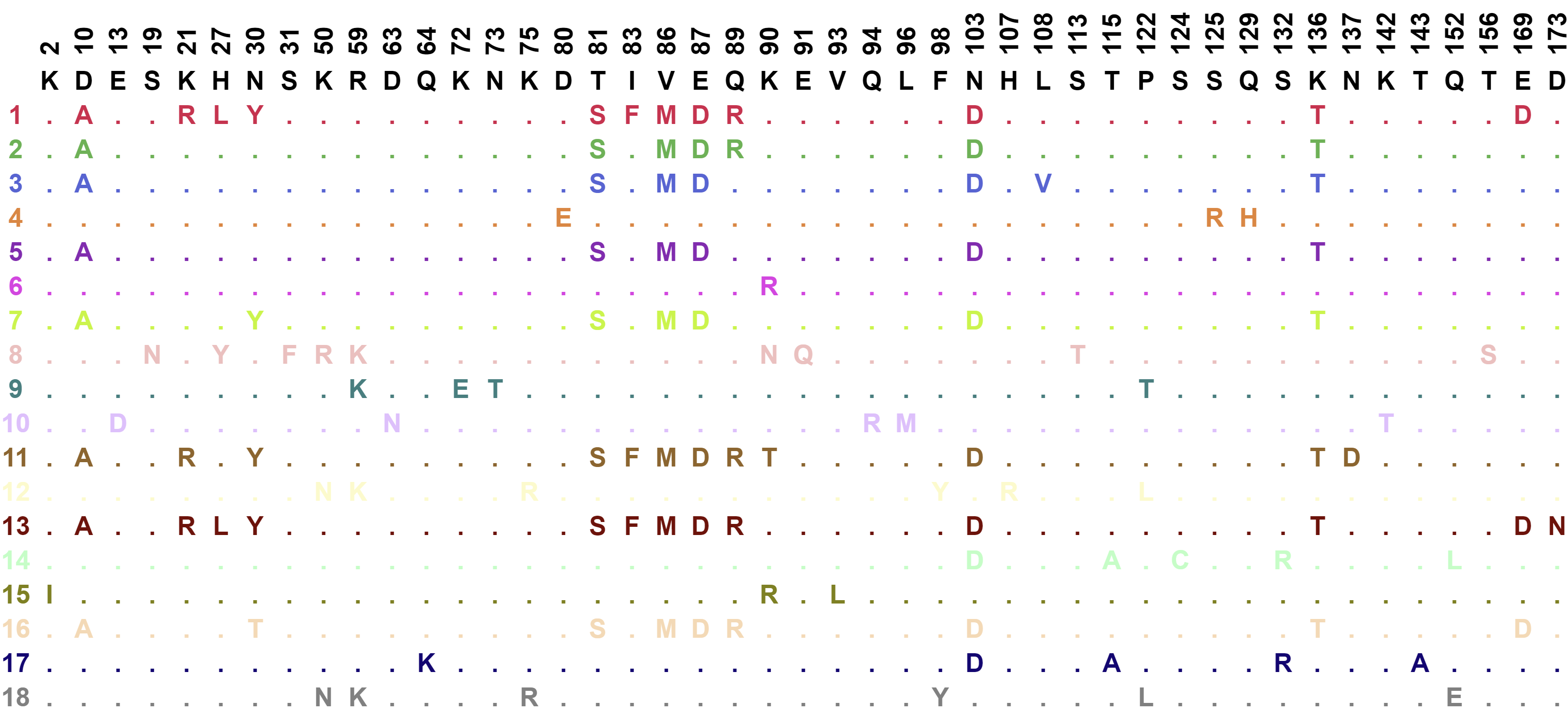

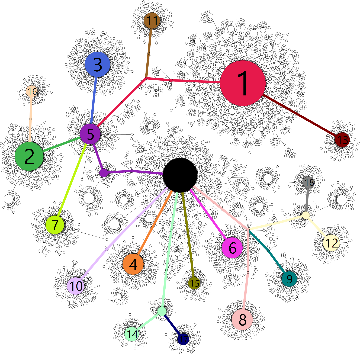

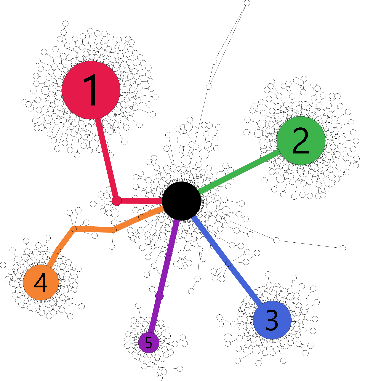

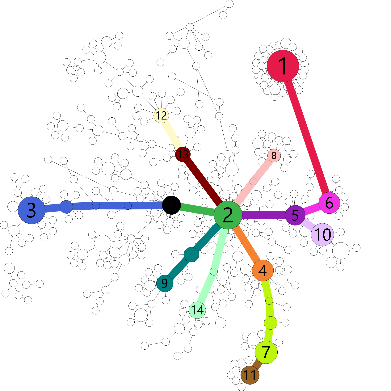

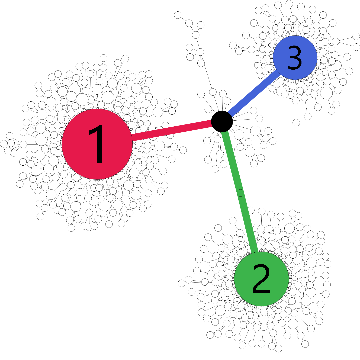

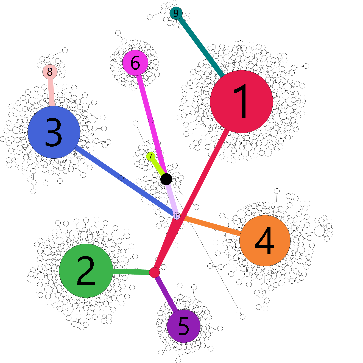

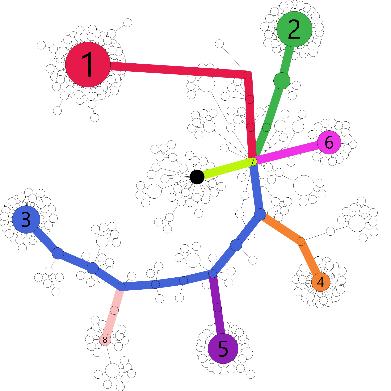

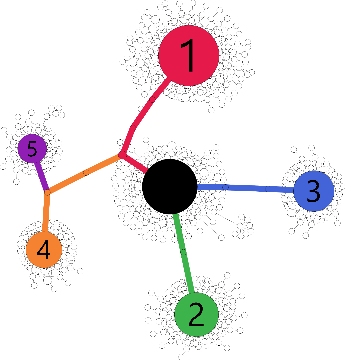

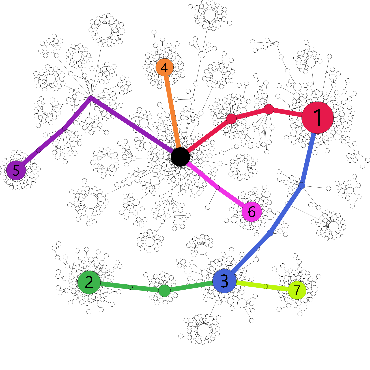

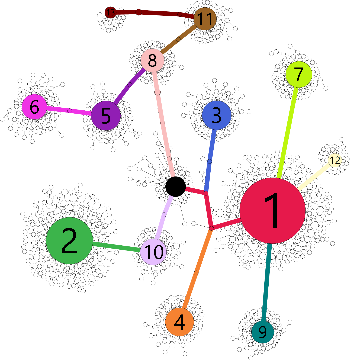

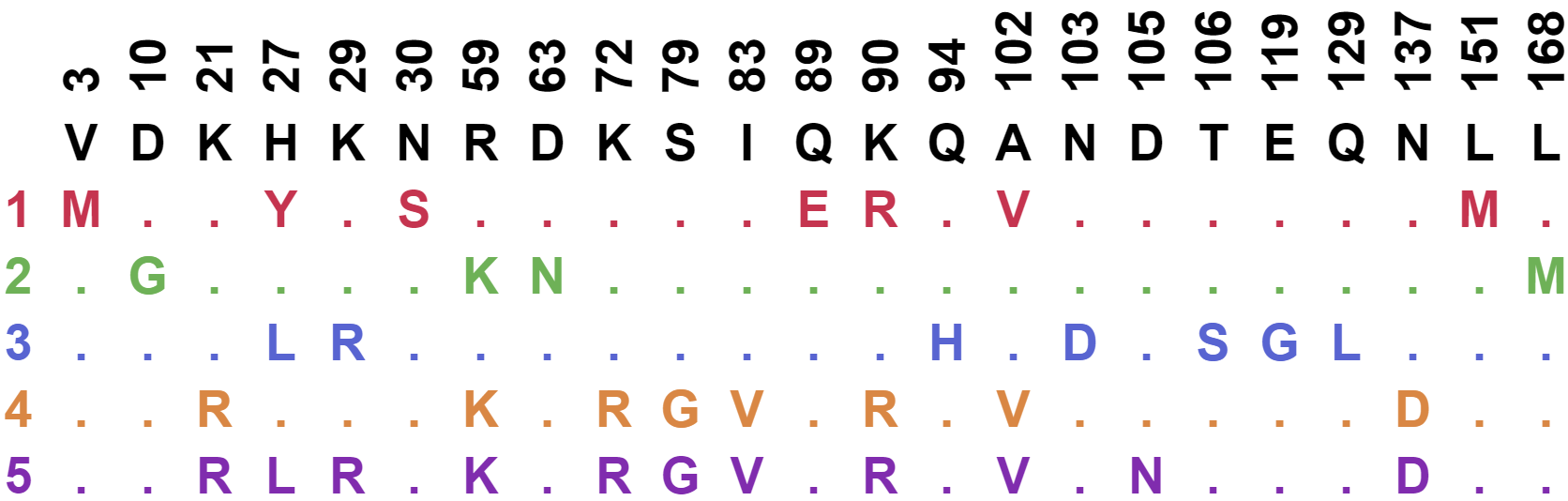

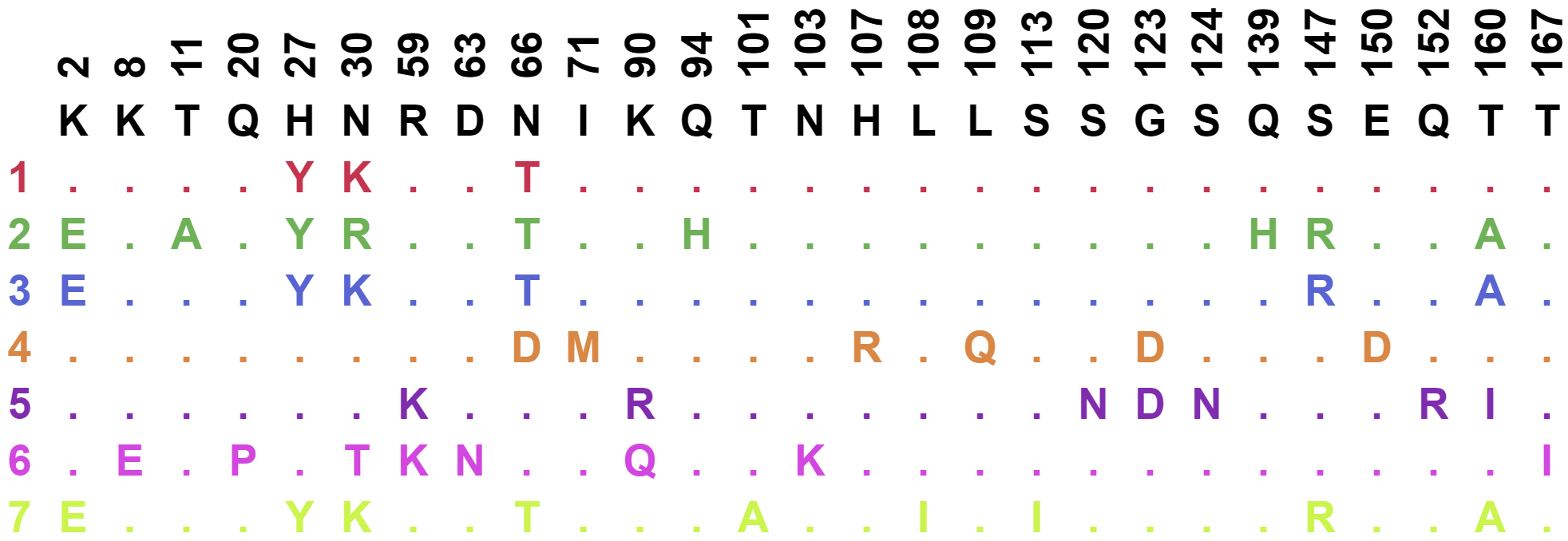

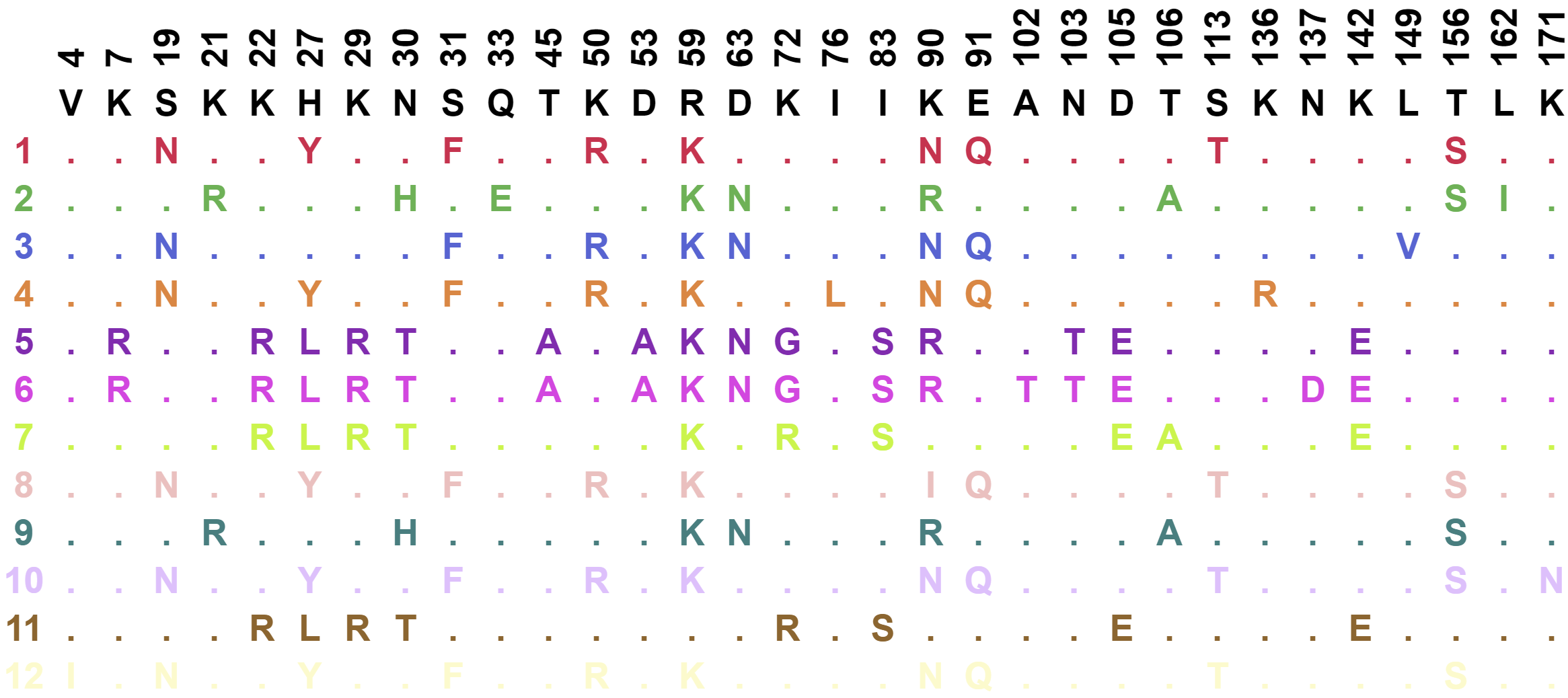

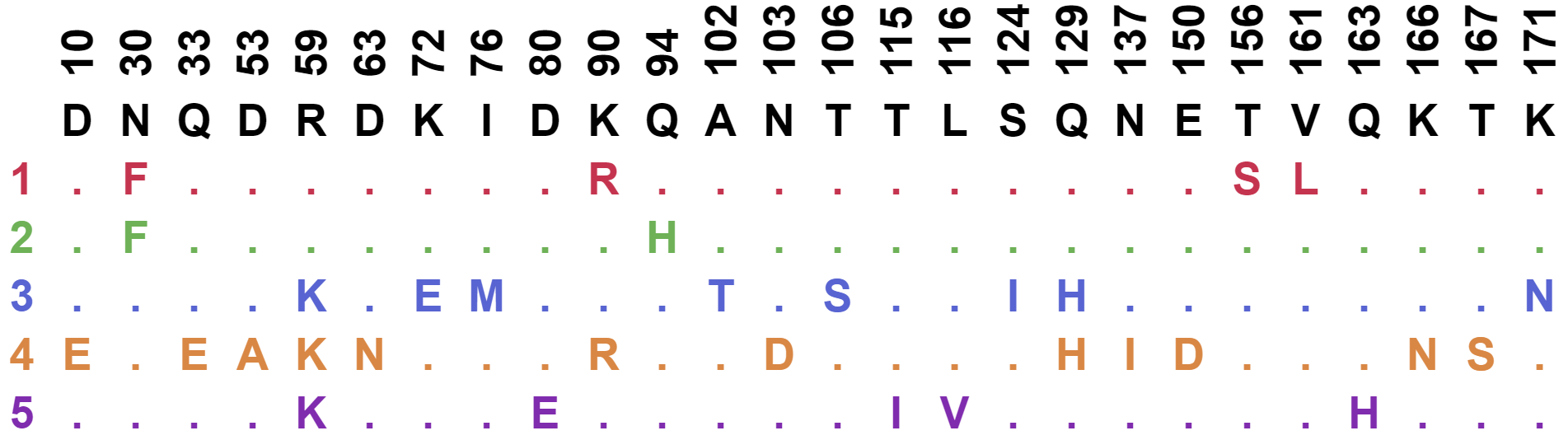

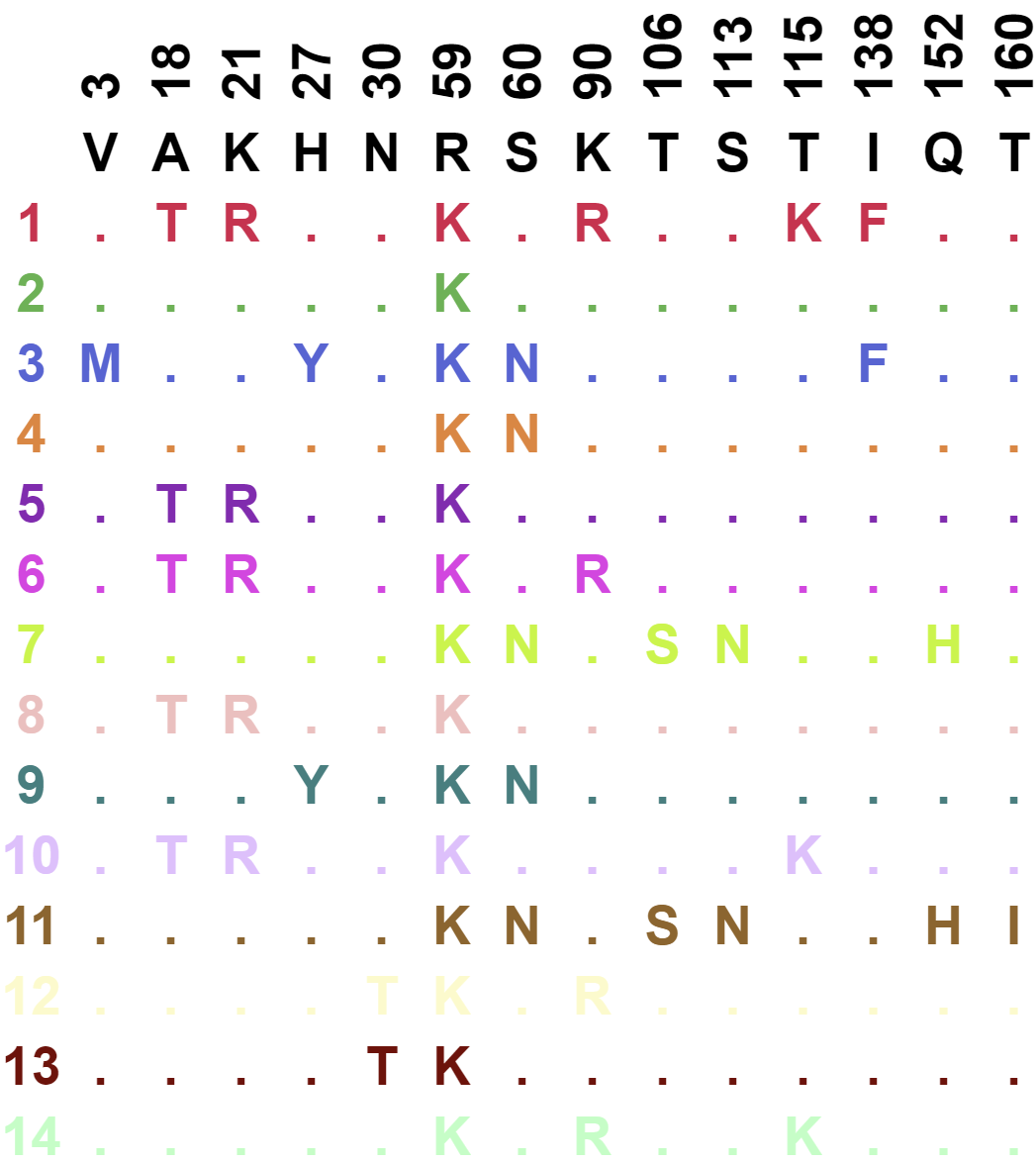

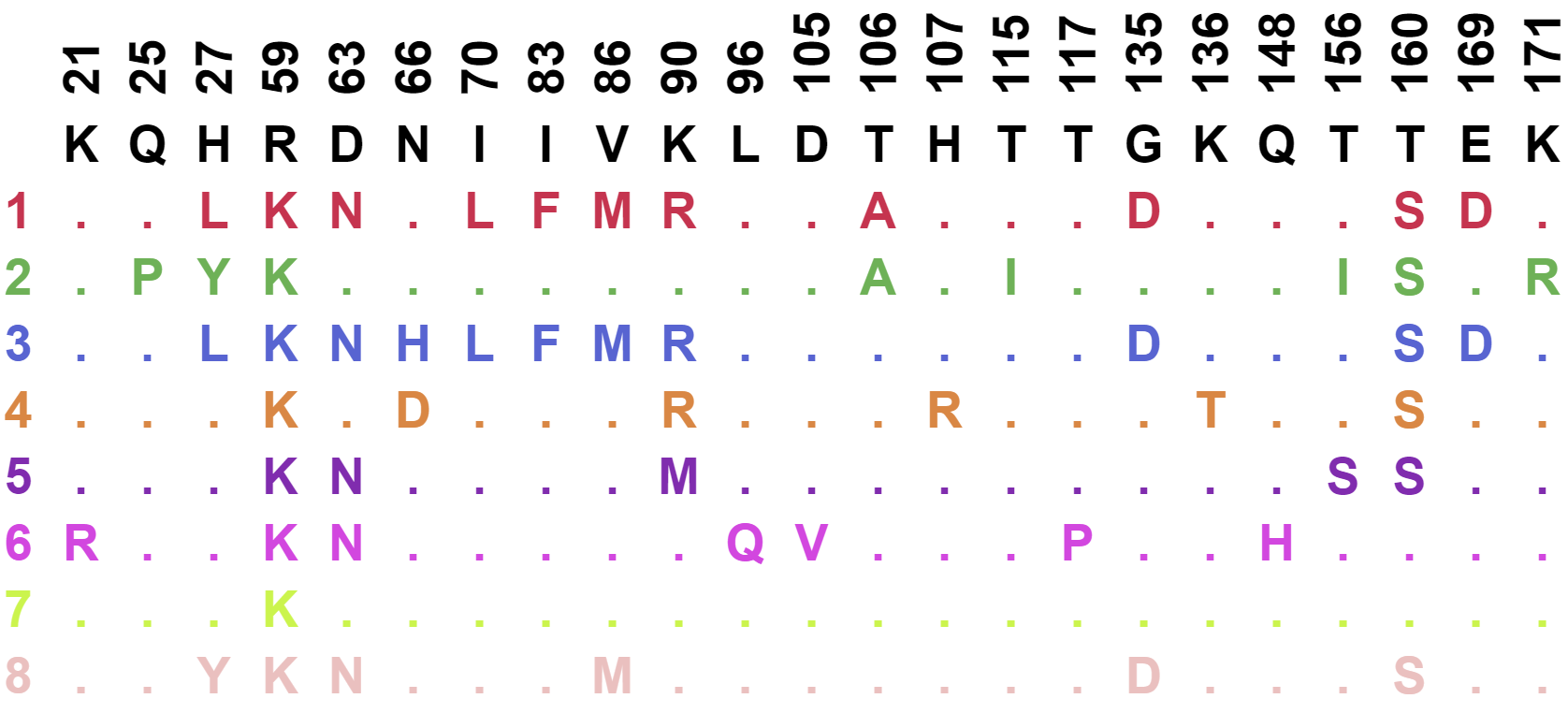

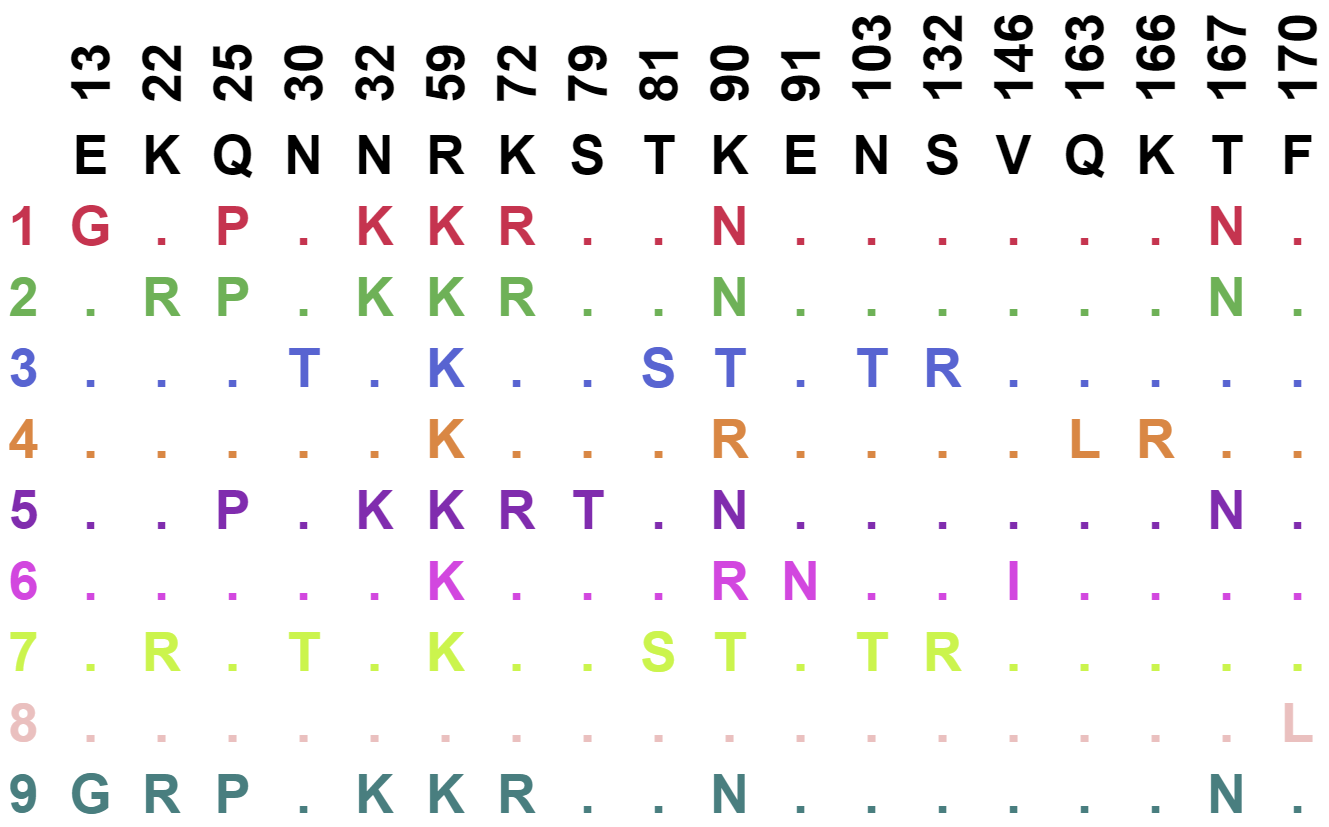

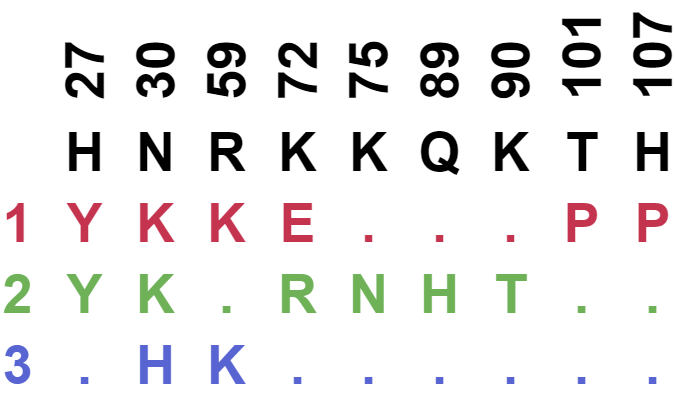

M24

M23

**Extended Data Fig. 6.** **Minimum spanning trees and mutations for mice immunized with mRNA-LNP or** **adjuvanted protein.** Minimum spanning trees of D1D2 sequences, similar to those presented in Fig. 5c, for additional mice immunized by mRNA-LNP every two or four weeks (M3 through M10), or by adjuvanted protein every two weeks (M16 through M24). M6 and M7 were combined into one sample. Each tree presents the inferred lineage and all amino-acid mutations found in each mouse. The central black dot represents the inferred ancestral sequence which corresponds to the input sequence. Each circle indicates a distinct amino-acid sequence. Circle size is proportional to the number distinct nucleotide sequences with the same translation. Colored circles mark the translations encoded by the largest number of distinct sequences, with the rank order indicated by number. Branch length corresponds to evolutionary distance, defined as the number of amino-acid differences.

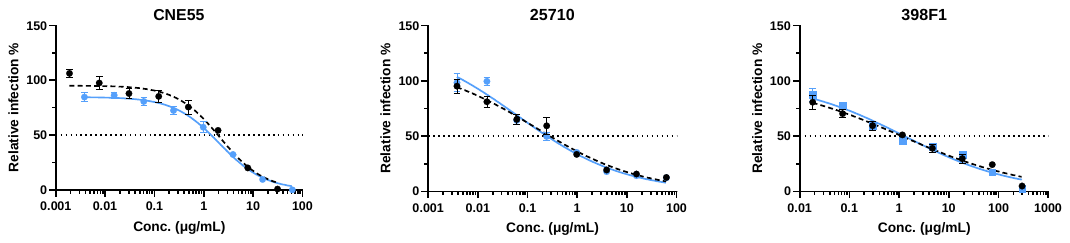

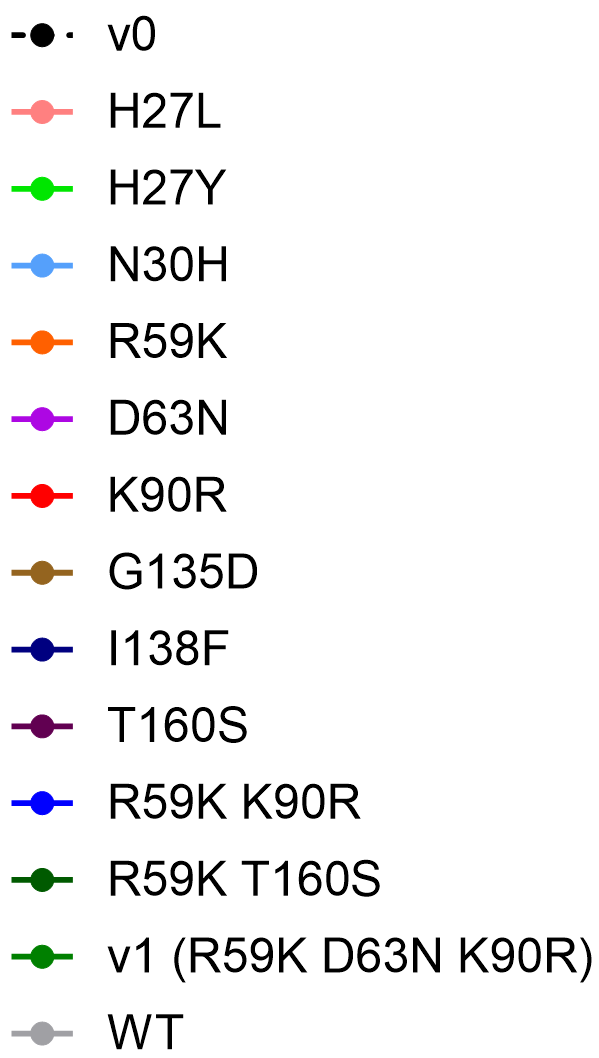

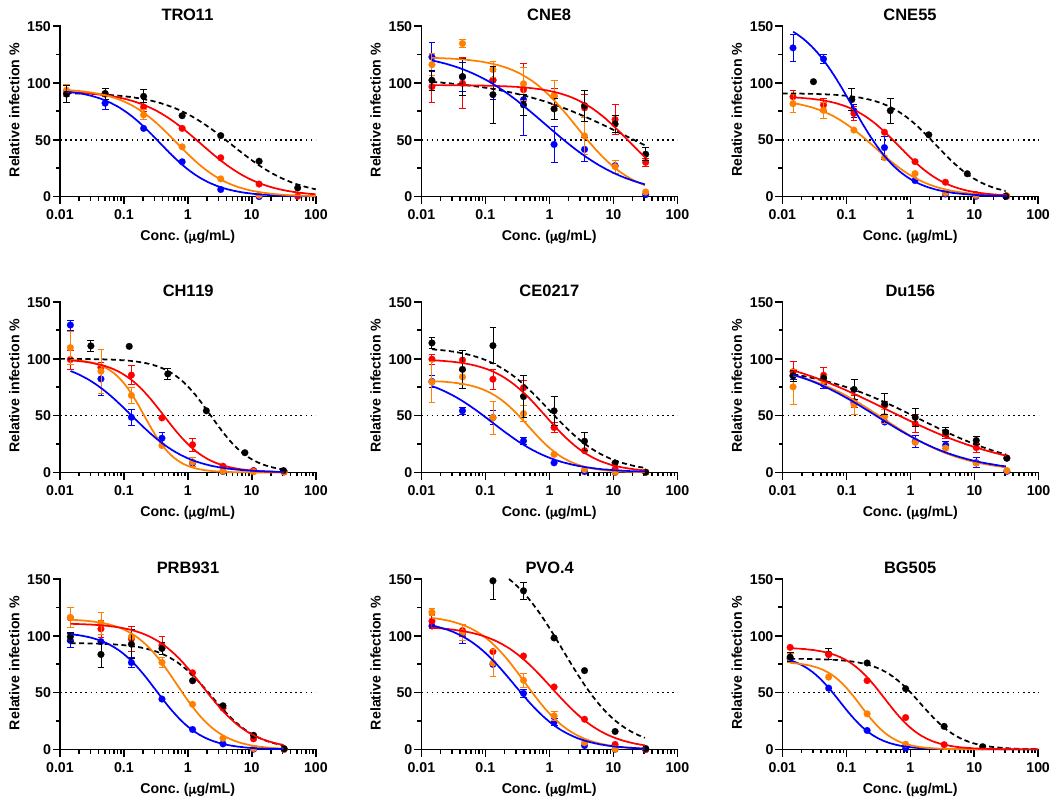

Extended Data Figure 7

**a**

**c**

**b**

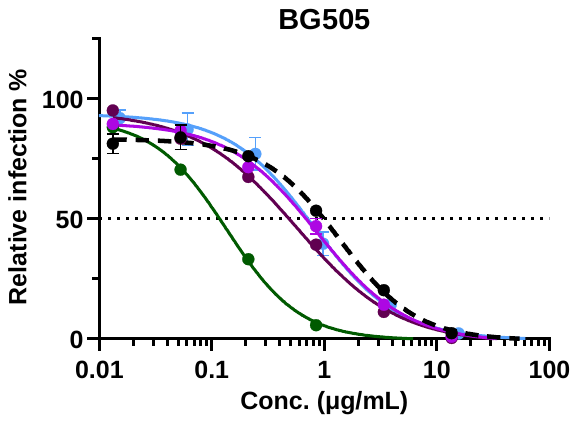

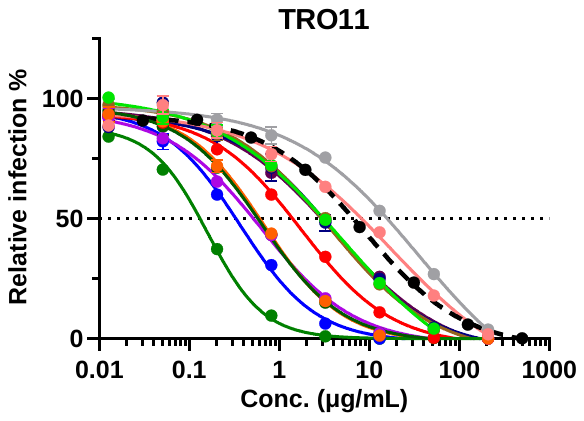

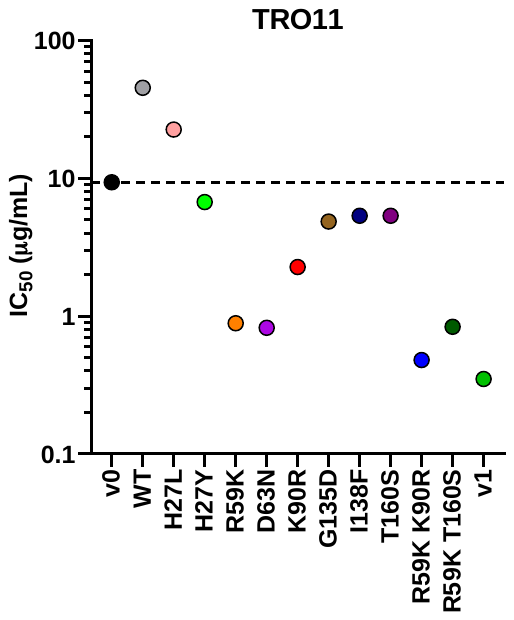

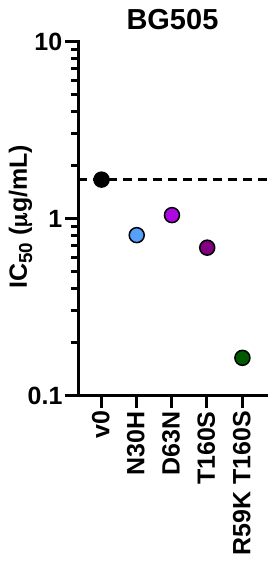

**Extended Data Fig. 7. Neutralization of CD4-Ig variants with single or double mutations against a panel of HIV-1 isolates. a** Neutralization curves of CD4-Ig variants modified with R59K (orange), K90R (red), or combined (blue) against several HIV-1 PV. IC_50_ values are presented in **Fig. 6a**. **b** Neutralization curves of CD4-Ig N30H, a recurring mutation from NGS analysis, against three HIV-1 PVs. **c** Neutralization curves of CD4-Ig variants against BG505 and TRO11 in an initial screening for potent CD4-Ig variants. All curves were fitted with a variable slope four parameters dose response model.

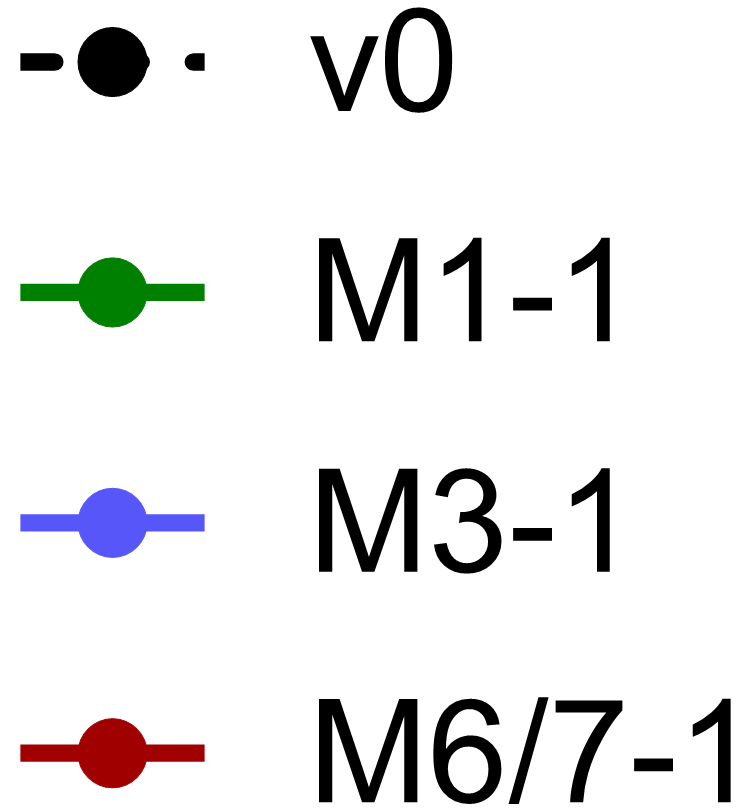

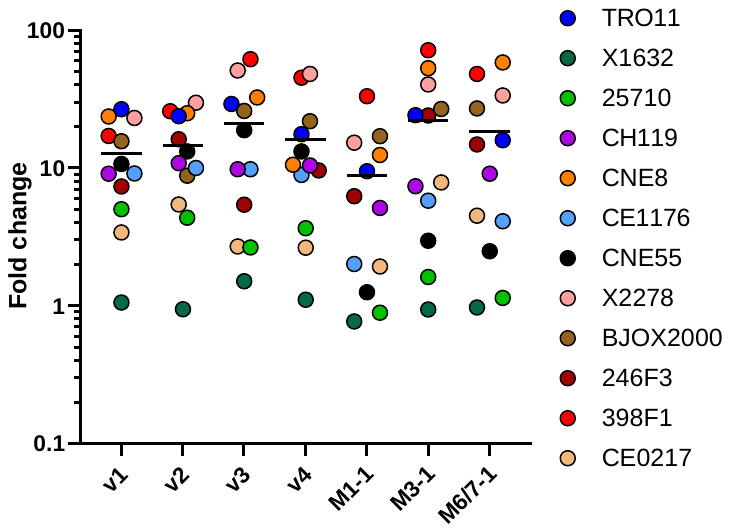

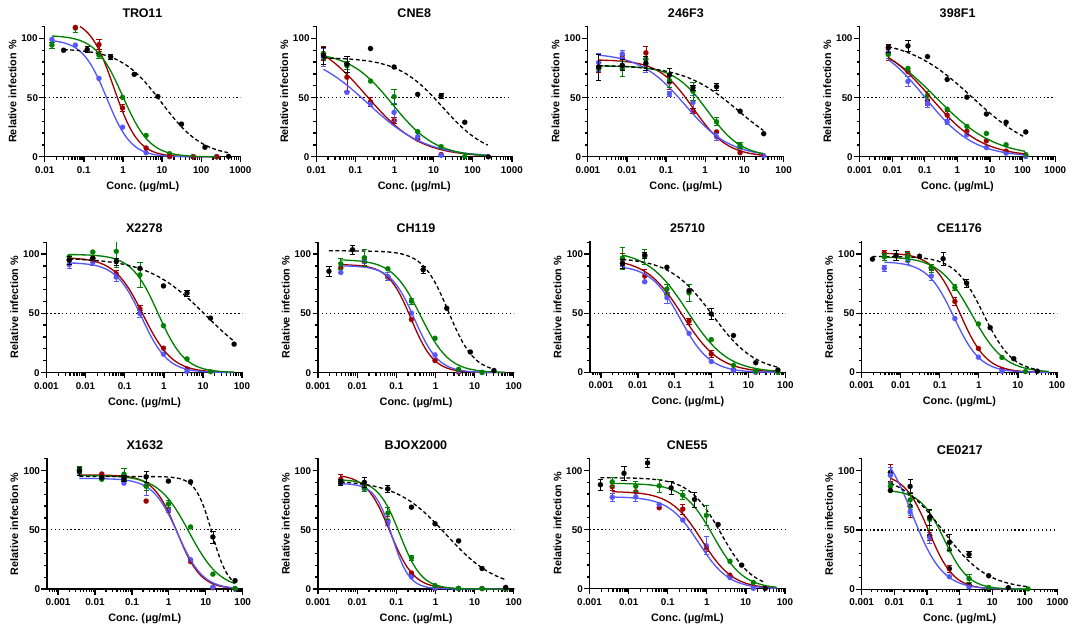

Extended Data Figure 8

**a**

**b**

**Extended Data Fig. 8. Naturally occurring D1D2 variants improved the neutralization potency of CD4-Ig-v0. a** Representative neutralization curves of CD4-Ig-v0 and three naturally emerging variants against a 12-isolate global panel of HIV-1 pseudoviruses. These variants represented the nodes with the largest number of progenies identified in M1, M3, and M6/7 in **Extended Data Fig. 6**. Curves were fitted with a variable slope four parameters dose response model. **b** Fold change of neutralization potency of isolates shown in Fig. 6d relative to CD4-Ig-v0. The center line indicates the geometric mean.

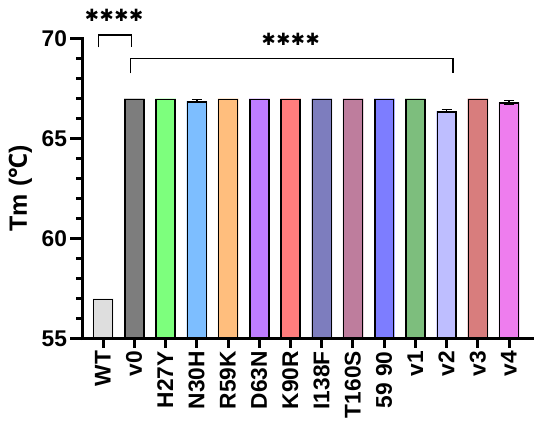

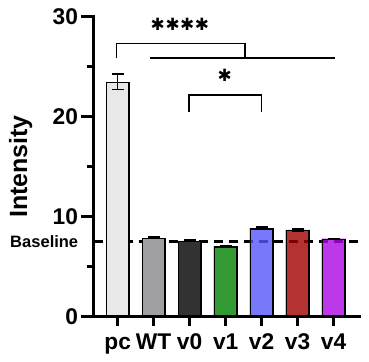

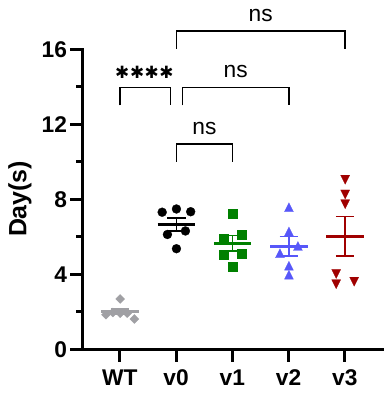

Extended Data Figure 9

**a**

**b**

**c**

**Extended Data Fig. 9. Affinity matured CD4-Ig variants retained bioavailability. a** The polyreactivity of CD4-Ig variants were measured by immunofluorescence assays using HEp-2 cells and 200 μg/ml of each antibody. The autoreactive antibody 2F5 served as a positive control (pc). Baseline (dashed line) was determined as the fluorescence intensity of negative human serum. Each bar is an average of four independent measurements. Error bars indicate SEM. All CD4-Ig variants were significantly less polyreactive than 2F5, as determined by two-way ANOVA with Dunnett’s multiple comparison (*p < 0.05; ****p < 0.0001). **b** Thermostability of CD4-Ig variants. Measured by differential scanning fluorimetry, each bar represents an average of two independent experiments. Significance was determined by two-way ANOVA with Dunnett’s multiple comparison (****p < 0.0001). **c** Pharmacokinetic studies of CD4-Ig variants in immunocompromised hFcRn mice. 8 mg/kg of the indicated CD4-Ig variants was infused intravenously into six nine-week old mice per group. Sera were collected at days 1, 3, 6, 14, 21 and 31. CD4-Ig concentration was measured by ELISA with anti-CD4 antibodies. Half-life was calculated by fitting a one-phase model. Each dot represents the half-life of a CD4-Ig variant in one mouse. Significance was determined two-way ANOVA with Šídák’s multiple comparisons (*p < 0.05; ****p < 0.0001).

**Extended Data Table 1. gRNA, ssDNA enhancer and NGS primers.**

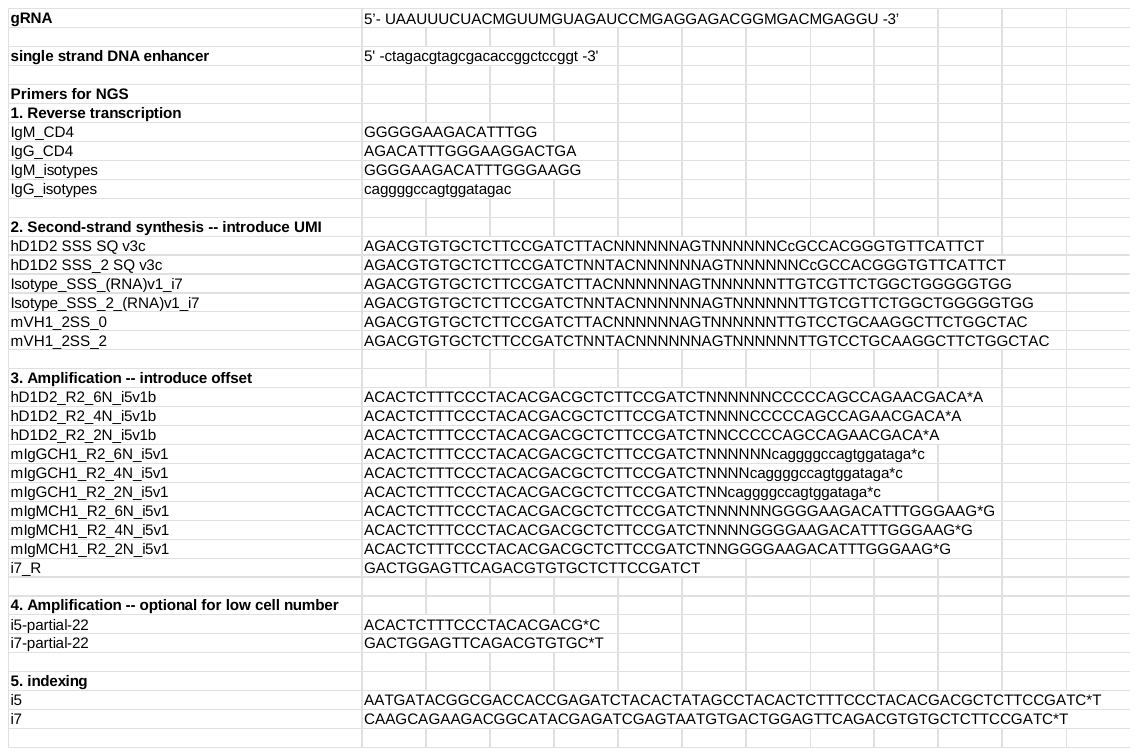
